# Habitat discontinuity and fidelity to migratory routes shape narwhal genetic structure and diversity

**DOI:** 10.64898/2026.07.12.737983

**Authors:** Marie Louis, Mikkel Skovrind, Bárbara Parreira, Alba Rey-Iglesia, Deborah Vicari, Ana Costa, Steven H. Ferguson, Eva Garde, Mads Peter Heide-Jørgensen, Kit M. Kovacs, Christian Lydersen, Lianne Postma, Shyam Gopalakrishnan, Eline D. Lorenzen

## Abstract

Rapid Arctic warming is reshaping marine ecosystems and altering the evolutionary trajectories of ice-associated species. Narwhals (*Monodon monoceros*) are Arctic endemics that are thought to be vulnerable to climate change. We present the first nuclear genomic assessment of narwhals across their distribution to evaluate population structure, demographic history, local adaptation, inbreeding and genetic load. Using genomes from 117 individuals, we identified three populations: Canadian Arctic Archipelago/West Greenland, Northeast Greenland/Svalbard and Southeast Greenland. Demographic reconstructions indicated low effective population size over at least 600,000 years, followed by population growth during the last glacial period. Our results suggest that population structure is maintained by habitat discontinuities and fidelity to migration routes. The latter may promote local adaptation, as genes related to long-term memory were found in regions putatively under selection in Canadian Arctic Archipelago/West Greenland narwhals, which undertake the longest migrations. Genome-wide diversity was uniformly low across populations. Inbreeding levels were inversely related to estimated population sizes. The small and rapidly declining Southeast Greenland population exhibited elevated recent inbreeding. Deleterious mutations were primarily masked in heterozygous genotypes, raising concerns for this population. Together, our results demonstrate that past climate, habitat discontinuities and migration fidelity jointly structure narwhal populations.

## Introduction

Anthropogenic climate change is reconfiguring ecosystems across the planet (IPCC 2023). In the Arctic, temperatures are increasing at two to four times the global average, on regional scales (Rantanen et al. 2022). Rapid bio-physical changes are cascading up through food webs, reshaping the distribution and evolutionary trajectories of species (Wassmann et al. 2011; Hamilton et al. 2017). Understanding the eco-evolutionary processes that facilitate population divergence, range shifts and adaptation to new habitats is thus crucial for informed species conservation in a rapidly changing world (Bernatchez et al. 2024).

For highly mobile species such as marine predators, the environment presents few physical barriers to gene flow (Palumbi 1994). However, in polar regions, sea ice can act as a seasonal or year-round barrier to movement for species that need to surface to breathe, such as cetaceans (Heide-Jørgensen and Laidre 2004). All Arctic endemic marine mammals are seasonally associated with sea ice, which is both a barrier to certain habitats but also in itself critical habitat upon which they rely for foraging and predator protection, as well as breeding and resting in some species (Laidre et al. 2008; Kuletz et al. 2024).

Past climatic shifts, in particular during the most recent glacial period ∼115-11.7 thousand years ago [ka] and subsequent post-glacial warming, have impacted the demographic histories of Arctic marine mammals across their ranges (Foote et al. 2013; Skovrind et al. 2021; Ruiz-Puerta et al. 2023; Westbury et al. 2025). Fine-scale structuring of contemporary populations has been associated with sea ice extent and habitat specialisation in bearded seals *Erignathus barbatus* (McCarthy et al. 2025) and polar bears *Ursus maritimus* (Laidre et al. 2022). In cetaceans, fidelity to migratory routes may act as an additional barrier to gene flow, as has been suggested for beluga whales *Delphinapterus leucas* (O’Corry-Crowe et al. 2018). Additionally, although not yet investigated in Arctic species, behavioural specialisations may also drive local adaptation, as is the case in temperate species such as killer whales *Orcinus orca* (Foote et al. 2016) and bottlenose dolphins *Tursiops truncatus* (Louis et al. 2021b).

Genetic diversity is fundamental for adaptation to environmental change at a local scale. It is expected that accelerating climate change in the Arctic may lead to population declines in endemic species (Pacifici et al. 2020; IPCC 2023; Descamps et al. 2026). This could erode genetic diversity and limit adaptive potential (Willi et al. 2006), and result in the accumulation of genetic load and an elevated risk of inbreeding depression (Keller and Waller 2002; Bozzuto et al. 2019). To fully evaluate the genomic vulnerability of Arctic species to near-future predictions of climate change, spatial baseline data on population structure and diversity across the full range of species are required.

Narwhals are endemic to the Atlantic sector of the Arctic, and are significant to Inuit communities across the region, who rely on the species for food as well as their cultural heritage. As top predators and one of the most sensitive Arctic marine mammal species to climate change, narwhals are bio-indicators of the large-scale changes affecting Arctic marine ecosystems. They have a narrow ecological niche (Laidre et al. 2008) and prefer cold water areas (between 0.5 °C and 1.7°C (Heide-Jørgensen et al. 2020b; Chambault et al. 2020). They show strong site fidelity to migration routes and seasonally used areas, which may reduce their dispersal ability.

Most narwhals undertake seasonal migrations, spending summer in ice-free coastal fjords, bays or sounds (Heide-Jørgensen et al. 2015) and winter in deeper offshore waters with dense drift ice (> 95% sea ice cover), where they forage intensively (Supplementary Fig. S1) (Laidre et al. 2004; Watt et al. 2015). It is generally assumed that narwhals return to the same local summering ground every year with migration distance varying among populations and regions (Dietz and Heide-Jørgensen 1995; Dietz et al. 2008; Westdal et al. 2010; Heide-Jørgensen et al. 2013, 2015; Watt and Ferguson 2015). Narwhals occur year-round in the deep, ice-covered waters between Northeast Greenland and Svalbard (Vacquié-Garcia et al. 2017; Ahonen et al. 2019); the migratory behaviour of these animals and their connectivity with other populations remains unknown (Hobbs et al. 2019).

Narwhals number ∼170,000 individuals globally (Hobbs et al. 2019) but they have extremely low levels of both nuclear and mitochondrial genetic diversity (Palsbøll et al. 1997; Westbury et al. 2019; Louis et al. 2020; de Greef et al. 2024). Deep-time demographic reconstructions based on nuclear genomic data indicate persistently low effective population size (*N_e_*) across at least the past 600,000 years (Westbury et al. 2019; de Greef et al. 2024). Long-term low effective population size can lead to highly deleterious mutations being purged, but mildly deleterious mutations accumulating (Dussex et al. 2023). However, mutational load has never been investigated in narwhals.

The genetic diversity and population structure of narwhals across their distributional area in the Atlantic Arctic remain incompletely characterized. Previous inferences were based on genetic markers with limited resolution, such as partial mitochondrial control region sequences (Palsbøll et al. 1997), microsatellites (Petersen et al. 2011) and mitochondrial genomes (Louis et al. 2020). More recently, nuclear genomic studies have provided evidence of population subdivision within the Canadian Arctic Archipelago (de Greef et al. 2024) or selected localities across the narwhal range (Lopes et al. 2025). A range-wide assessment of the structure and diversity of contemporary narwhal populations encompassing all management stocks based on nuclear data is still lacking.

To fill this knowledge gap, we analysed the nuclear genomes of 117 narwhals sampled across the twelve recognised management stocks in the Atlantic Arctic (Figure 1) (Hobbs et al. 2019). Our study presents the first genomic data from Northeast Greenland and nuclear data from Svalbard. We estimated patterns of differentiation and diversity across the narwhal range, and reconstructed deep-time and more recent population demographic histories to investigate drivers of variation. We tested for signatures of selection to assess local adaptation and estimated inbreeding levels and genetic load to evaluate genomic vulnerability. To elucidate the relative impacts of abiotic (climate, environment) and biotic (ecology, migratory behaviour) factors in driving patterns of population structure and diversity, we integrated our genomic findings with available stable isotope and tagging data from the literature.

**Figure 1.**
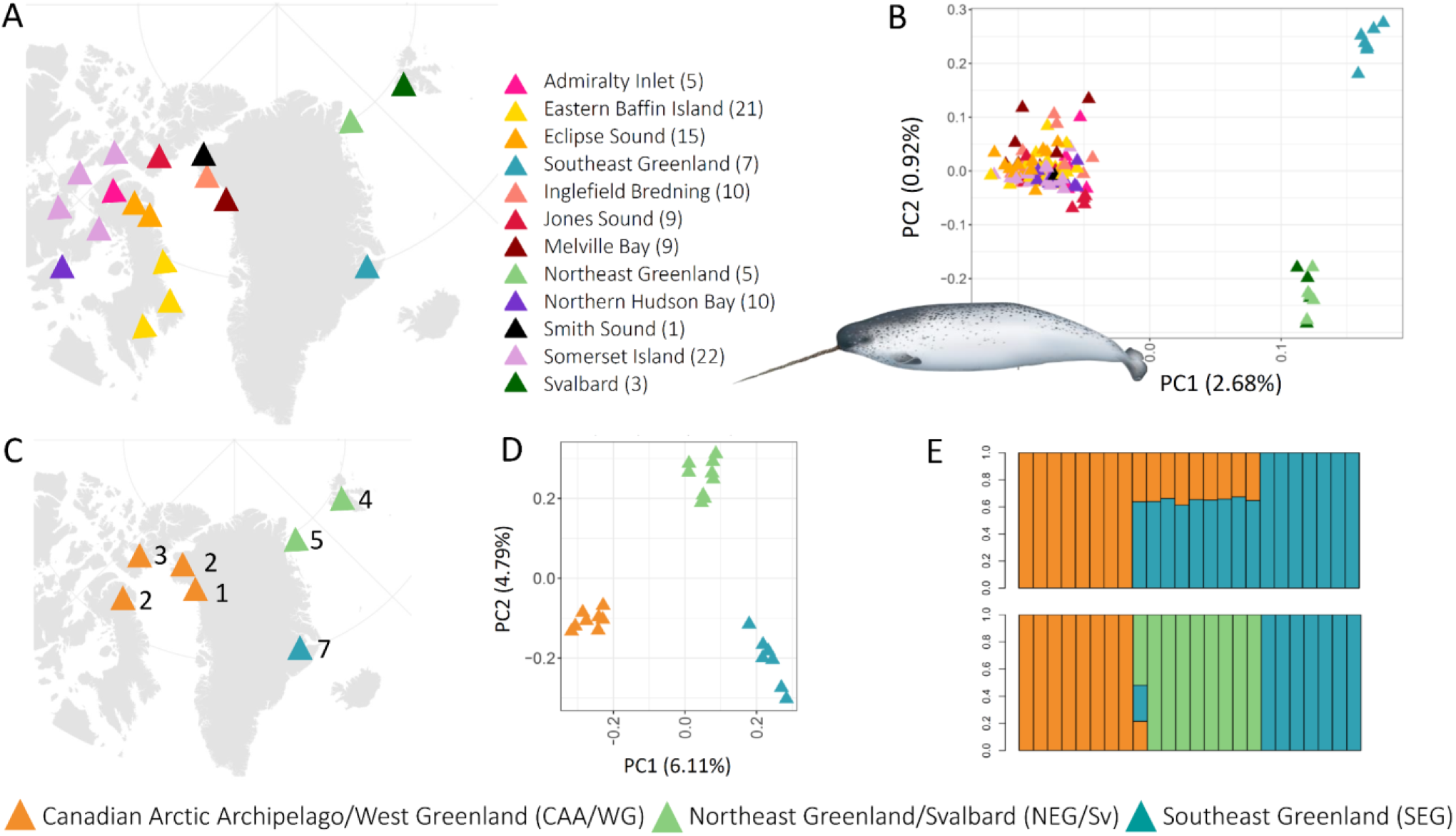
Population structure of narwhals across their range. A) Sample localities of the range-wide dataset (n = 117; mean 2.73× coverage). B) PCA of the range-wide data using PCAngsd based on 486,709 SNPs, showing the proportion of genetic variance captured by the two first principal components. C) Sample localities of the population-level dataset (n = 24; mean 9.63× coverage). D) PCA using PCAngsd and (E) ancestry proportions inferred by NGSadmix for *K*=2 (upper) and *K*=3 (lower) for the population-level data based on 274,852 SNPs; *K*=3 was identified as the best-fitting *K* to the admixture model. Narwhal illustration by Uko Gorter.

## Results

Our study is based on two genomic datasets (Fig. 1A,C, see methods section). The range-wide dataset included 117 narwhal individuals sequenced at 0.15-11.99× (mean = 2.73×, mode = 0.3×) coverage (Supplementary Fig. S2A, Supplementary Table S1). The dataset included 486,709 unlinked SNPs and did not include closely related narwhals (r<0.1, Supplementary Fig. S3). This dataset was used for range-wide inference of population structure.

The population-level dataset included nuclear genomic data from 24 narwhals sampled from the three putative populations identified in the preliminary analyses of the range-wide dataset (Fig. 1C). The population-level dataset included eight individuals from CAA/WG, nine from NEG/Sv and seven from SEG, sequenced at 7.18-12.75× (mean = 9.63×, mode = 9×) coverage and included 274,852 unlinked SNPs (Supplementary Fig. S2B, Supplementary Table S1). We used this dataset for demographic reconstruction and modelling, estimation of levels of inbreeding and mutation load and for selection analyses.

### Population structure

Based on Principal Component Analysis (PCA) generated using PCAngsd (Meisner and Albrechtsen 2018) and admixture analyses in NGSAdmix (Skotte et al. 2013) of the range-wide dataset, we identified three genetically and geographically distinct populations: 1) Canadian Arctic Archipelago/West Greenland (CAA/WG, n=101); 2) Northeast Greenland/Svalbard (NEG/Sv, n= 9); and 3) Southeast Greenland (SEG, n= 7) (Fig.1A-B, Supplementary Fig. S4-5). PC1 (2.68%) separated narwhals in CAA/WG from the localities east of Greenland. PC2 (0.92%) separated NEG/Sv from SEG narwhals.

The best estimate for the number of populations (*K*) was determined using evalAdmix ^(^Garcia-Erill and Albrechtsen 2020^)^ and was found to be *K*=3 (Supplementary Fig. S6A). *F*_ST_ estimates confirmed that the level of genetic differentiation among the three populations is higher than levels identified among the eight stocks present within the largest population CAA/WG (we had only 1 sample for the Smith Sound stock, therefore it was not included in this analysis, Supplementary Fig. S7).

To test for further population structure within CAA/WG, we ran a PCA of only the individuals sampled from localities within this region (n=101); however, no further population subdivisions were identified (Supplementary Fig. S8). However, narwhals sampled in Melville Bay clustered towards one end of PC1. Six out of ten Melville Bay samples were taken the same day, and included two pairs of related animals, from which we removed one individual from each pair for further analyses (Supplementary Fig. S3).

The PCA of the population-level dataset underscored the genetic differentiation among the previously identified populations (Fig. 1D). PC1 (6.11%) indicated that CAA/WG narwhals were most distinct from narwhals in SEG (*F*_ST_ = 0.054), with narwhals in NEG/Sv being intermediate (*F*_ST_ CAA/WG-NEG/Sv = 0.043; SEG-NEG/Sv = 0.040). PC2 (4.79%) separated NEG/Sv narwhals from the two other populations. Similar to the results of the range-wide dataset, *K*=3 was identified as the best fit (Fig. 1E, Supplementary Fig. S6B, S9).

### Evolutionary relationships

To explore the evolutionary relationships between the three identified narwhal populations, we used Treemix, admixture graphs (qpBrute and Admixture Bayes), *D*-statistics, and *f3*-statistics on the population-level dataset (Patterson et al. 2012; Pickrell and Pritchard 2012; Ní Leathlobhair et al. 2018; Liu et al. 2019; Nielsen et al. 2023).

Both Treemix and admixture graphs showed low levels of divergence among the three populations since their separation, which is indicated by short internal branch lengths in Treemix and low drift values on each internal branch of the admixture graphs (Fig. 2A, Supplementary Fig. S11-14). These results imply low levels of genetic divergence among populations, which is supported by the 2D-SFS that show highly correlated genomic variation among them (Supplementary Fig. S16). Thus, either the three populations diverged relatively recently, or they radiated from each other within a short amount of time.

**Figure 2.**
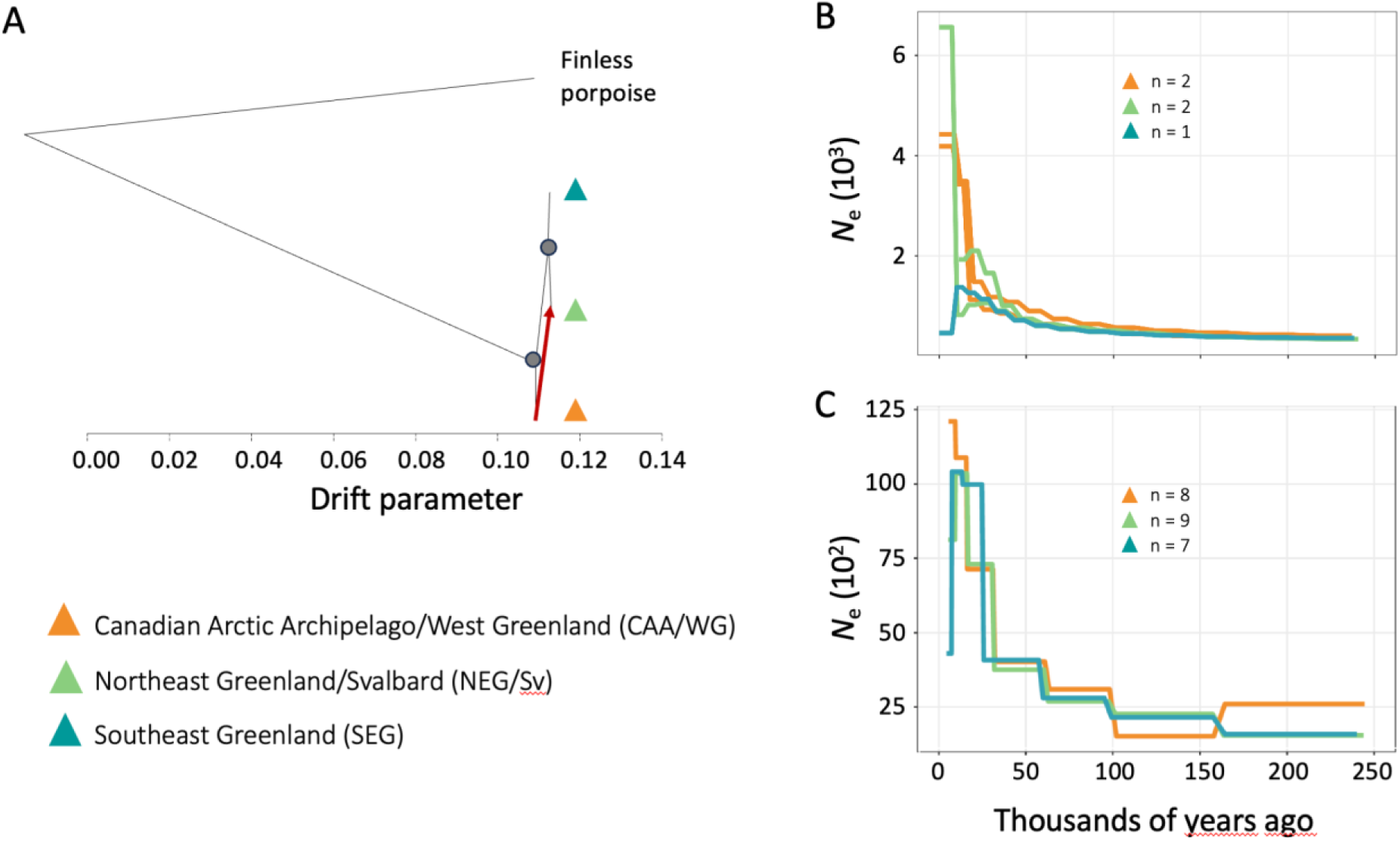
Demographic history of the three narwhal populations. A) TreeMix consensus tree; nodes have bootstrap values > 90%, showing the relationships among the three populations as a bifurcating maximum-likelihood tree with one migration edge (M=1, in red), inferred as the best topology. Branch lengths on the horizontal axis represent the level of genetic drift that has occurred along each branch. The tree is rooted using the finless porpoise (*Neophocaena phocaenoides*). B) Demographic reconstruction based on individual genomes using pairwise sequentially Markovian coalescent (PSMC) with a mutation rate of 1.56e-08 per generation (Westbury et al. 2019) and a generation time of 30 years (Garde et al. 2007). C) Population-based demographic reconstruction of each narwhal population using SMC++.

Consistent with the PCA and NGSadmix results, Treemix analyses, admixture graphs, and D-statistics all indicated that the initial split occurred between narwhals west and east of Greenland, followed by a second split separating the two populations east of Greenland (Supplementary Fig. S17). Using *D*-statistics, we found support for the tree (NEG/Sv,SEG;CAA/WG,out) with a *Z*-score of 0.18. This finding did not suggest admixture among populations, although Treemix indicated a migration edge from CAA/WG into NEG/Sv, supported by *f3*-statistics, with a significantly negative *Z*-score of -4.62 for (NEG/Sv;CAA/WG,SEG) (Fig. 2A, Supplementary Fig. S11).

### Demographic reconstruction

To reconstruct deep-time and more recent demographic history of the three narwhal populations, we used six complementary approaches. Based on the five individual genomes with coverage between 13 and 60× (Supplementary Table S1), we inferred long-term effective population sizes with the pairwise sequentially Markovian coalescent (PSMC) (Li and Durbin 2011). We found similar deep-time demographic histories across the genomes, with long-term low effective population size (*N_e_*) across the past 600 ky (Fig. 2B, Supplementary Fig. S18). The trajectories started to increase at the onset of the last glacial ∼115 kya, when one of the CAA/WG individuals appeared to diverge from the rest. We observed an increase in rate ∼30 kya, when the individuals from NEG/Sv and SEG appeared to diverge, with the onset of a sharp increase in CAA/WG following later, around the end of the Last Glacial Maximum, at which point PSMC inferences are no longer reliable (Li and Durbin 2011).

To gain insights into the more recent demographic history of narwhals, we used the population-level dataset of 24 individuals to estimate the site frequency spectrum (SFS, Supplementary Fig. S15-16). SMC++ (Terhorst et al. 2017) and stairwayplot (Liu and Fu 2015, 2020) revealed broadly similar demographic trajectories across the three populations, characterized by population growth during the last glacial period. The timing of the onset differed somewhat between analyses and similar to PSMC, SMC++ results indicated an increase in effective population size ∼115 kya with a steeper increase starting ∼30 kya (Fig. 2C). The stairwayplot estimated a population expansion ∼50 kya (Supplementary Fig. S19). We also used the program GONE (Santiago et al. 2020), see supplementary methods, but the results were not reliable because we observed a spurious, extremely large increase in *N_e_* in recent years in CAA/WG and NEG/Sv possibly due to low diversity and small sample sizes.

Finally, we investigated the population divergence process and migration events among the three populations by fitting a simple demographic model to the 3D-SFS using fastsimcoal2 (Excoffier et al. 2021) and moments (Jouganous et al. 2017; Supplementary Fig. S20). Both approaches support an initial split between the east and west of Greenland, with divergence time estimates ranging from ∼300 kya (fastsimcoal2) to ∼70 kya (moments; Supplementary Table S2). The subsequent divergence between NEG/Sv and SEG was estimated at ∼36 kya by fastsimcoal2 and ∼17 kya by moments.

Both methods further indicate that the West–East split was accompanied by a pronounced reduction in the effective population size of SEG to fewer than 2,000 individuals. The pronounced temporal gap between the initial East-West split and the later divergence of NEG/Sv and SEG suggests that the SEG population remained small for an extended period, corresponding to hundreds to a few thousand generations. Consistently, SEG shows the lowest effective population size across both modelling frameworks.

Absolute divergence times are confounded by migration, and accordingly the youngest split times were estimated using SMC++, which does not account for migration, as <6,500 years (Supplementary Table S4). Most split times estimated by fastsimcoal2 and moments correspond to the timing of increases in *N_e_* estimated in PSMC, SMC++ and stairwayplot. This further suggests that changes in *N_e_* may also reflect changes in connectivity. In agreement with our other analyses, migration rates in both fastsimcoal2 and moments were the lowest between CAA/WG and SEG, which are also the most distant populations geographically (Supplementary Table S2).

### Selection

We investigated whether behavioural differences among the three narwhal populations, ranging from long-distance migration to more resident behaviour, may drive local adaptation. To test for signatures of positive selection in each population, we applied population branch statistics (PBS) (Yi et al. 2010) and PCAdapt (Luu et al. 2017; Privé et al. 2020). Using PBS, we identified several outlier genomic regions, i.e. exceeding the 99.95th or 99.99th percentiles, that were highly differentiated along the branch of one of the three populations relative to the other two (Fig. 3, Supplementary Fig. S21).

**Figure 3.**
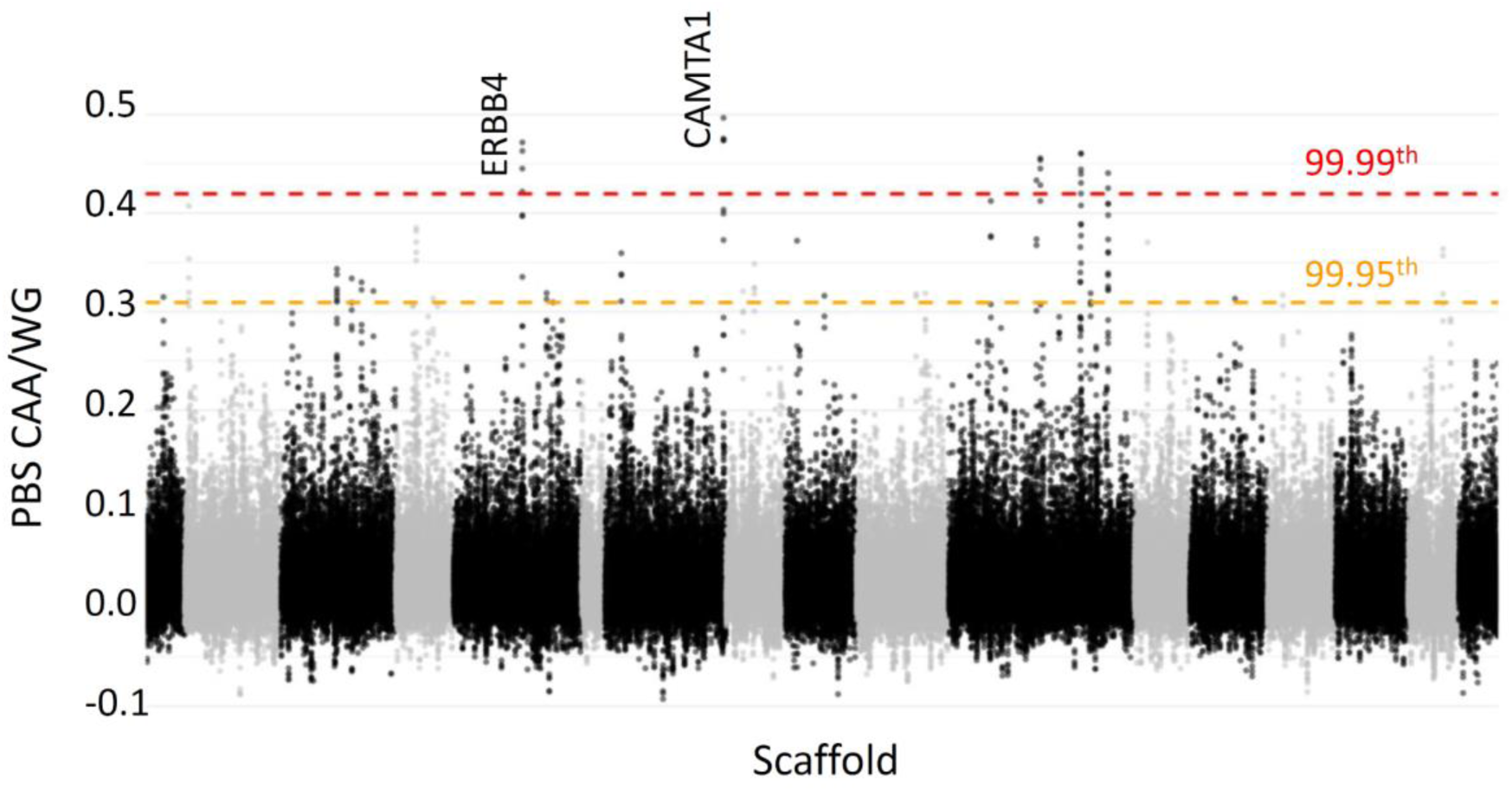
Local adaptation in the CAA/WG narwhal populations. Manhattan plot of the PBS values in windows of 50 k base pairs (kb) using a 10-kb step across large autosomal scaffolds - different scaffolds are coloured in successions of black and grey. The dashed orange and red lines indicate the 99.95th and 99.99th percentiles, respectively, above which we consider the values to be outliers. The genes in the 99.99th outliers linked to long-term memory (ERBA4 and CAMTA1) are indicated; the genotype plots can be found in Supplementary Fig. S25. Plots for the two other populations and the complete list of outlier genes can be found in Supplementary Table S5.

For each PBS outlier region, we identified overlapping genes (Supplementary Table S5). Among genes in PBS outlier regions potentially linked to ecology, we found one gene in NEG/Sv and one in SEG associated with feeding physiology: ACSL1 (fatty acid metabolism) in NEG/Sv and PNLIPRP3 (digestion and absorption) in SEG (Suzuki et al. 1990; Stelzer et al. 2016). However, these two genes were not identified as outliers in PCAdapt.

PBS identified two genes in outlier regions in CAA/WG, which were also outliers in PCAdapt analyses, CAMTA1 and ERBB4, which are associated with cognitive abilities and memory (Huentelman et al. 2007; Miller et al. 2011; Bas-Orth et al. 2016; Tian et al. 2017; Skirzewski et al. 2018; Domínguez et al. 2019) (Fig. 3, Supplementary Fig. S23-27). Variation within these two genes across individuals indicated higher derived allele frequency in the CAA/WG population than in the other two populations (Supplementary Fig. S25).

### Genetic diversity

To investigate diversity and inbreeding levels in narwhals, we computed heterozygosity (He) values and runs of homozygosity (ROH). We found very low but similar levels of diversity in the three populations (mean He 0.00038, SD = 0.000028, Fig. 4A, Kruskal–Wallis χ^2^ = 1.02, df = 2, P = 0.60).

**Figure 4.**
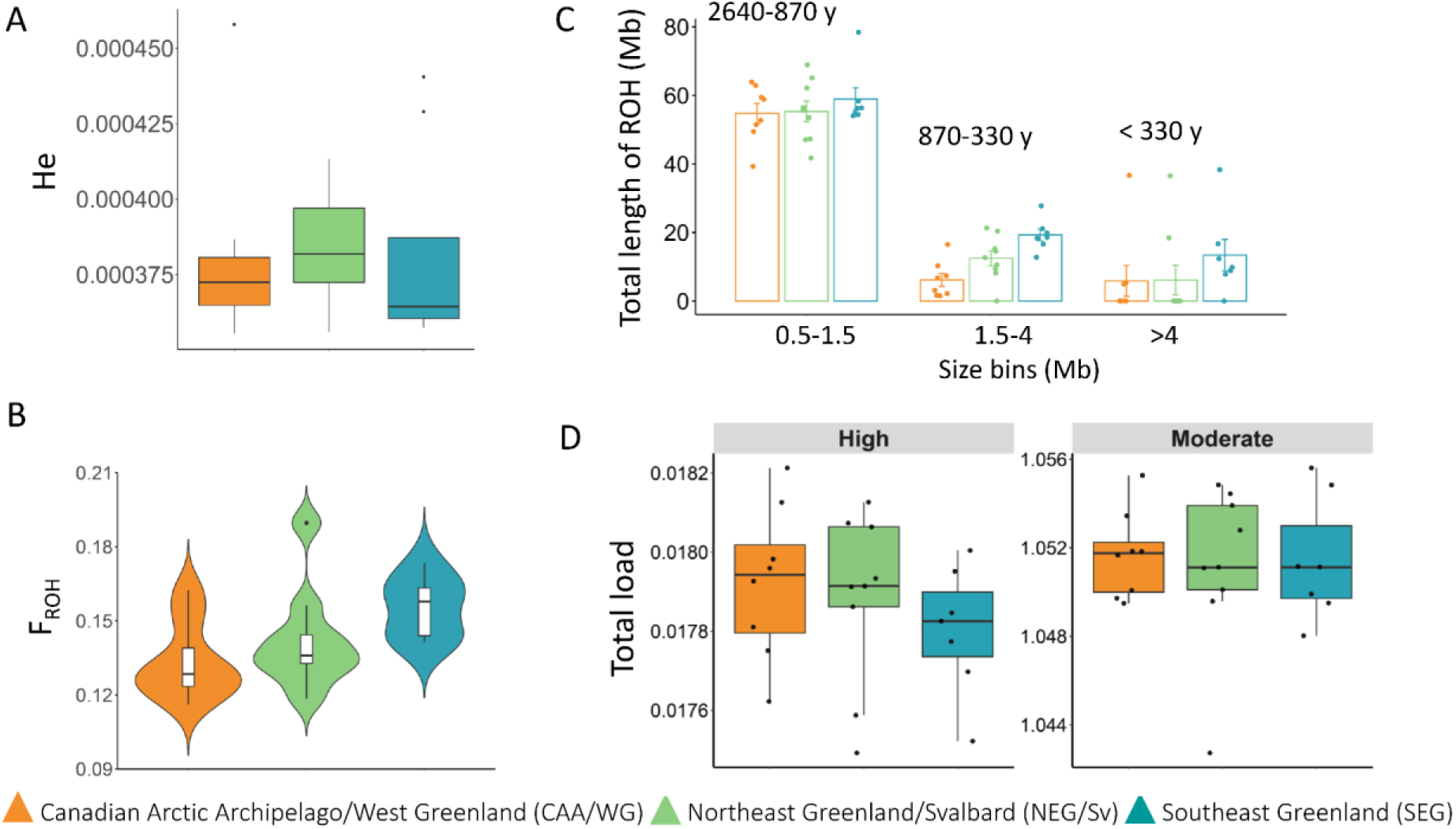
Diversity levels in the three inferred narwhal populations. A) Mean individual heterozygosity (He) estimated as the proportion of heterozygous genotypes from the site-frequency-spectrum. B) Mean individual inbreeding coefficient estimated as the fraction of the non-repetitive part of the autosomes in ROH per individual (F_ROH_). C) Inbreeding levels across time. Mean total lengths of ROH per population for three lengths categories corresponding to different time periods: 0.5 to 1.5 Mb (2640 to 870 years), 1.5 Mb to 4 Mb (870 to 330 years) and more than 4 Mb (less than 330 years). D) Genetic load (total load) calculated as the ratio between the total count of derived alleles in high or moderate impact mutations by the derived allele count in synonymous mutations for each population.

We further investigated diversity levels within populations by estimating ROH for each individual. We found similar results when using the window-based approach implemented in Plink (Purcell et al. 2007) and the MCMC approach implemented in bcftools (Narasimhan et al. 2016), as well as when varying parameters (Supplementary text).

SEG individuals had the highest mean ROH length, but differences were not statistically significant among the three populations (ANOVA, F(2,21)=2.32, P=0.12, Supplementary Fig. S28). Additionally, the proportion of the genome in ROH (F_ROH_) did not differ significantly among the three populations (ANOVA, F(2,21)=3.10, P=0.07), although SEG narwhals had a significantly higher F_ROH_ than the CAA/WG narwhals, after correcting for multiple comparisons (Tukey HSD P_CAA/WG-SEG_ = 0.05, Fig. 4B).

Patterns of ROH across length classes indicated that all three populations shared similar long-term demographic histories. There were no differences in total size of small ROH (0.5 to 1.5 Mb, reflecting ∼2640 – 870 years ago, Supplementary Fig. 4C, see statistical details in the supplementary material), which is consistent with long-term low effective population sizes. The two populations east of Greenland showed higher levels of intermediate-length ROH (1.5–4 Mb), indicating higher levels of inbreeding during the past ∼870–330 years. SEG individuals also had the highest mean total length of long ROHs (>4 Mb), reflecting elevated recent inbreeding (∼330 years). Although not all pairwise comparisons remain significant after correcting for multiple testing (see Supplementary material), our results indicate relatively more recent inbreeding in the smallest population (SEG).

### Genetic load

We investigated how the long-term demographic history of narwhals, particularly their persistently low effective population sizes (*N_e_*; Supplementary Fig. S18), has shaped the accumulation of genetic load. Theory predicts that species with long-term low *N_e_* purge highly deleterious mutations while accumulating mildly deleterious ones (Bertorelle et al. 2022; Dussex et al. 2023), and our results are consistent with these predictions. We observed a low proportion of highly deleterious mutations alongside an accumulation of moderately deleterious mutations. Both mutation types were more common in heterozygous genotypes (masked/inbreeding load) than in homozygous genotypes (realized load; Fig. 4D, Supplementary Fig. S29–S30). No significant differences in genetic load were detected among the three populations, regardless of mutation type or load category (masked or realised).

## Discussion

Narwhals are considered one of the most sensitive Arctic marine mammal species to ongoing climate change (Laidre et al. 2008); they have a narrow environmental and ecological niche, low genetic diversity, high fidelity to migratory routes and are highly sensitive to anthropogenic noise. By integrating our genomic DNA with ecological data from the literature, we show that both abiotic factors (climatic and environmental conditions) and biotic factors (behavioural traits) have shaped patterns of narwhal population structure and diversity. We also show a correlation between inbreeding levels and population size.

### Three distinct populations

Based on the analysis of 117 nuclear genomes sequenced at low coverage (mean = 2.73x, min = 0.15x, max = 11.99x, modal = 0.3×), we identified three genetically and geographically distinct populations: Canadian Arctic Archipelago/West Greenland (CAA/WG); Northeast Greenland/Svalbard (NEG/Sv); and Southeast Greenland (SEG) (Fig. 1A,B). The greatest genetic differentiation was between CAA/WG and SEG. This pattern is consistent with their clear separation based on RAD-sequencing data (Lopes et al. 2025) and on skin and bone collagen *δ*¹³C and *δ*¹⁵N stable isotope analyses (Watt et al. 2013; Louis et al. 2021a, 2025). Bone-collagen isotope analyses showed no overlap in the ecological niches of narwhals from West and Southeast Greenland, indicating long-term ecological divergence and region-specific foraging. The isotopic distinction was less clear between the NEG/Sv and SEG populations (Louis et al. 2025).

The CAA/WG narwhal population is the largest of the three, and encompasses an estimated ∼150-160,000 individuals, within nine of twelve recognised management stocks (Hobbs et al. 2019). We were unable to identify genetic sub-structuring within this group, in contrast with a recent study of Canadian narwhals using higher-coverage data (modal coverage of 10×) (de Greef et al. 2024). The discrepancy is likely due to the extremely low genetic variation across the narwhal genome (Westbury et al. 2019), and thus higher-coverage data such as used by de Greef et al. (2024) are required to resolve finer-scale population structure, which is valuable for stock assessments.

Southeast Greenland is the smallest of the three identified populations. The population is of particular conservation concern and has experienced severe declines over the past two decades (Hobbs et al. 2019; Hansen et al. 2024), driven by overharvesting and the loss of summer sea ice (Heide-Jørgensen et al. 2020a). Aerial surveys conducted in 2022 estimated the population size to be 365 individuals (CV=0.68, 95% CI: 173–769), representing a decline of approximately 85% since 2008 (Heide-Jørgensen et al. *in press*; NAMMCO 2023). Biological samples collected from the Inuit hunt and official hunters catch reports of the SEG population have also indicated a skewed demographic structure, including a declining proportion of pregnant females, an overrepresentation of old males and a marked absence of calves and juveniles (Garde et al. 2022).

Our study presents nuclear genomic data from narwhals in the northeast of their range. These data showed that narwhal in Northeast Greenland and Svalbard belong to the same population. Narwhals are protected within the Northeast Greenland National Park and are not accessible to hunters due to the remoteness of the region. Consequently, apart from recent aerial surveys conducted over sections of the park that estimated the presence of several thousand individuals (Hansen et al. 2024), little information is available on narwhals in this region; the five NEG samples in our study were collected via biopsy sampling from a helicopter operating from an ice breaker. Narwhals occur year-round between Northeast Greenland and Svalbard (Ahonen et al. 2019), but their migratory behaviour remains unknown. In Svalbard, narwhals are rarely observed near shore, although aggregations of several hundred individuals have been reported occasionally (Storrie et al. 2018) and narwhals are consistently recorded on passive acoustic recorders during winter in ice-covered areas in the east of Svalbard (Llobet et al. 2023). Our genomic analysis included data from three juveniles that were satellite-tagged in 1998 (Supplementary Table S1), all of which remained relatively close to Svalbard during the 4-46 days of tags operation (Lydersen et al. 2007).

To further characterize genetic variation within and among the three identified populations, and to investigate their joint and population-specific demographic histories, signatures of local adaptation and levels of inbreeding and genetic load, we focused on the 24 narwhal individuals sequenced at higher coverage (Fig. 1C).

### Demographic history shaped by climate

Our deep-time demographic reconstruction based on individual genomes indicated similar long-term trajectories across the three populations, consistent with a shared demographic history and indicating little-to-no population structure (Fig. 2B, Supplementary Fig. S18). Both individual- and population-based demographic reconstructions showed an increase in *N_e_* starting ∼115 kya during the onset of the Last Glacial Period, when the trajectory of CAA/WG diverged (Fig. 2). Both approaches also indicated a sharp increase in the rate of *N_e_* ∼30 kya at the start of the Last Glacial Maximum (LGM), at which time the trajectories of NEG/Sv and SEG diverged. An increase in *N_e_* may be due to a real change in population size, or alternatively shifts in population connectivity, given that variation in *N_e_* reflects changes in the rate at which genetic lineages coalesce (Mazet et al. 2015). Narwhals are closely associated with sea ice and polar water masses (Laidre et al. 2004), and population increase may have been facilitated by colder climates that expanded available habitat. During winter, narwhals preferentially inhabit deep offshore waters where they forage intensively; selected areas have high sea ice concentrations, but with predictable open-water patches (<5%) where they can breathe (Laidre and Heide-Jorgensen 2005; Kenyon et al. 2018). Narwhals are found within a narrow temperature range year-round, but especially during their feeding season. It is unclear whether this is driven by prey availability, thermoregulatory constraints, predator refugia or a combination (Heide-Jørgensen et al. 2020b). Regardless, our demographic reconstructions suggest population subdivision has been a factor contributing to the observed increases in nuclear *N_e_*.

A more recent, post-glacial population expansion was suggested by the demographic trajectory of 121 mitochondrial genomes—covering most of the individuals analysed herein—and the star-like patterns of several lineages within their haplotype network (Louis et al. 2020). This may have been driven by the expansion of favourable habitat, as ecological modelling suggested narwhal habitat increased between the LGM and the present (Louis et al. 2020). A similar post-glacial pattern of population and habitat expansion has been reported in several other polar marine predators, including Arctic beluga whales (Skovrind et al. 2021) and bowhead whales (*Balaena mysticetus*) (Foote et al. 2013), and other North Atlantic baleen whales (Cabrera et al. 2022b). The expansion of North Atlantic baleen whales was correlated with an increase in their prey. Although similar analyses are not available for narwhal prey species, and a post-glacial expansion seems counterintuitive given their close association with sea ice and polar water, other factors, such as increase in prey abundance, may have contributed to the increase.

We used several approaches to estimate divergence times among the three narwhal populations, but none of them were reliable for inferring absolute values. However, in agreement with our demographic reconstruction, all approaches identified the split between western and eastern Greenland as older, occurring four to eight times earlier than the divergence between the two eastern populations (Supplementary Table S4).

Our population structure and admixture analyses showed NEG/Sv as intermediate between CAA/WG and SEG (Fig. 1D, Supplementary Fig. S12-14). This is consistent with higher estimates of inferred migration between CAA/WG and NEG/Sv, and between NEG/Sv and SEG in our demographic modelling (Supplementary Fig. S20, Supplementary Table S2). We estimated the directionality of gene flow to be from west to east (Fig. 2A), which is supported by the detection of narwhal DNA in marine sediment cores from the seas north of Greenland (Schreiber et al. 2025). The Nares Strait is the waterway connecting Baffin Bay (Atlantic Ocean) and the Lincoln Sea (Arctic Ocean). The strait was closed until ∼9 kya by shelf-based ice (Jennings et al. 2011; Georgiadis et al. 2020). Narwhal DNA retrieved from a marine sediment core from Lincoln Sea dating ∼8 ky BP (Schreiber et al. 2025) and fossil evidence from ∼7 ky BP on the northern coast of Ellesmere Island (Evans 1989) indicate that narwhals were present in the strait relatively soon after it opened. Furthermore, environmental proxy data retrieved from sediment cores from Nares Strait and Lincoln Sea suggest the influx of warmer Atlantic water from the west after the opening of the Nares Strait (Schreiber et al. 2025), supporting our finding of gene flow occurring from west to east.

In summary, our combined findings based on demographic reconstruction and modelling, estimates of gene flow, drift and phylogeographic patterns of mitochondrial diversity, suggest populations west and east of Greenland were isolated and diverged before or during the last glacial period, and came into contact again in the early Holocene possibly via migration from west Greenland and Canada into northeast Greenland after the Nares Strait opened. This indicates that past climate-associated environmental shifts, in particular the colder period of the last glacial period, affected the demographic history of narwhals, and suggests that changes in *N_e_* reflect changes in both population size and connectivity.

### The role of restricted ecological niche and site fidelity to migratory routes

Our inferred genetic structure of narwhals mirrors broad-scale habitat discontinuities across their range in terms of differences in temperature and sea ice occurrence, and is correlated with differing behavioural traits. Narwhals are restricted by a narrow temperature range throughout the year (Heide-Jørgensen et al. 2020b). This may limit dispersal ability and sustain population structure between CAA/WG and SEG. Waters around the southern tip of Greenland lack sea ice and may be too warm for narwhals (Chambault et al. 2020). Extensive tagging of narwhals in both CAA/WG and SEG has not shown migration from east to west (or vice versa) (Heide-Jørgensen et al. 2013, 2015). No narwhals have been observed further south than 66°N in West Greenland or 64°N in East Greenland (Heide-Jørgensen et al. 2015; Hobbs et al. 2019), supporting our inference of limited ongoing gene flow. The lack of connectivity is also supported by the lack of overlap in their ecological niches (Louis et al. 2021a, 2025). Stable isotope analyses of skin tissue have indicated that narwhals in SEG have a more pelagic diet than in CAA/WG, including more capelin *Mallotus villosus* (Watt et al. 2013). This west-east ecological differentiation is also observed in other Arctic top predators including bowhead whales (Westbury et al. 2025) and polar bears (Westbury et al. 2023), suggesting the potential influence of isotopic value differences at the base of the food web in addition to diet variation.

Persistent year-round sea ice in the Nares Strait might restrict narwhal movement north of Greenland between CAA/WG and NEG/Sv. Narwhals overwinter in areas with dense drift ice, but still require access to cracks and leads for breathing. Ancient DNA from marine sediments suggests that narwhals have occupied the Nares Strait and Lincoln Sea continuously since the early Holocene (Schreiber et al. 2025); however, their present-day use of this region remains poorly understood. Narwhals have been observed in the Nares Strait during summer in recent years, but the winter and spring distribution of these animals is unknown, as the strait is typically covered by landfast ice from October through mid-July (Heide-Jørgensen et al. 2024). Identifying contemporary migratory patterns across the strait will be critical for understanding genetic connectivity, particularly given that mating occurs from late winter to early spring (Heide-Jørgensen and Garde 2011; Kelley et al. 2015).

Behavioural traits, in particular fidelity to migratory routes, may drive the observed differentiation between NEG/Sv and SEG. Tagging studies of dozens of narwhals from Scoresby Sound (SEG population), the world’s largest fjord system, reveal highly consistent interannual movement patterns (Heide-Jørgensen et al. 2015). These narwhals stay within Scoresby Sound during summer and early autumn, migrate to the continental shelf pack ice southeast of the fjord in late autumn, overwinter there through spring, and return to the fjord the following summer (Heide-Jørgensen et al. 2015, 2020b).

Narwhals in NEG/Sv appear to behave differently. Acoustic recorders document that at least some animals in this population remain in the offshore pack-ice year-round (Ahonen et al. 2019). Our findings of genetic differentiation between NEG/Sv and SEG suggest that strong fidelity to established migratory routes restrict dispersal and contribute to the maintenance of population subdivision. However, we cannot disentangle this effect from isolation-by-distance, as no intermediate samples are available between Northeast Greenland and SEG, which are separated by a distance of >1,000 km. Similar mechanisms involving culturally transmitted behaviours, such as migratory routes or foraging strategies, have been shown to drive and maintain population structure in highly mobile cetaceans, including southern right whales (*Eubalaena australis*), humpback whales (*Megaptera novaeangliae*), killer whales and bottlenose dolphins (Baker et al. 1986; Louis et al. 2014; Carroll et al. 2015; Foote et al. 2016)).

We investigated whether variation in migratory behaviour among narwhal populations may reflect local adaptation. Narwhals in CAA/WG undertake the longest known annual migration for this species, with individuals from Somerset Island, Canada, travelling up to 3,000 km round-trip between their summer and winter grounds (Dietz and Heide-Jørgensen 1995; Dietz et al. 2008; Westdal et al. 2010; Heide-Jørgensen et al. 2013). Narwhals in SEG also migrate, but over shorter distances–approximately 700 km round trip–between their offshore wintering areas and their coastal summering grounds (Heide-Jørgensen et al. 2015).

The long-distance migration in Canadian narwhals likely requires complex memory capabilities. Two genomic regions identified as putatively under selection by both PBS and PCAdapt, with PBS indicating selection in CAA/WG, were associated with cognitive abilities. The CAMTA1 gene is associated with episodic and long-term memory performance (Huentelman et al. 2007; Miller et al. 2011; Bas-Orth et al. 2016) and ERBB4 is associated with learning, spatial and social memories, as well as the brain’s mental workspace for holding and using information (Tian et al. 2017; Skirzewski et al. 2018; Domínguez et al. 2019). Differences in genes associated with long-term memory have also been identified in populations of peregrine falcons (*Falco peregrinus*) that differ in their migratory distances (Gu et al. 2021). Our results suggest that differences in ecology and behaviour contribute to local adaptation in narwhals, as observed in other highly social cetaceans (Foote et al. 2016; Louis et al. 2021b). However, we note the findings of our selection analyses should be interpreted with caution, as outliers can arise from demographic effects or other selective processes (Bank et al. 2014), and we acknowledge that our conclusions are primarily correlative.

### Low diversity and implications for conservation

We found similar levels of heterozygosity in all three narwhal populations, which are among the lowest recorded in mammals (Morin et al. 2020). Our ROH results, representing inbreeding levels over relatively recent timescales (past few thousand years), were inversely correlated with current population sizes (Fig. 4). We observed the lowest proportion of ROH in the genomes from CAA/WG, which has the largest estimated population size of the three narwhal populations (Hobbs et al. 2019). The highest proportion of ROH was in SEG, which has the lowest estimated population size of only a few hundred individuals (NAMMCO 2019, 2023). SEG also showed a higher proportion of long ROH (> 4 Mb), which reflects the past 10-20 generations, indicating recent inbreeding in this population. Although we lack a comprehensive abundance estimate for NEG/Sv, a recent survey estimated several thousand individuals in the Northeast Greenland National Park (Hansen et al. 2024).

Narwhals have persisted at small *N_e_* and have had low genome-wide diversity for at least the past 600,000 years (Figure S18) (Westbury et al. 2019). Empirical and theoretical evidence indicate that long-term low *N_e_* can reduce genetic load due to purging of highly deleterious alleles (e.g. in Alpine ibex, *Capra ibex*, vaquita, *Phocoena sinus* (Grossen et al. 2020; Robinson et al. 2022), making populations less susceptible to the impact of bottlenecks or inbreeding depression. However, while highly deleterious alleles are generally purged by selection, moderately or mildly deleterious alleles may increase in frequency, as small *N_e_* will reduce the effects of purifying selection (Bertorelle et al. 2022; Dussex et al. 2023). Our genetic load results are concordant with this prediction; across the three narwhal populations, we found a relatively low proportion of highly deleterious mutations and an accumulation of moderately deleterious mutations (Fig. 4D, Supplementary Fig. S22-23). Also, both types of mutations were present at higher frequency in heterozygous (i.e. masked/inbreeding load) than homozygous genotypes (i.e. realised load). This finding contradicts expectations of a lower masked genetic load in species, such as narwhals, that have maintained small population sizes over long time periods (Bertorelle et al. 2022; Dussex et al. 2023). It may reflect a relatively recent increase in population size, as indicated by demographic reconstruction based on both nuclear (Fig. 2B,C) and mitochondrial genomes (Louis et al. 2020).

Recent studies have shown that fitness can be reduced by the accumulation of mildly to moderately deleterious mutations, even when highly deleterious mutations are efficiently purged (e.g. Soay sheep *Ovis aries* and killer whales (Stoffel et al.; Kardos et al. 2023)). This raises concerns for the smallest and rapidly declining narwhal population in Southeast Greenland, where individuals have a large proportion of their genome in intermediate and long ROH. During prolonged bottlenecks in small populations, the increase in mildly deleterious mutations, or the expression of previously masked highly deleterious mutations, may reduce fitness (Kardos et al. 2023; Dussex et al. 2023). Our finding of a relatively high masked load despite a long-term low *N_e_* in SEG raises additional concern if the population continues to decline, as this could lead to the unmasking of deleterious mutations and a subsequent decrease in fitness.

### Conclusions

Our results indicate that past climate fluctuations have shaped the demographic trajectories of the three narwhal populations. Contemporary population structure is likely maintained by habitat discontinuities and strong fidelity to migratory routes, which may also contribute to patterns of local adaptation. We present a genomic assessment of narwhal vulnerability and show that inbreeding levels are inversely related to current population sizes. The smallest and rapidly declining population, Southeast Greenland (SEG), exhibits signatures of recent inbreeding and a relatively high masked genetic load, raising concerns given ongoing pressures from overharvesting, rapid environmental change and increasing human activity.

## Material and methods

### Samples

We collected tissue samples from 121 narwhals spanning the species’ range, and representing all 12 narwhal stocks recognized by the North Atlantic Marine Mammal Commission (NAMMCO) (Fig. 1A). For narwhals a stock is defined as a summer aggregation and serves as a management unit based on distribution, telemetry, ecological and, in some cases, genetic data (Hobbs et al. 2019). Samples were obtained during summer between 1982 and 2019 through subsistence hunts, biopsy sampling associated with satellite-tagging studies, and, for Northeast Greenland, targeted helicopter-based biopsy sampling (Supplementary Table S1). Samples were sent to Denmark under CITES exemption number DK003. 112 of these samples have been previously analysed using mitochondrial genomes (Louis et al. 2020).

From 120 samples, we first conducted preliminary analyses of population structure and then selected 24 individuals from the three genetic clusters identified—CAA/WG, NEG/Sv, and SEG (Fig. 1C), and included an additional individual from SEG, to increase sample size which happened to be related with one of the 24 individuals—for sequencing at sufficient coverage to support analyses of demographic history, inbreeding and selection. Because these samples included several related individuals, the final datasets comprised 117 individuals for the range-wide analyses (referred to as range-wide dataset) and 24 individuals for the more population-level analyses (referred to as population-level dataset).

We included two available high-coverage genomes from CAA/WG in the deep-time demographic history analysis (Westbury et al. 2019): PRJNA508363 and PRJNA520934.

We used the beluga (sample SAMN12242088), the sister species within Monodontidae and the finless porpoise (sample SRR940959), a member of the closely related family Phocoenidae (McGowen et al. 2020), as outgroups and to reconstruct ancestral allele states (Yim et al. 2014).

### DNA laboratory work

We extracted DNA from tissue samples using the Qiagen Blood and Tissue Kit following the manufacturer’s protocol with minor modifications (details in the supplementary methods). Libraries were prepared on sheared DNA and sequenced according to two different protocols. For 102 samples, libraries were built using Illumina NeoPrep following the NeoPrep Library Prep System Guide applying default settings and sequenced on an Illumina HiSeq 2500 with 80bp SE technology. For 31 of those 102 samples as well as for 19 additional samples, libraries were built using a second library preparation method: the BEST protocol (i.e. Blunt-End Single-Tube library building for modern and ancient DNA (Carøe et al. 2018)). Of these 50 samples, libraries were sequenced on an Illumina HiSeq Xten with 150bp PE technology for 48 samples and on a Novaseq 6000 with 150bp PE technology for 28 samples. Of these 26 were sequenced on both platforms. To avoid any issue with differences in sequencing platforms, we did not merge data from Hiseq and Novaseq and only used the Novaseq data for the 25 samples of the population-level dataset but are making all data accessible on SRA (Table S1). We only merged data from both platforms to increase coverage in PSMC analyses.

### Bioinformatics

#### Mapping

Sequencing reads were processed using Paleomix v. 1.2.13.1 (Schubert et al. 2014). After trimming using AdapterRemoval v. 2.2.2 (Schubert et al. 2016), the remaining reads were mapped to the narwhal reference genome (GCF_005190385.1) using bwa v. 0.7.15 with the mem algorithm (Li and Durbin 2009) requiring a mapping quality score of 30 for the narwhal raw sequencing reads, while we used a mapping quality score of 20 for the finless porpoise and beluga raw sequencing reads given we were mapping to another species (the narwhal). Picard-tools v. 2.6.0 (http://broadinstitute.github.io/picard/faq.html) was used to remove duplicate reads. Indel realignment was performed using GATK v. 3.8.1 (McKenna et al. 2010). We set a filter of 20 for the base score quality as we included Novaseq data.

We only kept the 22 largest scaffolds and further filtered the data to only keep the autosomes using bedtools v. 2.25.0 and samtools v. 1.2 (Li et al. 2009; Quinlan and Hall 2010). To identify the scaffolds corresponding to the X and Y chromosomes, we used the SeXY pipeline (Cabrera et al. 2022a) and ran a PCA for each large scaffold to see if we could detect any clustering by sex. We also removed repeat regions and regions of excessive coverage, because they could be the result of unmasked repeat regions in nuclear mitochondrial DNA (NUMTs), or some other mapping artifact (see details in the supplementary material). Coverage was estimated using ANGSD v. 0.939 (Korneliussen et al. 2014).

#### Variant calling

We called SNPs taking genotype uncertainty into account by calculating genotype likelihoods in ANGSD for both the range-wide (n=120) and population-level (n=25) datasets, creating a beagle file and keeping SNPs with a minimum MAF of 0.05 and having data in a minimum of 75% of the individuals. In ANGSD, all analyses described below were run considering only SNPs with a phred quality of 20 and a mapping quality score of 30.

We called genotypes (i.e. generation of a vcf file) for the population-level dataset using bcftools v. 1.12 mpileup multiallelic and rare-variant calling on the filtered bam files (Li et al. 2009; Danecek et al. 2014). Variable sites with a minimum phred score quality of 20, minimum depth of 3, and genotype quality of 20 were retained in vcftools v. 0.1.16 (Danecek et al. 2011). We kept SNPs with a minimum MAF of 0.05 and having genotype data in a minimum of 75% of all the individuals and a minimum of five individuals in each of the three populations. The vcf file was also filtered for monomorphic and non-biallelic sites. After filtering, 727,253 SNPs were kept in the narwhal vcf, and 823,928 SNPs and 768,978 SNPs when including the finless porpoise and the beluga as outgroups respectively.

### Population structure analyses

#### Inferring relatedness

We ran NgsRelate (Korneliussen and Moltke 2015; Hanghøj et al. 2019) to identify pairs of related individuals both in the range-wide (n=120) and population-level (n=25) datasets using the pairwise relatedness measure (Hedrick and Lacy 2015). Our goal was to remove strongly related individuals, as those were driving our population structure results. We removed three individuals from pairs of related narwhals from the CAA/WG population in the range-wide dataset, reducing it to 117 individuals and one individual from the SEG population-level dataset, therefore reducing it to 24 individuals. We plotted the pairwise relatedness coefficients in heatmaps for the range-wide dataset both for n=120 and n=117 using R v. 4.0.5 (R Core Team 2021) and packages ggplot2 (Wickham 2016a), dplyr (Wickham et al. 2023), tidyr (Wickham and Girlich 2022) and readr (Wickham et al. 2022).

#### Inferring linkage disequilibrium

We used NgsLD (Fox et al. 2019) to obtain a set of unlinked SNPs for both range-wide and population-level datasets. A set of unlinked sites was produced, considering that SNPs are in LD until 20 kb, using a minimum weight of 0.5. Population structure analyses were run on the set of 486,709 unlinked SNPs for the 117-sample dataset and on a set of 274,852 SNPs for the 24-sample dataset. All the other analyses (demographic history or selection) were run on the unpruned SNPs (excluding the two scaffolds showing clustering by sex) because either the program included the pruning (e.g. Treemix) or linkage information is important for the inferences.

#### PCA and admixture analyses

To infer population structure, we used PCAs in PCAngsd (Meisner and Albrechtsen 2018) and admixture analysis in NGSAdmix (Skotte et al. 2013) using 486,709 unlinked SNPs for the range-wide dataset and 274,852 unlinked SNPs for the population-level dataset. NGSAdmix was run 100 times for each *K* value between 2 and 5. Consistency between runs was checked, and the runs with the highest likelihood for each *K* value were plotted in R.

We used evalAdmix (Garcia-Erill and Albrechtsen 2020) to estimate the best number of clusters (*K*) using the NGSadmix results from both datasets. EvalAdmix outputs a matrix of pairwise correlation of residuals between individuals and values close to 0 indicate a good fit of the data to the admixture model. For *K*=3, we observe a better fit of the data to the admixture model than for *K*=2 (see supplementary methods, Figure S6). We also checked admixture proportions amongst the 100 runs for various R values, and found they were consistent among runs for *K*=3, but not for higher values of *K*.

We ran PCAngsd on the range-wide and population-level datasets. We did the same for a dataset of all the CAA/WG narwhals (n=101) to test for fine-scale population structure within this region (CAA/WG dataset). We plotted all of the results from PCAngsd using R.

For the range-wide and CAA/WG datasets, we also ran the single-read PCA in ANGSD v.0.939, which involves random sampling of a single read for each sample at each site. Note that we did not perform LD pruning for this analysis, as it was run directly on the bam files. We plotted the results using R.

#### Fixation statistics

For the range-wide dataset, we estimated *F*_ST_ using ANGSD (Nielsen et al. 2012; ^K^orneliussen et al. 2014^)^. First, we calculated the site allele frequency spectrum likelihood (saf) based on individual genotype likelihoods. Then, we computed the unfolded 2D-SFS from the saf for each pair of populations in winsfs (Rasmussen et al. 2022) using the finless porpoise as the ancestral state, as well as the folded 2DSFS. We estimated *F*_ST_ between pairs of populations using the realSFS fst index function in ANGSD. We used the same procedure for the population-level dataset, although we used ANGSD to estimate the 2D-SFS for these data.

All analyses below were run on the population-level dataset, and in one case (deep-time demographic history) single individuals with high sequencing coverage.

### Evolutionary relationships

We reconstructed the evolutionary relationships of the three inferred narwhal populations using Treemix and several admixture graph analyses. All analyses were run twice, including either the finless porpoise or the beluga as an outgroup/root. Results were similar, regardless of the outgroup.

#### Treemix

We reconstructed the evolutionary relationships of the three inferred narwhal populations as a Maximum Likelihood bifurcating tree using TreeMix v. 1.13 (Pickrell and Pritchard 2012). We first ran TreeMix ten times for 0 to 4 migration events. We estimated the optimal number of migration events as 1 using the optM R package (https://cran.r-project.org/web/packages/OptM/index.html). We then ran TreeMix 100 times for 1 migration event and obtained a consensus tree and bootstrap values using the R package BITE (Milanesi et al. 2017). We estimated the residual covariance matrix for the consensus tree using TreeMix.

#### Admixture graphs - qpBrute and Admixture Bayes

We reconstructed the evolutionary history of the three populations as an admixture graph (Patterson et al. 2012). We used a heuristic search algorithm, qpBrute (https://github. com/ekirving/qpbrute), which explores the space of all possible admixture graphs of a given maximum complexity, under a brute-force approach (Ní Leathlobhair et al. 2018; Liu et al. 2019). The analysis included 823,928 SNPs and 768,978 SNPs when including the finless porpoise and the beluga as outgroups, respectively. The package tried all possible six starting graph orders, and we found three graphs with no *f4* outliers among a total of 14 unique graphs, two of which were mirror graphs of each other. We computed the mean log-likelihoods of the three models and their Bayes Factors using the MCMC algorithm implemented in the R package ADMIXTUREGRAPH v. 1.0.289. We assessed the convergence of the chains using the output from the R package CODA v. 0.19-490 (Plummer et al. 2006). The Bayes factors indicated that the models had similar log-likelihoods, and were equally likely (*K*<0.1).

We used Admixture Bayes to estimate high-probability admixture graphs (Nielsen et al. 2023). This Bayesian approach uses a reversible jump Markov Chain Monte Carlo (MCMC) to sample high-probability admixture graphs. It has the advantage of searching the entire state space to find the best fitting graph(s) and to provide a level of confidence in the sampled graphs. We ran three independent MCMC chains with 400,000 iterations and 16 chains and used the default values for the “analyzeSample” step. We assessed convergence using the EstimateConvergence R script (https://github.com/avaughn271/AdmixtureBayes).

#### *D*-stats /*f3*-statistics

We used *D*-statistics and *f3*-statistics to further assess the relationships among the three identified narwhal populations. *D*-statistics describe an excess of shared derived alleles between taxa which could be the result of introgression or ancestral population structure. It thus allows for the detection of departures from the ‘tree-ness’ of a given topology (Green et al. 2010; Durand et al. 2011; Patterson et al. 2012). We calculated *D*-statistics of the form (H1, H2; H3, OUT): (SEG, CAA/WG; NEG/Sv, OUT), (SEG, NEG/Sv; BB, OUT) and (NEG, BB; SE, OUT) using the qpDstat tool in Admixtools v. 7.0.2 (Patterson et al. 2012). To test for significance, we calculated a *Z*-score based on blocked jackknife estimates of the standard deviation of the *D*-statistics.

We also estimated *f3*-statistics, which measure allele frequency correlations between populations (Patterson et al. 2012), to test whether a target population (C) is admixed between two source populations (A and B). We estimated *f3*-statistics using the threepop function in Treemix. We estimated the statistics for the data with the individuals correctly assigned to their populations and after randomly reshuffling the individuals among populations to test whether our results differ from random.

### Demographic reconstruction

To reconstruct the demographic history of the narwhals across both ancient and recent timescales, we applied a combination of PSMC, SMC++, and Stairway Plot. PSMC captures long-term changes in effective population size (*N_e_*) using a single high-quality genomes, while SMC++ extends this resolution to more recent periods by incorporating multiple genomes. Stairway Plot complements these approaches by using the site frequency spectrum (SFS) to infer the more recent past.

#### Population size reconstruction with PSMC, SMC++ and stairwayplot

We used the pairwise sequential Markovian coalescent model (PSCM (Li & Durbin 2011) to infer deep-time changes in *N_e_* through time. To infer more recent demographic history we used Sequential Markov Coalescent + plenty of unlabelled samples (SMC++) (Terhorst et al. 2017) and stairwayplot (Liu and Fu 2015, 2020), which are based on population scale data and can include individuals with lower coverage than PSMC. We also used SMC++ to estimate divergence times among the three inferred populations. We scaled the results of all three analyses using a generation time of 30 years (Garde et al. 2015), and the per-generation mutation rate previously estimated for the species: 1.56e-08 per generation (Westbury et al. 2019). We plotted the results in R with packages ggplot2, scales (Wickham 2016b), RcolorBrewer (Neuwirth 2014) and reshaped2 (Wickham 2007).

#### PSMC

We ran PSMC on single individuals with the highest coverage from each of the three identified populations: two published genomes from CAA/WG (PRJNA508363 and PRJNA520934, coverage of 59.9 and 56.0×) (Westbury et al. 2019), three genomes sequenced as part of this study including two from NEG/Sv (20.3× and 15.0×) and one genome from SEG (12.7×). We only used autosomes, and removed the scaffolds where individuals clustered by sex. We ran the PSMC inference using the parameter values for human autosomes (see supplementary material). We also ran PSMC with 100 rounds of bootstrapping.

We acknowledge that two of our samples have depth of coverage <20x which may lead to false negative detection of heterozygous sites and impact the results (Nadachowska-Brzyska et al. 2016). However, we observed very similar trajectories across samples until ∼40,000 years ago, and PSMC results are not reliable in the past ∼20,000 years in any case (Li & Durbin 2011).

#### SMC++

For SMC++, we generated the vcf file as described earlier, but no MAF filter was applied because the analysis is based on the SFS. The vcf file was converted to SMC++ format using the vcf2smc function for each retained autosomal scaffold. The regions identified as repeats and showing excessive coverage were included in a mask file. SMC++ estimates the SFS conditioned on the TMRCA of a single individual, called the “distinguished” individual hereafter. We made the distinguished individual vary over three individuals with the highest coverage within each of the three populations. Population size histories were estimated using the estimate option in SMC++ using the default settings for the estimate function. To estimate population splits, we first estimated population histories using the estimate option, i.e. the marginal estimates. We computed the joint frequency spectrum (2D-SFS) for each pair of populations using the vcf2smc function. We used the split function to refine the marginal estimates into an estimate of the joint demography and estimate divergence times between pairs of populations.

#### Stairwayplot

We used stairwayplot v. 2 to reconstruct changes in *N_e_* through time. We ran the analysis using the unfolded SFS, generated using both winSFS and ANGSD, and obtained similar demographic trajectories; the results based on the SFS generated in winsfs are presented herein. We ran the methods both including and excluding singletons, and obtained similar trajectories, results including singletons are therefore presented.

#### Demographic Modeling with FASTSIMCOAL2 and moments

To investigate the historical processes that shaped the current population structure and take gene flow into account, we estimated demographic parameters from the SFS using FASTSIMCOAL2 v. 2.7 (Excoffier et al. 2013, 2021) and moments (Jouganous et al. 2017).

To gain insights into the historical processes shaping the observed population structure of narwhals, which is potentially influenced by post-glacial environmental shifts, we estimated demographic parameters from the SFS. We inferred present and ancestral population sizes, divergence times, and historical migration events among each pair of populations. We focused on the 24 higher-coverage samples: eight from CAA/WG, seven from SEG and nine from NEG to construct a three-population site frequency spectrum (3D-SFS). We generated the 3D-SFS using allele frequency genotypes obtained from winsfs, as described above. To reduce bias in determining ancestral allelic states, we used the folded 3D-SFS, based on the estimates produced by winsfs.

Demographic inference was performed using two approaches: the composite likelihood method implemented in fastsimcoal2 (Excoffier et al. 2013, 2021), which simulates genetic data under complex demographic scenarios to fit the observed SFS, and moments (Jouganous et al. 2017), which uses diffusion approximations to model allele frequency changes over time. We assumed a simple divergence model with gene flow among three populations corresponding to CAA/WG, SEG, and NEG/Sv (Supplementary Fig. S20). In this model, ancestral narwhal populations west and east of Greenland diverged simultaneously from a common ancestor, followed by a subsequent split between the NEG/Sv and SEG populations.

For demographic inference using fastsimcoal2, the expected likelihood was obtained through one million coalescent simulations (-n1,000,000) and 100 conditional optimization cycles (- L100). We performed 100 independent runs with different starting parameter values and retained the estimates from the run that achieved the maximum likelihood. To assess model fit, we visually compared the expected and observed marginal one-dimensional SFS, as well as the three two-dimensional SFS pairs, to evaluate how well the model could reproduce the observed data. We used a mutation rate of 1.56e-08 mutations/ site/ generation (Westbury et al. 2019) and a generation time of 30 years (Garde et al. 2007) to scale the demographic history to years. A summary of the defined parameters and their search ranges are given in Supplementary Table S3. Input files including template and parameter estimation files are available in the github repository (see data availability section). To obtain confidence intervals for the estimated parameters, we applied a non-parametric bootstrap approach. Specifically, we generated 100 bootstrapped 3D-SFS datasets by resampling sites from the observed SFS with replacement, assuming independence among SNPs. Each bootstrapped SFS included the same number of sites as the original dataset, including monomorphic sites. For each bootstrap replicate, we approximated the expected SFS using 1 million coalescent simulations and 100 optimization cycles. We initialized the optimization using the parameter set that maximized the likelihood in the observed SFS (.pv), which allowed us to reduce the number of replicates to three per bootstrap. Confidence intervals were then computed from the parameter estimates of the best-fitting replicate (i.e., the one with the highest likelihood) for each bootstrap.

We further estimated the three population demography using moments to robustly replicate the results from fastsimcoal2. Similar to fastsimcoal2, we used the 3D-SFS and estimated a simple 3 population divergence model with migration, with an East-West split followed by a subsequent split in the East between the SEG and the NEG/Sv populations. Moments uses a diffusion approximation approach to estimate the SFS given the demographic parameters. To estimate the demographic parameters, we used 100 replicates with randomly chosen starting parameters and ran all of these replicates to convergence. The models were compared across replicates using likelihoods computed under a multinomial distribution for the observed SFS using the expected SFS as the parameter for the multinomial distribution.

In addition to the simple divergence model, we added growth to all three populations after the population split. Adding the population growth to the three populations led to a complete lack of convergence for all replicates. Thus, we excluded population growth models from any further analyses.

### Local adaptation

#### PBS

To test for signatures of positive selection in the three geographically distinct populations, we used population branch statistics (PBS) (Yi et al. 2010) and PCAdapt (Yi et al. 2010; Luu et al. 2017; Privé et al. 2020). PBS identifies alleles that have undergone significant frequency shifts in a target population, compared to two other populations. The approach identifies highly differentiated genomic regions in each population. We computed population branch statistics and *F*_ST_ in ANGSD for each population. We used the site allele frequency likelihoods generated for each population, and the 2D-SFS generated as described in the fixation statistics section. We computed *F*_ST_ using the Hudson estimator and PBS using the realSFS function in ANGSD in windows of 50 kb with a 10 kb slide. We adopted an outlier-based approach, and selected the most extreme values of the empirical distribution— specifically >99.95th or >99.99th percentile. For each identified outlier region, we identified overlapping genes and analyzed haplotype patterns of genes of interest across samples.

We estimated nucleotide diversity and Tajima’s D for each population, based on the 24-sample dataset, using ANGSD from the SFS (Nielsen et al. 2012), and plotted the results for the scaffolds including outlier genes of interest. We generated the SFS as described in the fixation statistics section. We calculated nucleotide diversity for each site and estimated Tajima’s D from the SFS using a sliding-window size of 50 kb and a step size of 10 kb.

### PCAdapt

We also performed a genome scan using the package PCAdapt (Luu et al. 2017; Privé et al. 2020). PCAdapt identified SNPs whose allele frequency differences among populations, characterised by PCA, are unexpectedly large, as outlier candidates for local adaptation. It has the advantage of not requiring grouping of individuals a priori into populations, and is robust when there are admixed individuals. We first evaluated the best number of principal components (PCs) to consider by examining the percentage of variance explained on a scree plot, and which PCs ascertain putative population structure on the PCA. PCAdapt models the correction between individual allele count and the first *K* PCs. It evaluates the association strength using the robust Mahalanobis distance, and detects outliers by converting these distances into P-values based on a chi-squared distribution with *K* degrees of freedom. We ran the analyses with all SNPs and after SNP thinning to account for linkage disequilibrium using a window size of 100 SNPs and an r2 threshold of 0.1. We identified outlier SNPs (α = 0.1) in each analysis accounting for a false-discovery rate of 10% by calculating q-values with the qvalue function in R package “qvalue” (Storey et al. 2020). We also applied the Bonferroni or Benjamini– Hochberg correction, as a comparison.

### Diversity statistics

#### Heterozygosity

We estimated heterozygosity for each of the three populations based on the 1D-SFS generated by ANGSD, as the product of the number of variants in the second column of the est.ml file, which corresponds to the heterozygotes, divided by the total number of sites. We tested whether mean heterozygosity was significantly different among the three populations using Wilcoxon tests.

#### ROH

Runs of homozygosity (ROH) arise when an offspring receives two copies of the same ancestral haplotype from both parents. ROH can be broken up by recombination in subsequent generations. The ROH length distribution across the genome can provide information about demographic history and levels of inbreeding (Ceballos et al. 2018). We used both the window-based approach implemented in Plink v. 1.90b6.21 (Purcell et al. 2007) and the MCMC approach implemented in bcftools v. 1.15 to identify ROH. We estimated ROH with no MAF filter for all the parameters combination but made a comparison including a filter for MAF 0.05.

In Plink, we set the sliding window size to 300 kb with a minimum of 50 SNPs and a minimum density of 1 SNP per 40 kb, as required to call a ROH. To take genotyping errors into account, we allowed up to 3 heterozygous sites per 300 kb window within ROHs (Ceballos et al. 2008). We set the length between two SNPS to be 1000 kb for them to be considered in two different ROH. We ran the estimations varying some of the parameters (see details in the supplementary material). We tried different values (5, 10, 15, 20) for the number of missing calls per window and evaluated when the number of usable windows plateaued, which was 10.

We identified ROH using bcftools with the option G30 specifying that allele frequencies should be estimated from each of the three inferred populations, or from the total 24 individuals (as a comparison). For both programs, we calculated mean ROH length and maximum ROH length per individual. We estimated F_ROH_ as the fraction of the autosomal scaffolds within ROH per individual, after removing filtered regions such as repeats grouped by population. We plotted mean ROH length, maximum ROH length, and mean F_ROH_ per population in R using the package ggplot2.

The length of the homozygote tracts decline as a function of recombination rate and time, and the parts of the genome in ROH of different lengths can provide information about past changes in effective population size. We calculated the age of the ROH using the formula from Thompson (2013): g = 100/2*ROHlength*r, where g is the age of the ROH in generations and r the recombination rate in cM. We used the recombination rate of humans of 1.133 cM/Mb (Dumont and Payseur 2008). We converted the age from generation to years using the generation time of 30 years for narwhals (Garde et al. 2007).

Differences in inbreeding statistics among the three populations were compared using ANOVA and Tukey’s post hoc tests - implemented in the R package multcomp (Hothorn et al. 2008) or Kruskal–Wallis and Dunn-tests - implemented in the R package dunn.test (Dinno 2017), depending on whether the data satisfied the required assumptions: normality and homogeneity of variances.

#### Genetic load

To infer genetic load, we filtered out genotypes with a depth of coverage <5×, as well as heterozygous genotypes, where the minor allele was not supported by at least 2 reads. We annotated the vcf file with snpeff v. 5.2c (Cingolani et al. <u>2012</u>) and the available annotation for the narwhal. We identified the ancestral state of each allele by creating a consensus sequence using the beluga and finless porpoise genomic data (see details in the supplementary material) and selecting the most common base using the doFasta 2 option together with doCounts 1 in ANGSD.

We used Plink, vcftool, bedtools v. 2.30.0 and R following Humble et al. (2023) pipeline to incorporate the ancestral state and polarise the alleles in the vcf file. Note that we removed the sites with warnings after running snpeff, and restricted the analysis to the sites where the reference is ancestral and the alternate is derived based on our outgroup (roughly 94% of the sites). We also ran the analysis varying the filters.

Following Rasmussen et al. (2023) we counted the total number of derived alleles categorised as high impact and moderate impact (total allele count) for each of the 24 individuals, as well as separated by zygosity: in homozygous state (realized allele count) and in heterozygous state (masked allele count). Moderate-impact variants are nondisruptive variants that might change protein effectiveness (i.e. missense mutations). High-impact variants are assumed to disrupt the protein, and can cause protein truncation, loss of function or initiate nonsense-mediated decay.

To take differences in coverage into account, we normalized the derived allele count by the derived allele count per sample in synonymous positions, to estimate the total load, realized load and the masked load. To compare the three inferred populations, we calculated the means of relative counts of genetic load (for high- and moderate-impact variants) per population. We statistically compared total, masked, and realised load for high and moderate impact mutations among the three populations using a linear model (anova) after checking for normality and homogeneity of variances. Our data satisfied normality and homogeneity of variance.

## Data availability

The code and scripts are available at: https://github.com/LorenzenLab/Narwhal-range-wide-study/tree/main/code_narwhal_rangewide, the raw data will be made publicly available on SRA upon publication, Biosample accession numbers: SAMN60394159-SAMN60394279.

## Acknowledgements

This study was funded by the Carlsberg Foundation Distinguished Associate Professor Fellowship, grant no CF16-0202, and the Villum Fonden Young Investigator Programme, grant no. 13151, to E.D.L. Sample collection in Greenland was funded by the Greenland Institute of Natural Resources (GINR). Samples from Northeast Greenland and Svalbard also had funding from the Norwegian Polar Institute, the Collaboration in the North programme (previously called the Norwegian-Russian Environmental Cooperation) and the Norwegian Research Council (Icewhales programme). ML was supported by a postdoctoral fellowship from the Greenland Research Council and GINR. We thank the local hunters and organizations (e.g., Hunters and Trappers) who contributed samples from South and West Greenland and from across Nunavut, Canada. Funding for the Canadian samples was provided by Fisheries and Oceans Canada and the Nunavut Wildlife Management Board.

## Author contributions

Conceptualization: ML, EDL; data curation: ML; formal analysis: ML, BP, DV, AC, SG; funding acquisition: EDL; investigation: ML, MS, ARI; resources: SHF, EG, MPHJ, KMK, CL, LP, EDL, supervision: EDL; visualization: ML, BP, EDL; writing – original draft: ML, EDL; writing – review and editing: All authors

## Notes

### Competing Interest Statement

The authors have declared no competing interest.

