## Supplementary material for "Habitat discontinuity and fidelity to migratory routes shape narwhal genetic structure and diversity"

Supplementary Figures S1-S30
Supplementary Tables S1-S5
Supplementary Results
Supplementary Methods

#### Supplementary figures

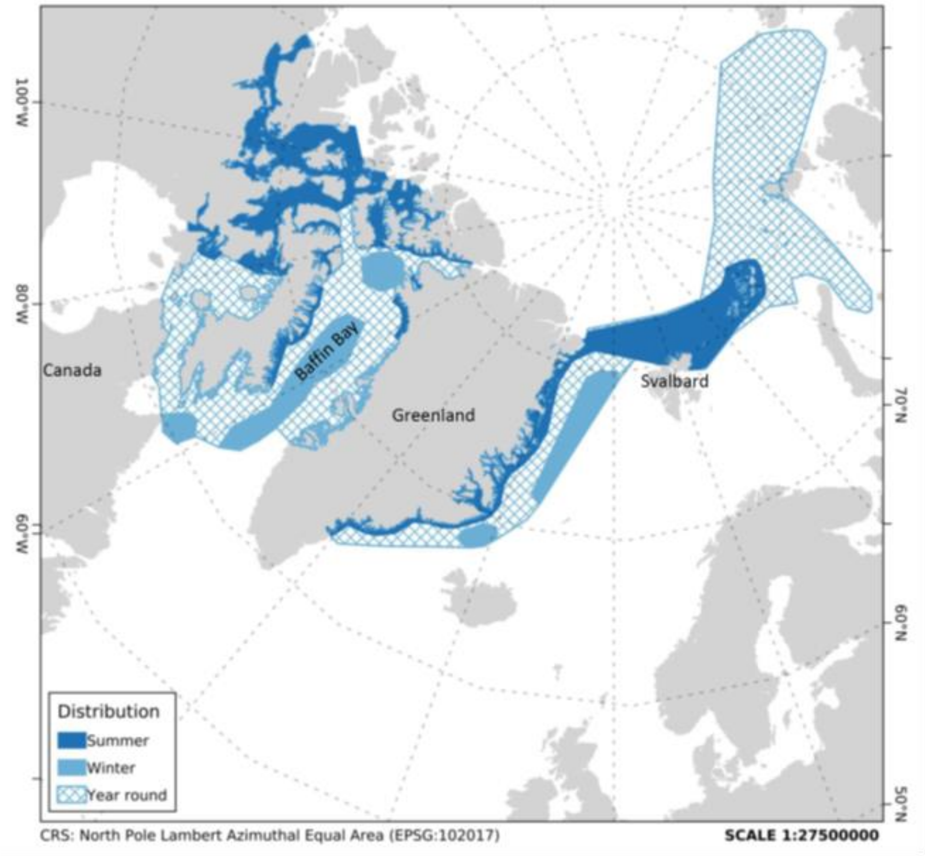


**Supplementary Figure S1. Distribution of narwhals in summer, winter and year-round.** Adapted from (Hobbs et al. 2019). The coastline data are from Wessel and Smith, 1996 (Wessel and Smith 1996). Note that a recent study indicated that narwhals are present year-found between North-East Greenland and Svalbard (Ahonen et al. 2019).

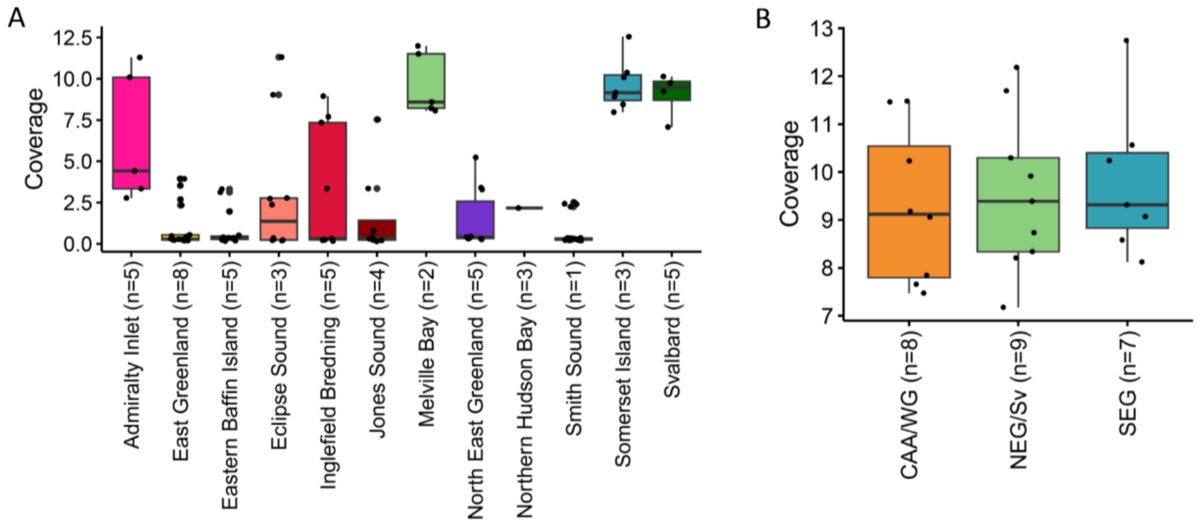


**Supplementary Figure S2. Sequencing coverage of the narwhal data analysed.** Coverage as estimated for the A) range-wide dataset based on 117 individuals from all 12 narwhal stocks, and B) population-level dataset based on 24 individuals from three populations: Canadian Arctic Archipelago and West Greenland (CAA/WG); Northeast Greenland and Svalbard (NEG/Sv); Southeast Greenland (SGE). Sample sizes are shown in parenthesis.

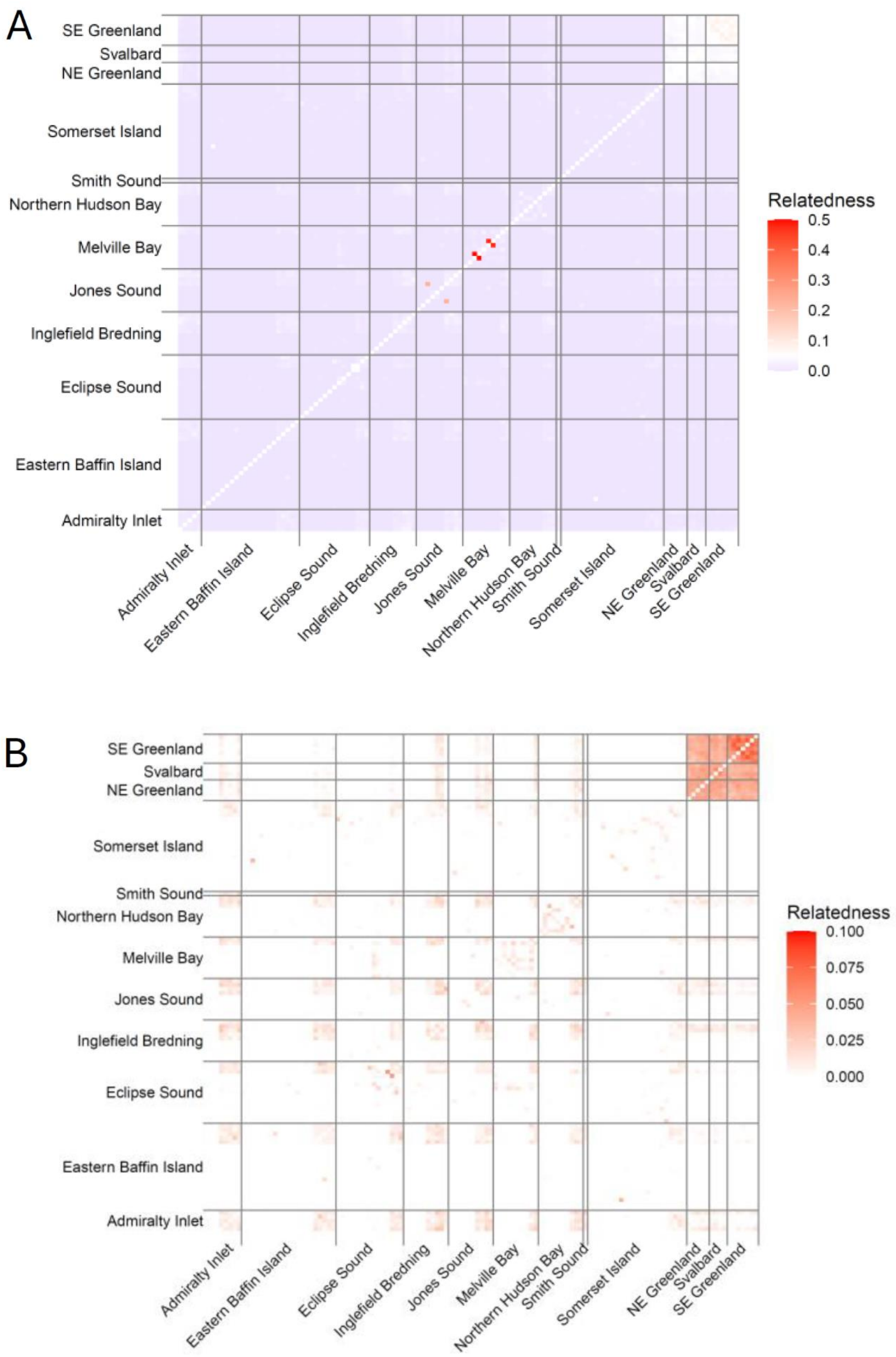


**Supplementary Figure S3. Relatedness among narwhal individuals.** Pairwise relatedness measured for A) 120 individuals and B) 117 individuals included in the final range-wide dataset, i.e. after removing one individual from each of the three pairs of relatives (r>0.1). Samples are ordered by stock/geographical region.

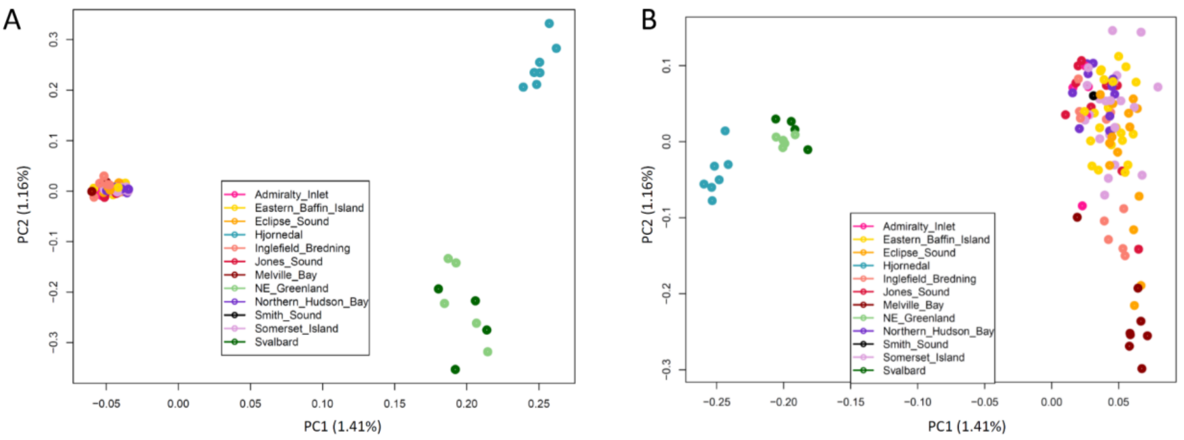


**Supplementary Figure S4**. **Population structure of 117 narwhals using the single-read PCA approach.** PCA of the range-wide data including all 12 stocks showing first and second principal components (PCs), with the proportion of genetic variance captured by each component indicated for a MAF (Minor Allele Frequency) of A) 0.01 and B) 0.05.

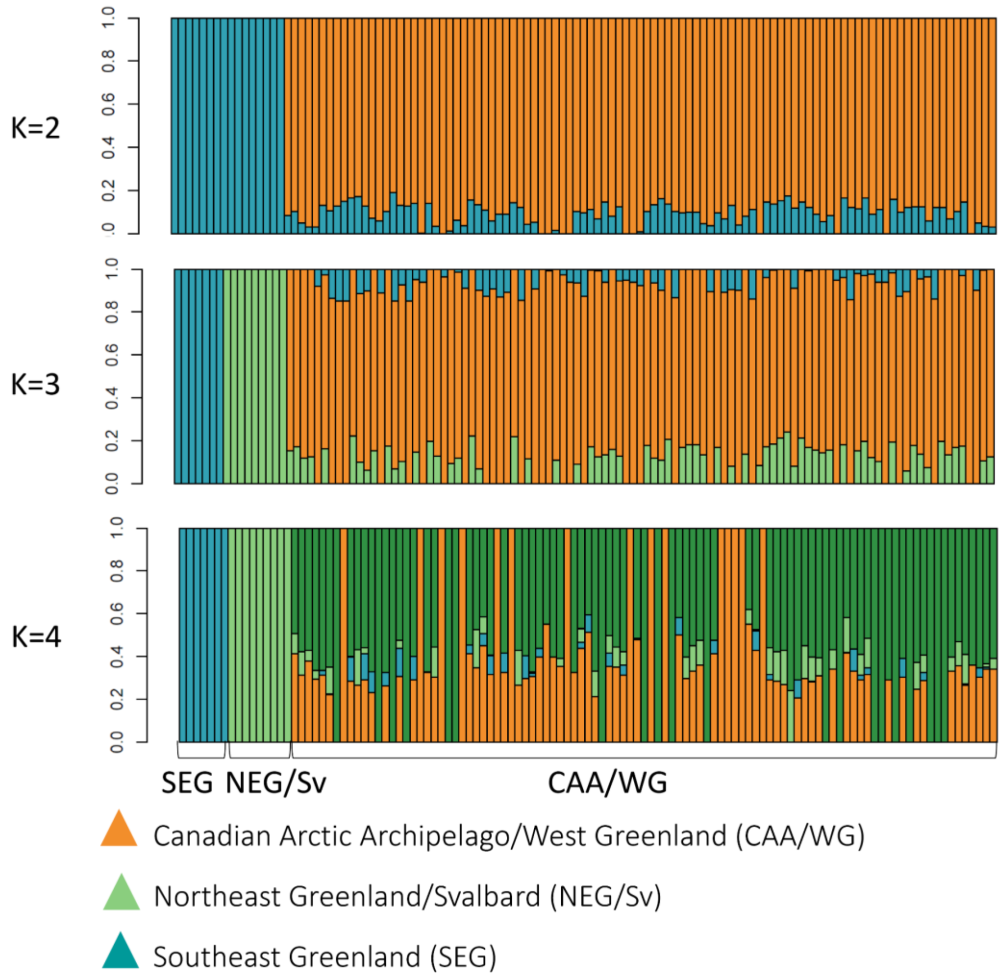

**Supplementary Figure S5. Population structure of the range-wide dataset using admixture analyses.** NGSAdmix ancestry proportions for each of the 117 individuals inferred in NGSAdmix for *K*=2 to *K*=4.

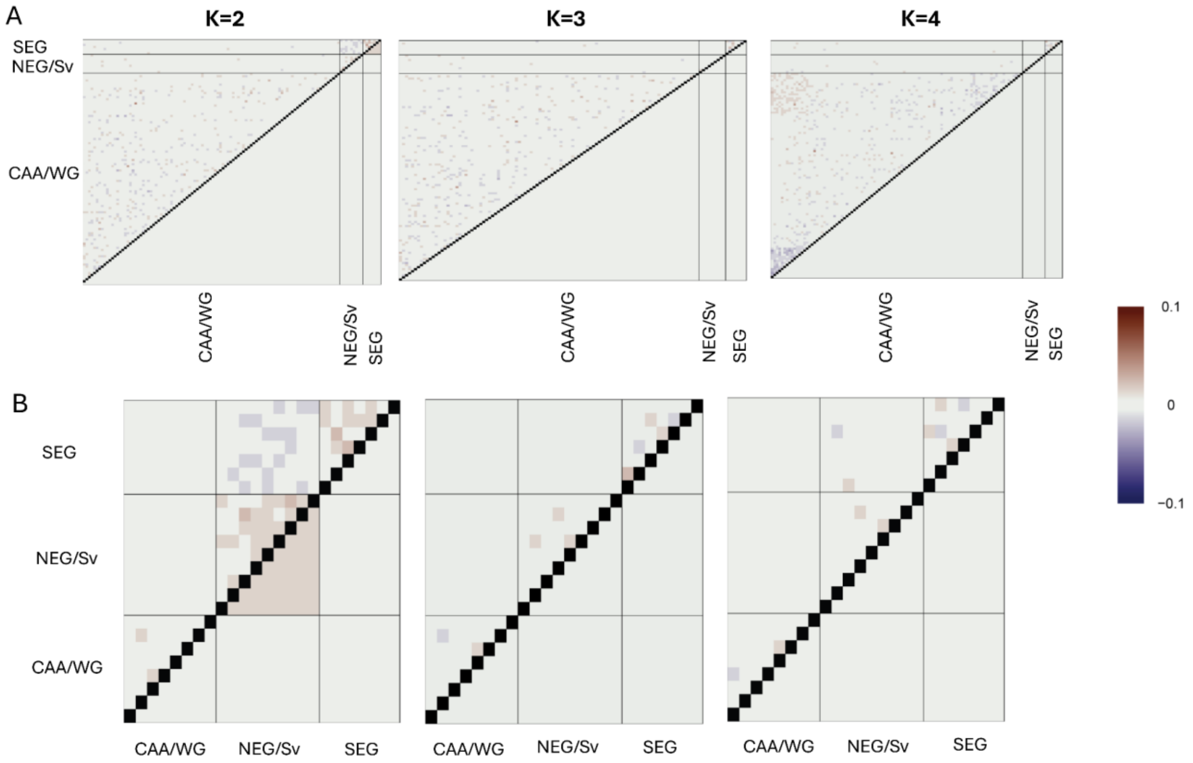


**Supplementary Figure S6. Fit of the data to the admixture models** for A) the range-wide (n=117), and B) population-level (n=24 individuals) datasets. Plot of the matrix of pairwise correlation of residuals between individuals for *K*=2 to *K*=4. Correlation values close to 0 indicate a good fit of the data to the admixture model. Populations are: Canadian Arctic Archipelago and West Greenland (CAA/WG); Northeast Greenland and Svalbard (NEG/Sv); Southeast Greenland (SEG).

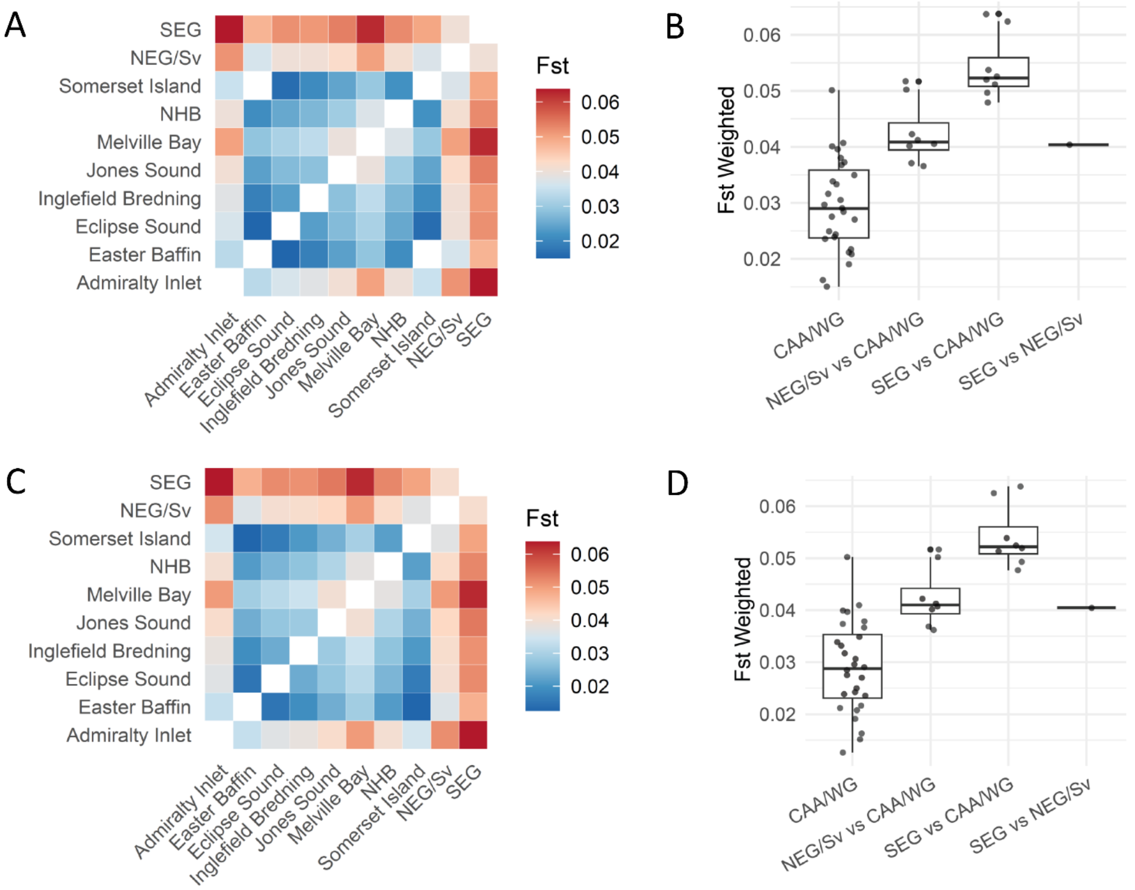


**Supplementary Figure S7. Fixation statistics among narwhal stocks.** Note that the NEG and Svalbard individuals have been grouped together based on population structure results, despite being part of two different stocks, and we only have one individual from the Smith Sound stock, it is therefore not included here. $\mathrm{F}_{\mathrm{ST}}$ between pairs of narwhals stocks based on the folded (A, B) and unfolded (C, D) 2D-SFS. Heatmaps of pairwise $F_{ST}$ (A and C) and mean pairwise $F_{ST}$ among stocks within CAA/WG and among the three regional populations (B and D). Abbreviated populations/stocks are: Canadian Arctic Archipelago and West Greenland (CAA/WG); Northeast Greenland and Svalbard (NEG/Sv); Southeast Greenland (SEG); NHB: Northern Hudson Bay.

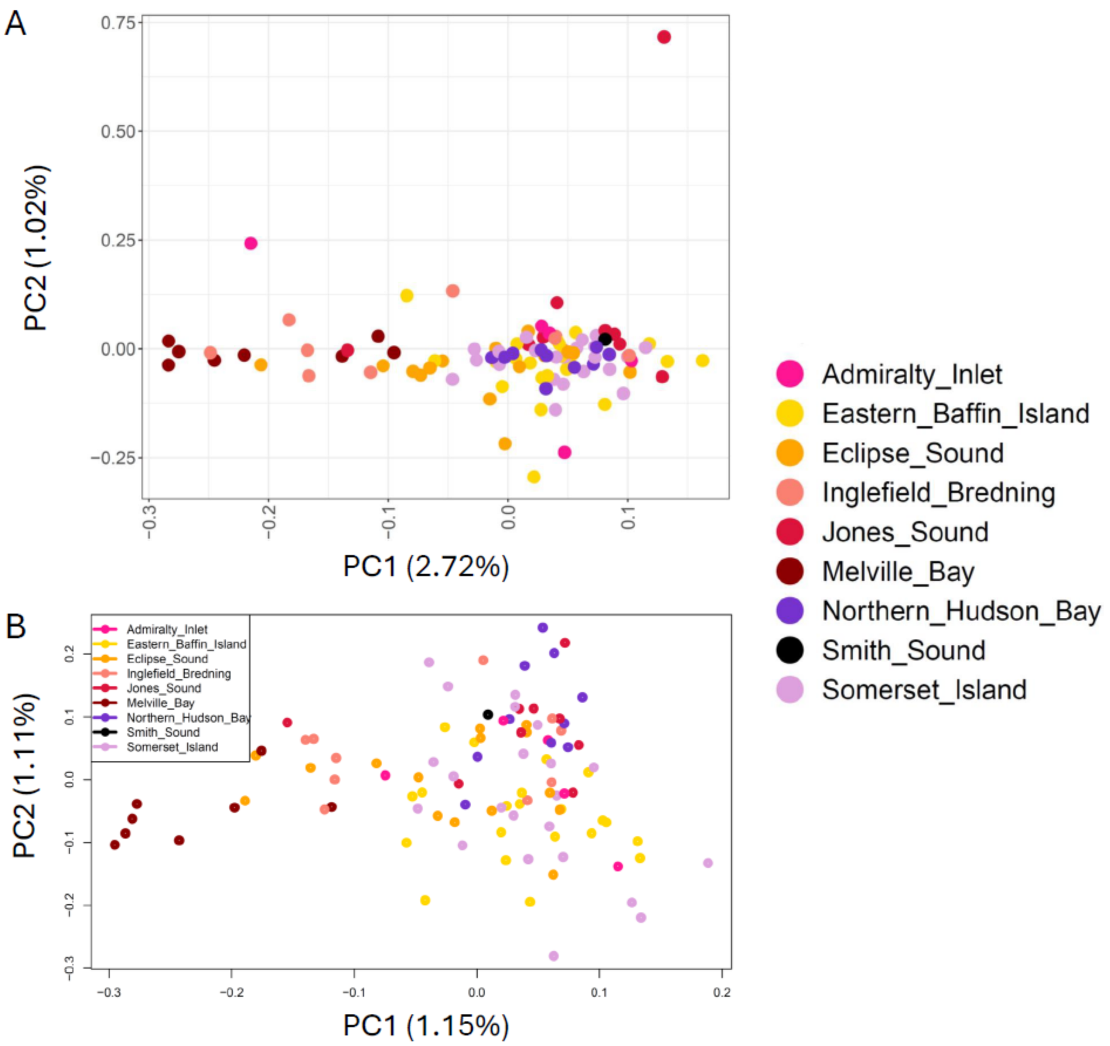


**Supplementary Figure S8**. **PCA of the 101 narwhal samples from CAA/WG** (Canadian Arctic Archipelago and West Greenland) using A) PCAngsd and B) the single read sampling method. Both graphs display first and second principal components (PCs), with the proportion of genetic variance captured by each component.

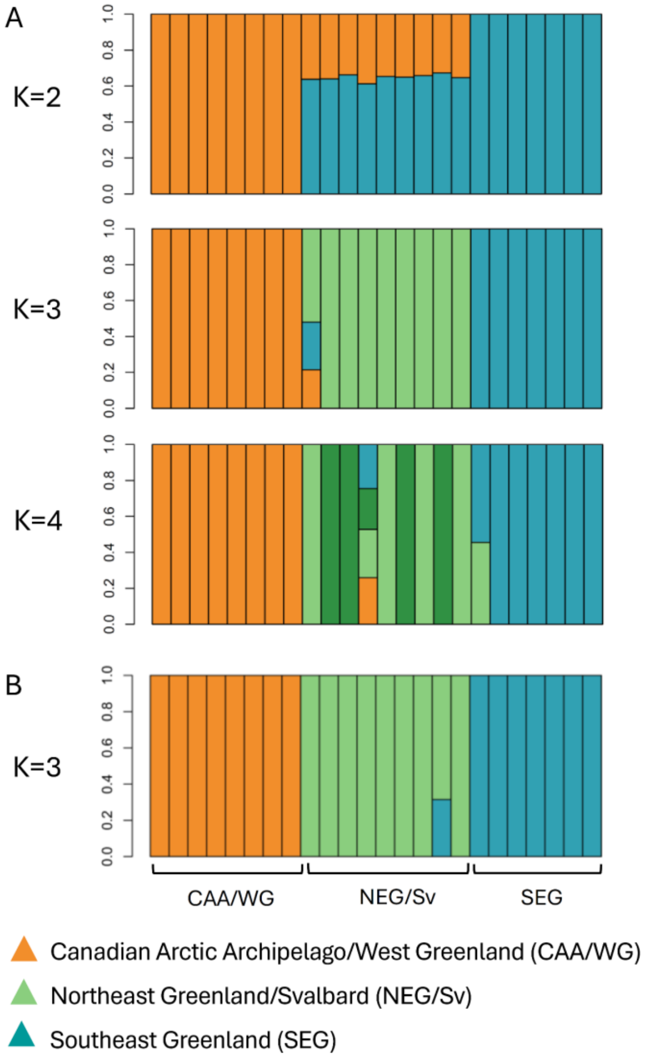


**Supplementary Figure S9. Population structure of 24 narwhals using admixture analyses.** NGSAdmix ancestry proportions for each of the 24 individuals with medium coverage data inferred in NGSAdmix for A) *K*=2 to *K*=4 for the highest likelihood and B) *K*=3 for the second highest likelihood.

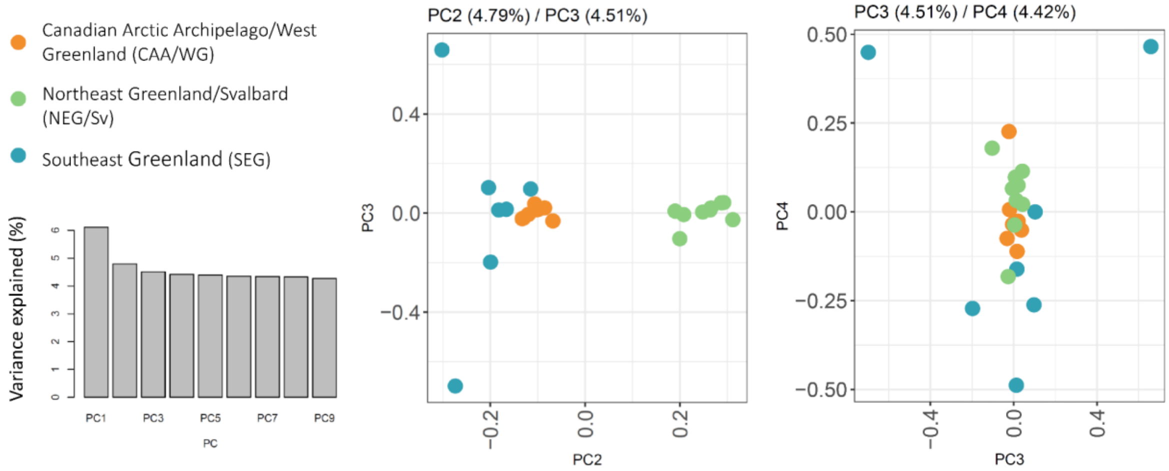

**Supplementary Figure S10. Population structure of 24 narwhals using PCAngsd for PC2-4.** PCA of the 24 narwhal samples with medium coverage using PCAngsd, displaying the variance explained for each of the principal components (PCs) between 1 and 9, the second and third PCs and the third and fourth PCs with the proportion of genetic variance captured by each component.

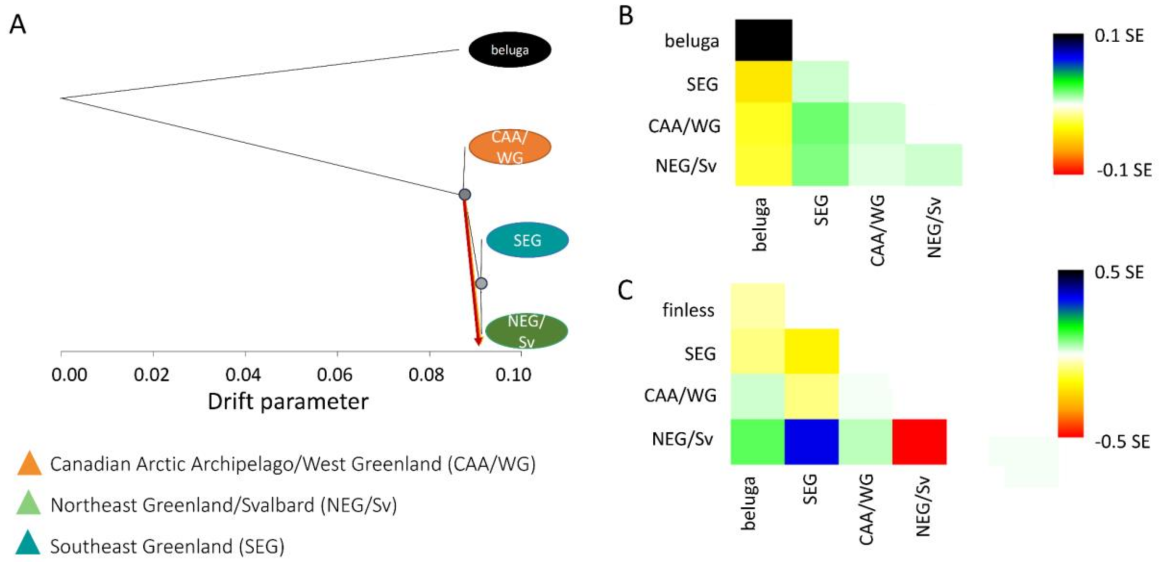


**Supplementary Figure S11**. **Treemix results.** A) TreeMix consensus tree, nodes have bootstrap values > 90%, showing the relationships among three populations as a bifurcating maximum-likelihood tree with one migration edge (M=1, in red), inferred as the best topology. Branch lengths on the horizontal axis represent the amount of genetic drift that has occurred along each branch. The tree is rooted using the beluga. Residual fit of the observed versus the predicted squared allele frequency difference, expressed as the number of SE of the deviation for the tree rooted with B) the beluga and C) the finless porpoise (see Figure 2). SE values are represented by colors according to the palette on the right.

Residuals above zero indicate populations that are more closely related to each other in the data than in the best-fit tree and have potentially undergone admixture. Negative residuals represent populations that are less closely related in the data than represented in the best-fit tree.

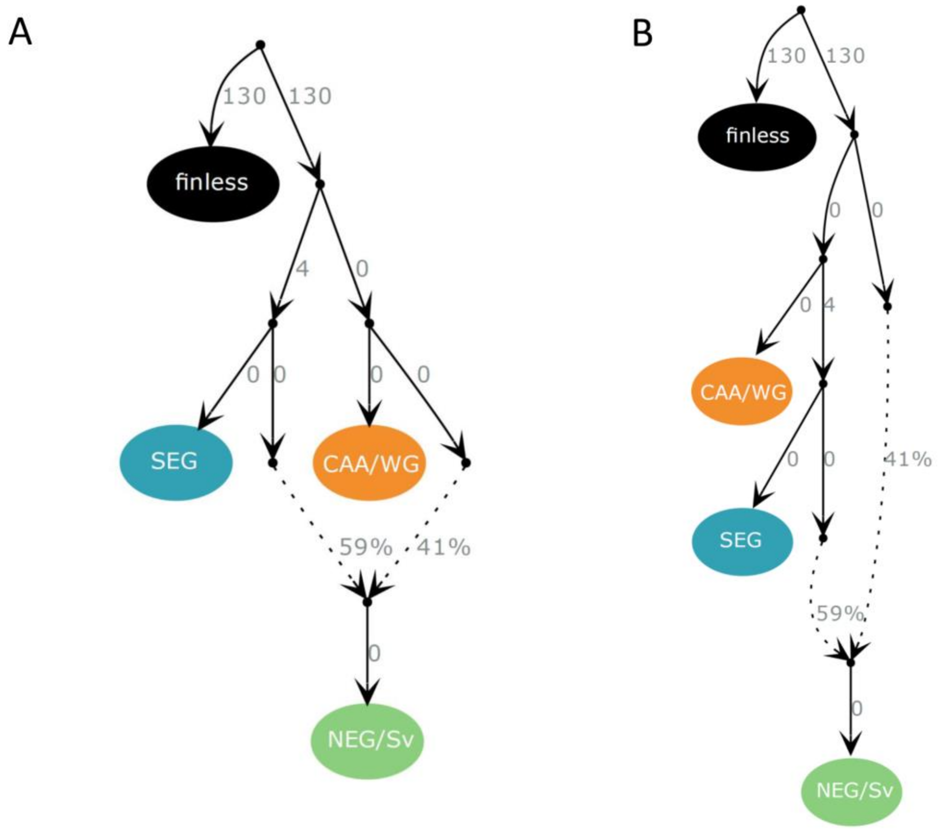


**Supplementary Figure S12**. **Evolutionary relationships between narwhal populations using the finless porpoise (finless) as the outgroup.** Admixture graphs were built using 823,928 SNPs. The graphs A) and B) were the two graphs out of 14 possible graph combinations presenting no outlier *f*-statistics (i.e. all |Z| were <3). The two models were equally likely after computing Bayes Factors. Continuous lines indicate phylogenetic relationships between populations/samples and the numbers at their right side the estimated genetic drift. Dotted lines show admixture edges and the number at their right side the percentage of ancestry deriving from each lineage.

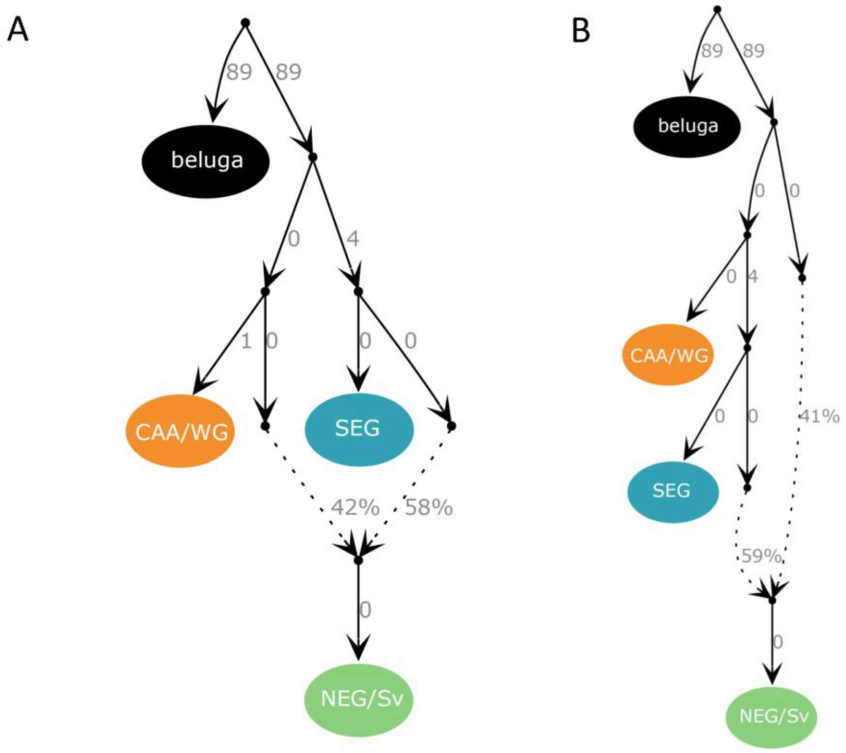


**Supplementary Figure S13**. **Evolutionary relationships between narwhal populations using the beluga as an outgroup.** Admixture graphs were built using 768,978 SNPs. The graphs A) and B) were the two graphs out of 14 possible graph combinations presenting no outlier *f*-statistics (i.e. all |Z| were <3). The two models were equally likely after computing Bayes Factors. Continuous lines indicate phylogenetic relationships between populations/samples and the numbers at their right side the estimated genetic drift. Dotted lines show admixture edges and the number at their right side the percentage of ancestry deriving from each lineage.

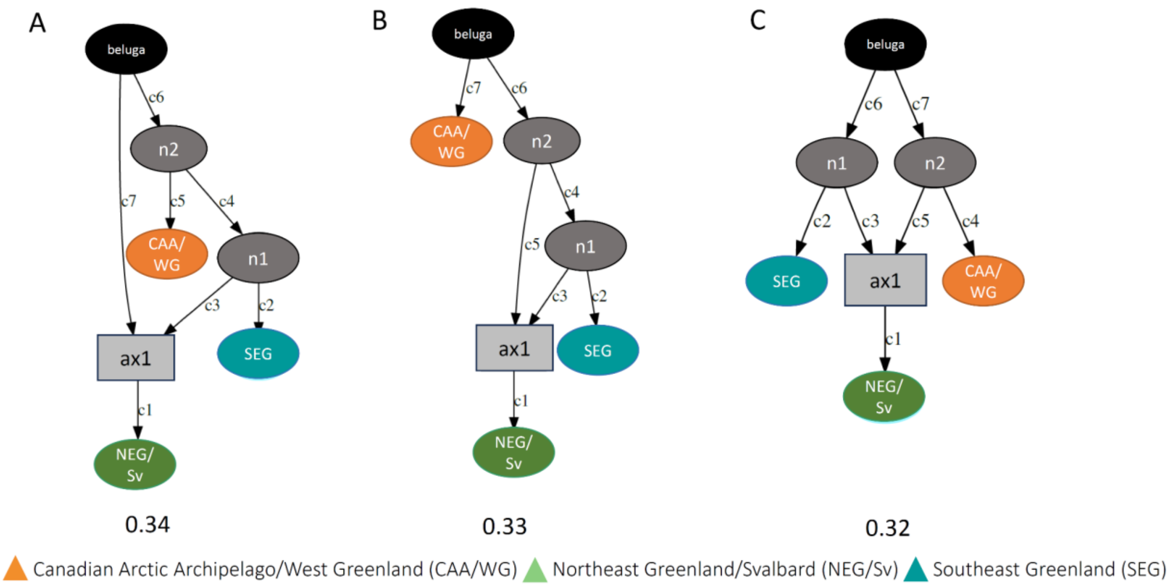


**Supplementary Figure S14**. **Evolutionary relationships between narwhal populations using Admixture Bayes.** There were three high-probability admixture graphs, the posterior probabilities are indicated below the graphs. The graph is rooted using the beluga, results were highly similar when using the finless porpoise (results not shown).

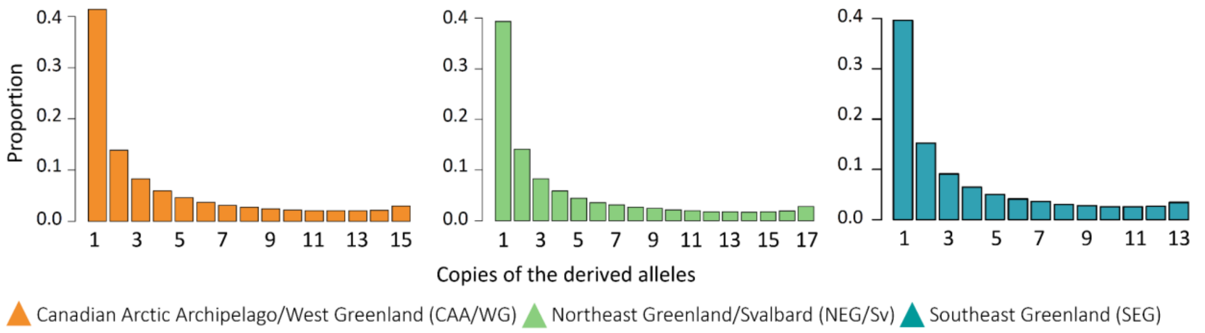


**Supplementary Figure S15. Unfolded 1D-site frequency spectrum (1D-SFS)** estimated using winsfs for each of the three narwhal populations.

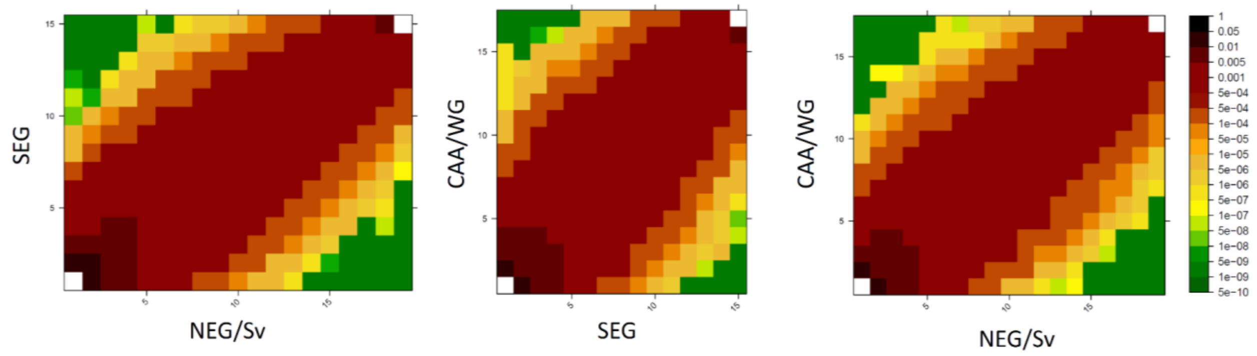


**Supplementary Figure S16. 2D-Site Frequency Spectrum (2D-SFS)** estimated using winsfs for each pair of populations, with Canadian Arctic Archipelago and West Greenland (CAA/WG); Northeast Greenland and Svalbard (NEG/Sv); and Southeast Greenland (SEG).

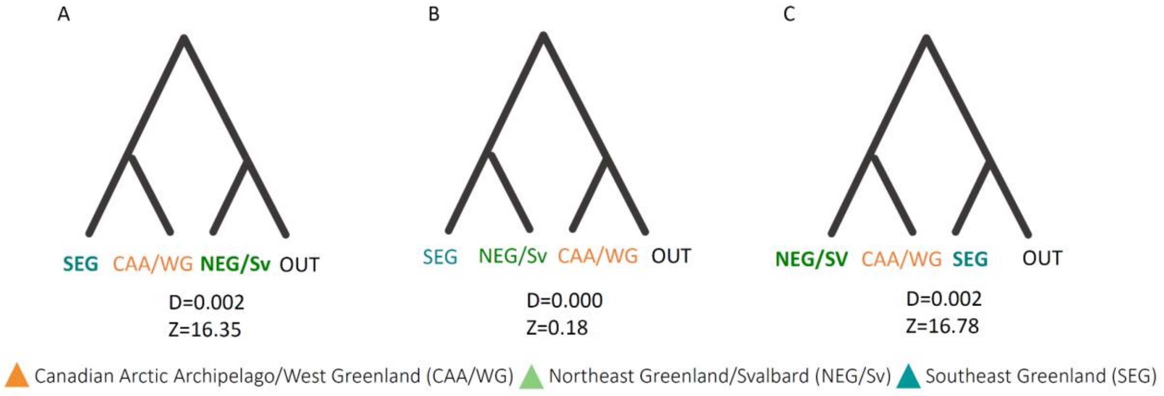


**Supplementary Figure S17. D-statistics of the three narwhal population**s of the form (SE, CAA/WG; NEG/Sv, OUT), (SE, NEG/Sv; CAA/WG, OUT) and (NEG/Sv, CAA/WG; SE, OUT) using the beluga as the outgroup (OUT). We get very similar results using the finless porpoise as the outgroup. It is important to note that the D-statistics were calculated using the following formula: D = (nBABA-nABBA) / (nBABA+nABBA). Populations in bold indicate which populations share an excess of derived alleles, when the D-statistics are significant.

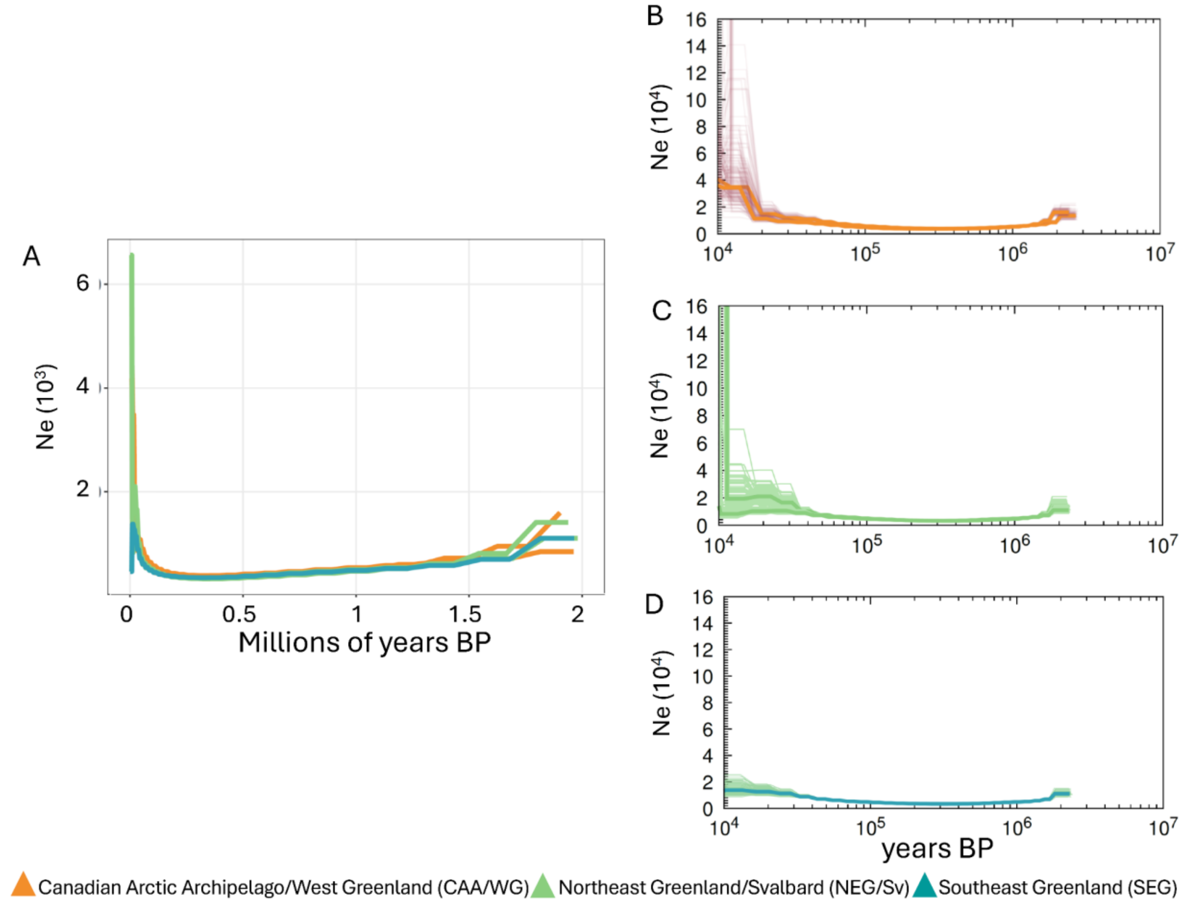


**Supplementary Figure 18. Demographic reconstruction based on individual genomes** using pairwise sequentially Markovian coalescent (PSMC) with a mutation rate of 1.56e-08 per generation (Westbury et al. 2019) and a generation time of 30 years (Garde et al. 2007) for two individuals from CAA/WG; two for NEG/Sv; and one for SEG showing A) the main curve only. B-D) Region-based PSMC for the main curve (bold) and 100 bootstraps replicates (B-D) for B) the CAA/WG individuals, C) the NEG individuals and D) the SEG individuals.

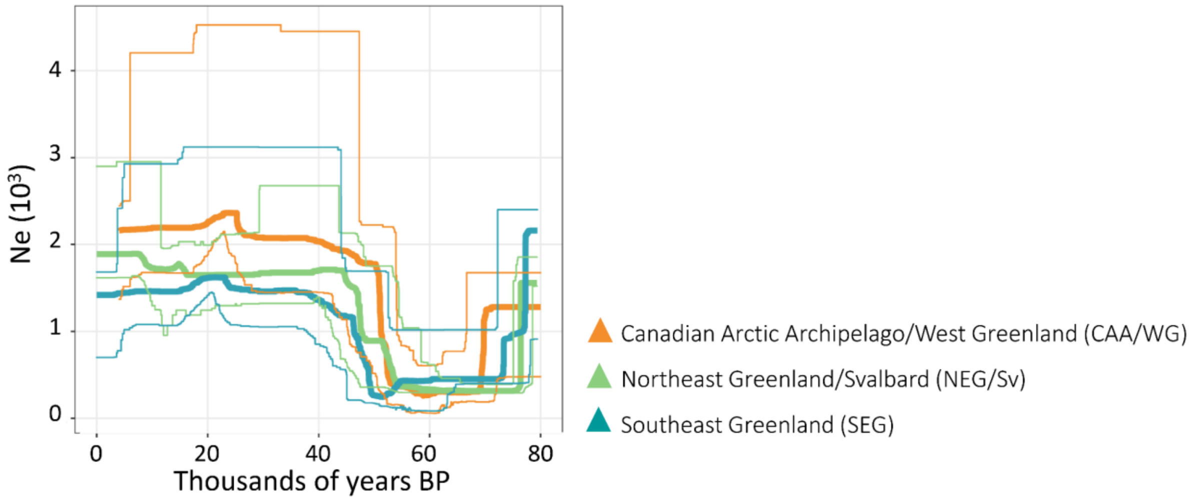


**Supplementary Figure S19. Demographic reconstruction based on population data using stairwayplot.** Changes in effective population size ($N_e$) through time inferred for high coverage narwhal individuals: two for CAA/WG; two for NEG/Sv; and one for SEG using pairwise sequentially Markovian coalescent (PSMC) using a mutation rate of 1.56e-08 (add metrics) (Westbury et al. 2019) and a generation time of 30 years (Garde et al. 2007); for the main curve (bold) and 100 bootstraps replicates (C-E) for C) the CAA/WG individuals, D, the NEG individuals and E) the SEG individuals.

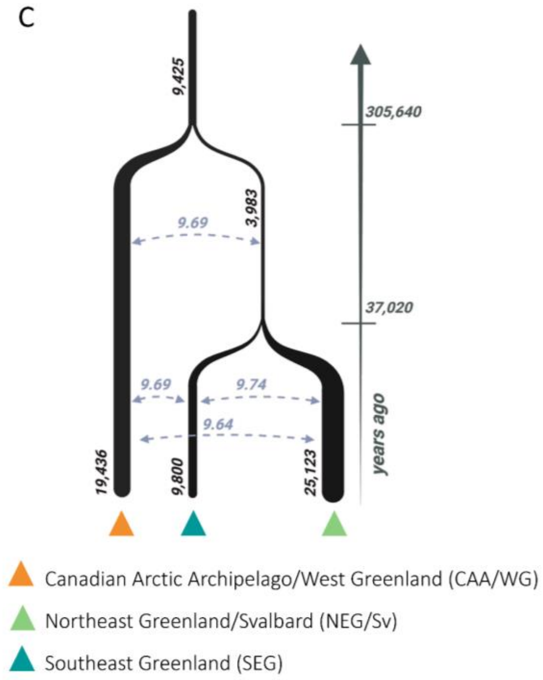


**Supplementary Figure S20. Demographic history of the three narwhal populations inferred using fastsimcoal2.** Maximum likelihood parameter estimates for ancestral and contemporary effective population sizes ($N_e$), divergence times, and migration rates between each pair of populations. Effective population sizes are given in numbers of haploid individuals; migration rates are expressed in units of *Nm* (the number of migrants per generation).

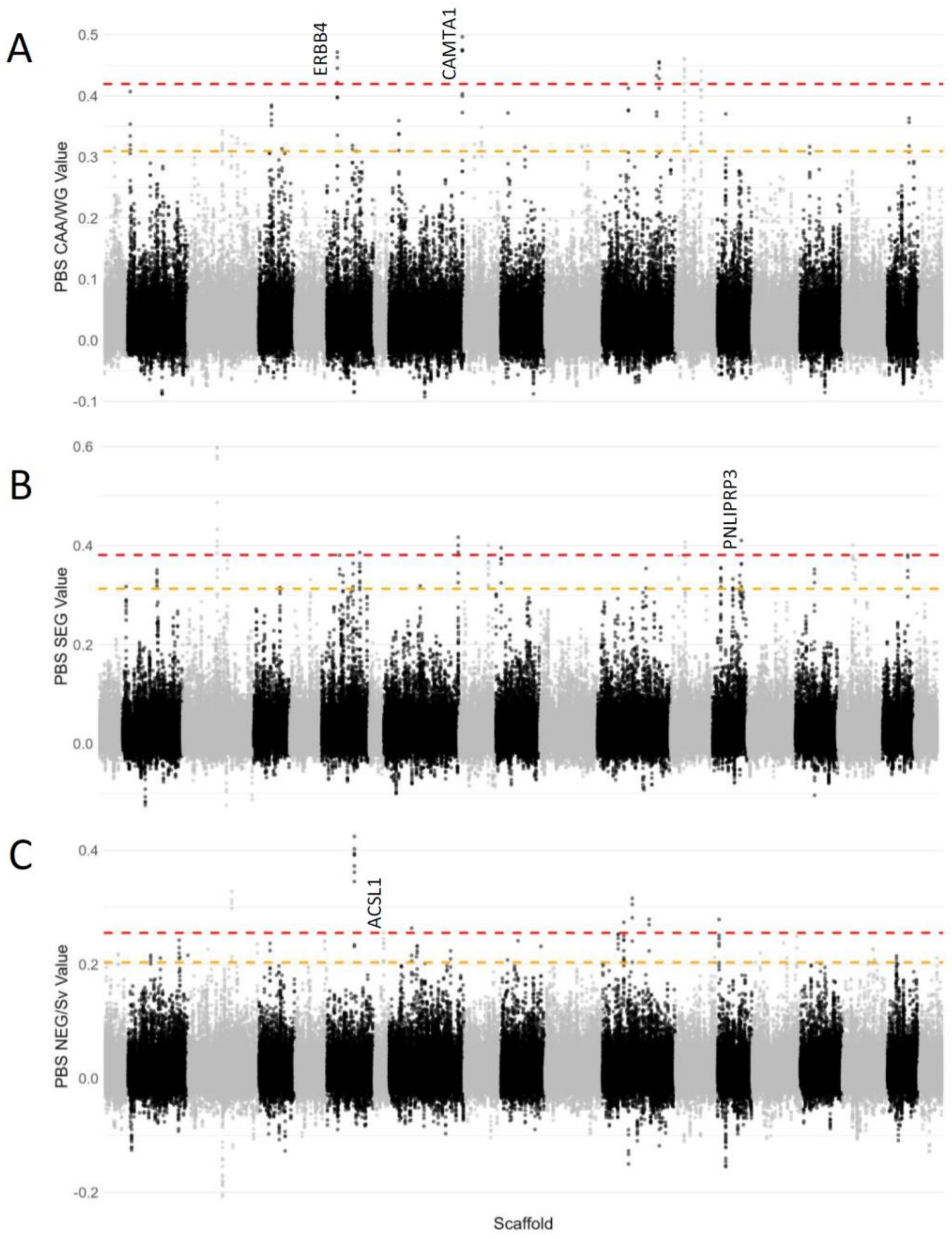


**Supplementary Figure 21. Signatures of selection using PBS**. Manhattan plot of the PBS values in windows of 50 k base pairs (kb) using a 10-kb step across large autosomal scaffolds - different scaffolds are colored in successions of black and grey for each population: A) Canadian Arctic Archipelago and West Greenland (CAA/WG); B) Southeast Greenland (SEG) and C) Northeast Greenland and Svalbard (NEG/Sv). The dashed orange and red lines indicate the 99.95th and 99.99th percentiles, respectively, above which we consider the windows as outliers. The 99.99th outliers gene linked to long-term memory (ERBA4 and

CAMTA1 in CAA/WG) and feeding ecology (ACSL1 in NEG/Sv and PNLIPRP3 in SEG) are highlighted

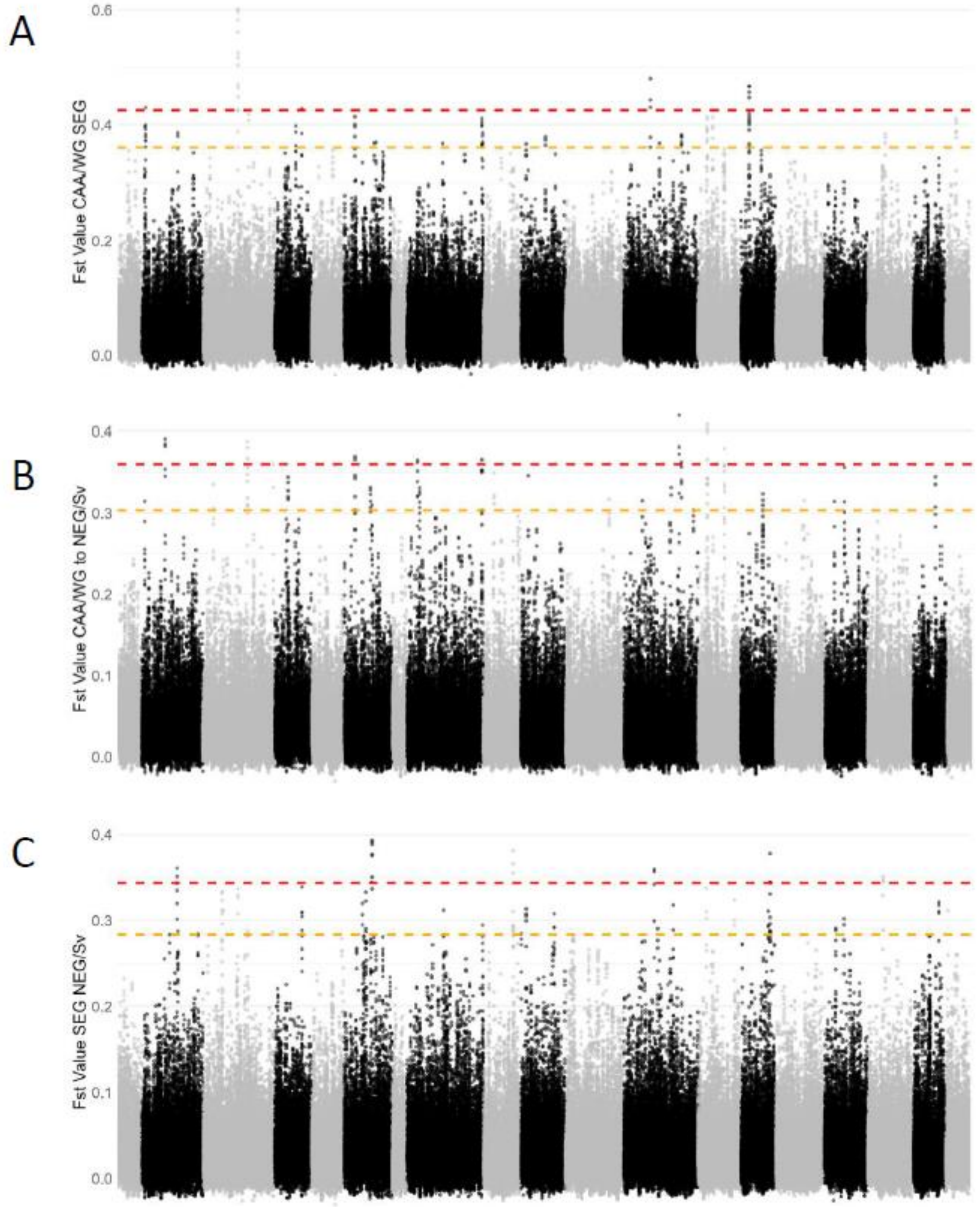


**Supplementary Figure 22. Pairwise $F_{ST}$ values**. Manhattan plot of the pairwise $F_{ST}$ values in windows of 50 k base pairs (kb) using a 10-kb step across large autosomal scaffolds - different scaffolds are colored in successions of black and grey for each A) Canadian Arctic Archipelago and West Greenland (CAA/WG) and Southeast Greenland (SEG) B) CAA/WG and Northeast Greenland and Svalbard (NEG/Sv); and C) SEG and NEG/Sv). The dashed orange and red lines indicate the 99.95th and 99.99th percentiles, respectively.

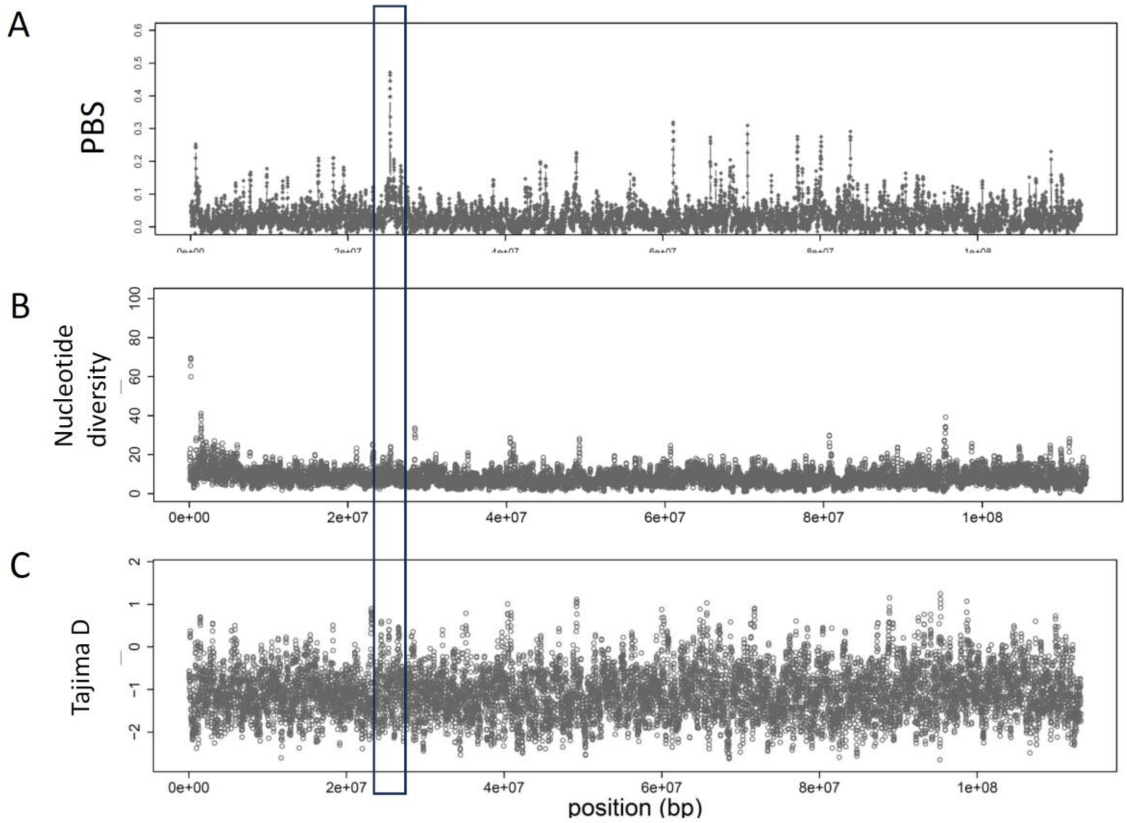


**Supplementary Figure 23. Diversity patterns around gene ERBBA.** PBS, nucleotide diversity and Tajima D values for scaffold NW_021703772.1 where the ERBBA gene is found (highlighted in the box) for Canadian Arctic Archipelago and West Greenland (CAA/WG) narwhals.

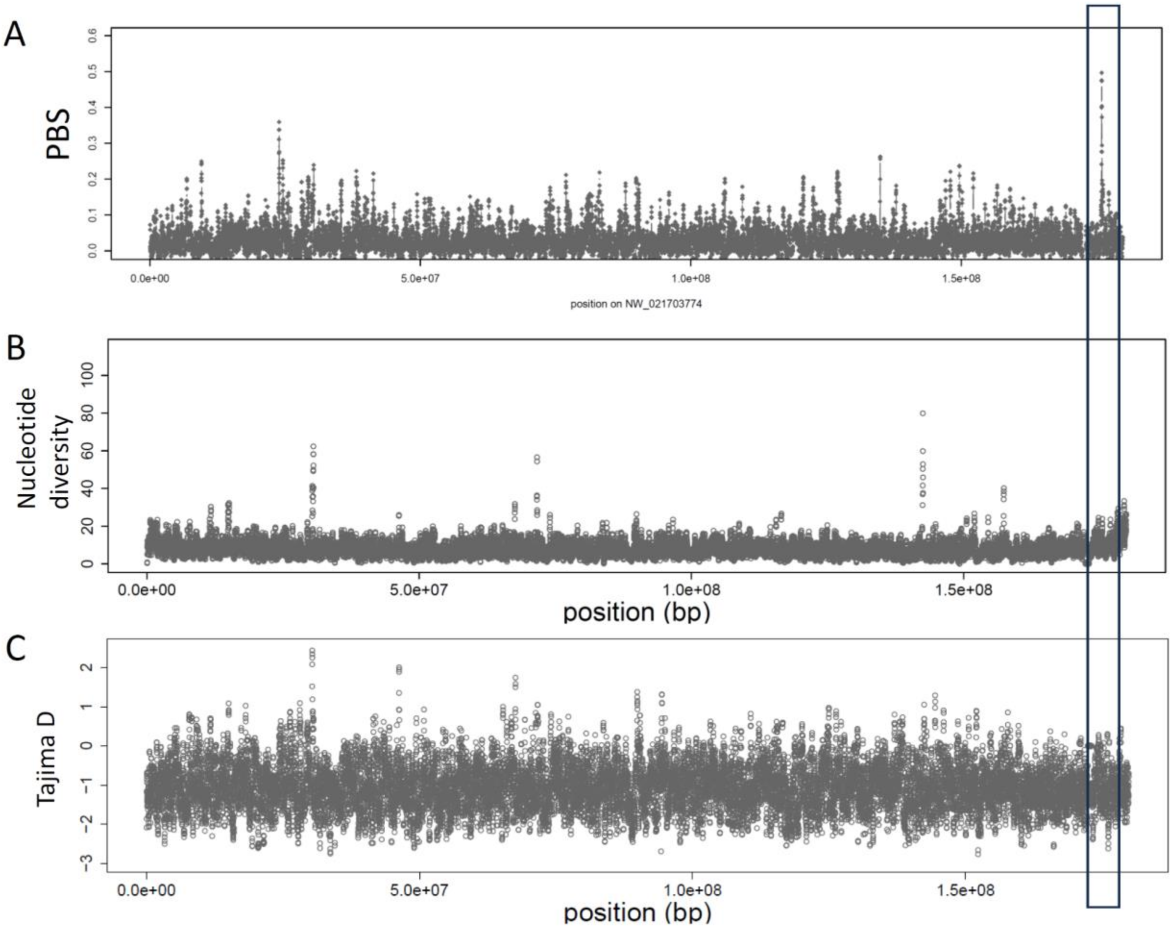


**Supplementary Figure 24. Diversity patterns around gene CAMTA1.** PBS, nucleotide diversity and Tajima D values for scaffold NW_021703774.1 where the CAMTA1 gene is found (highlighted in the box) for Canadian Arctic Archipelago and West Greenland (CAA/WG) narwhals.

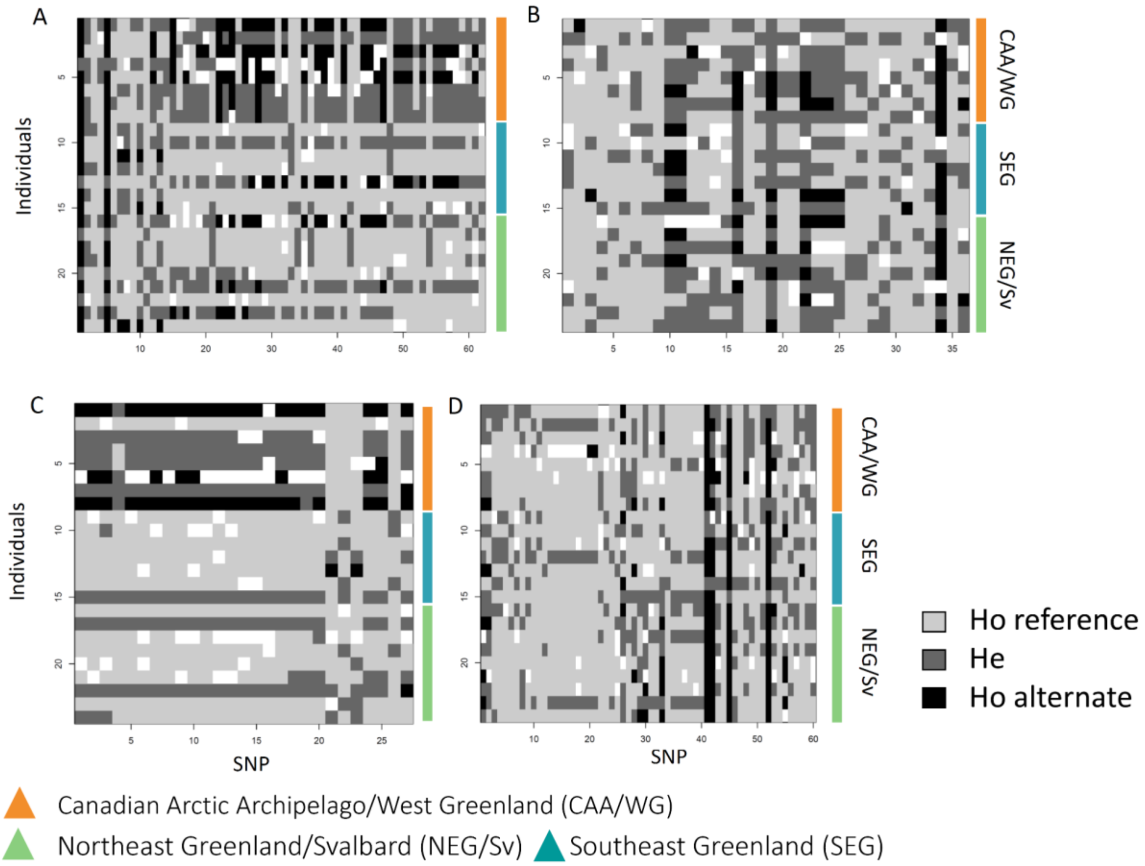


**Supplementary Figure 25. Genotypes for the genes identified as outliers in Canadian Arctic Archipelago and West Greenland (CAA/WG) narwhals** in the PBS analyses (in the 99.99th percentiles) and involved in long-term memory and a random sequence on the rest of the scaffold a) ERBB4 and b) a random sequence on the same scaffold, and c) CAMTA1 and d) a random sequence on the rest of the scaffold. The two other populations are Southeast Greenland (SEG) and Northeast Greenland and Svalbard (NEG/Sv).

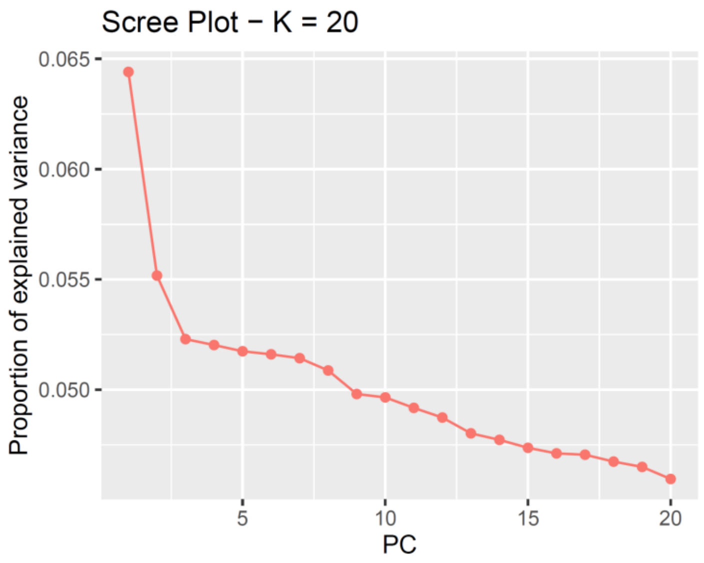

**Supplementary Figure 26. Screeplot for the PCAdapt analysis** showing the percentage of variance explained by each Principal Component (PC). In an ideal scenario , the eigenvalues related to random variation are lying on a straight line while the ones related to population structure are lying on a steep curve. It is recommended to follow Cattell's rule and keep a number of PC (K) corresponding to eigenvalues on the left of the straight line. Here, although we do not observe a totally straight line, the plot suggests 2 is the optimal number of PCs to consider.

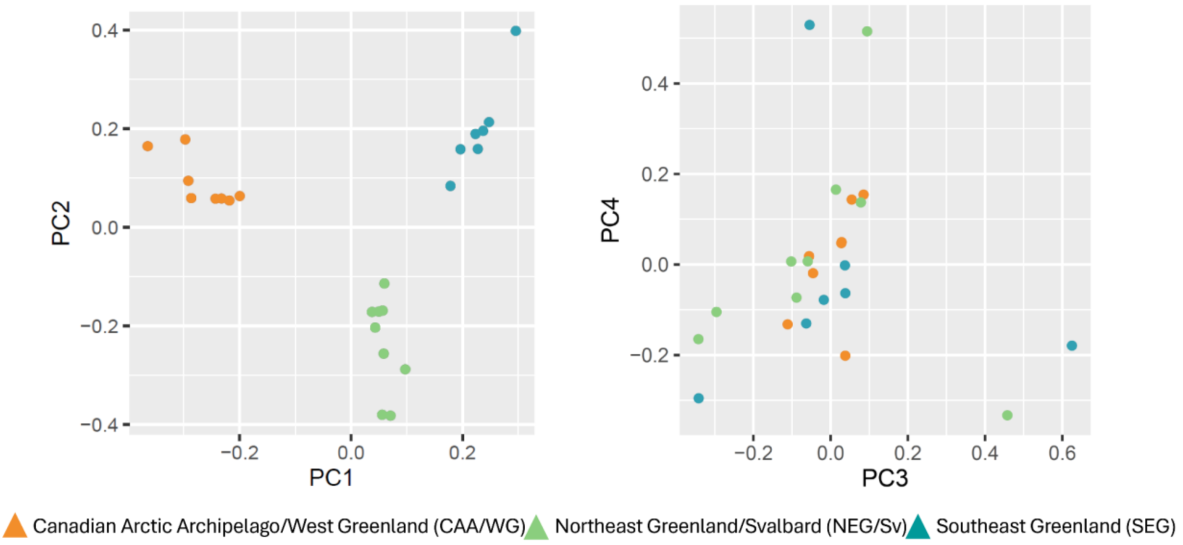


**Supplementary Figure 27. Score plot for the PCAdapt analysis**, displaying the projections of the individuals onto a) Principal Components (PCs) 1 and 2 and b) PCs 3 and 4. Individuals are colored according to their population of origin. The choice of the number of PCs (*K*) to consider can be limited to those reflecting relevant levels of population structure, in our case 1 and 2.

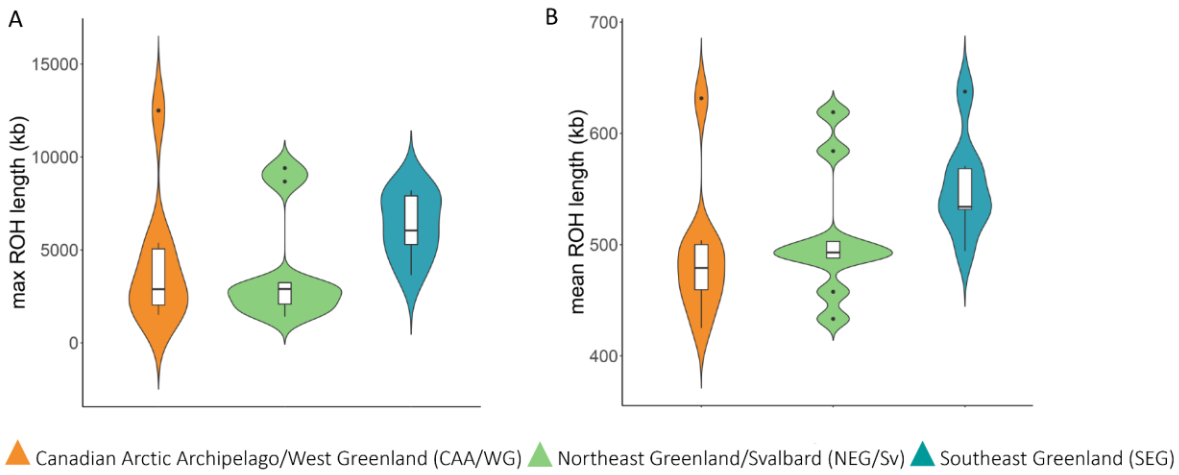


**Supplementary Figure 28. Levels of inbreeding in the three narwhal populations.** A) maximum length of Runs of homozygosity (ROH) and mean length of ROH in the three populations.

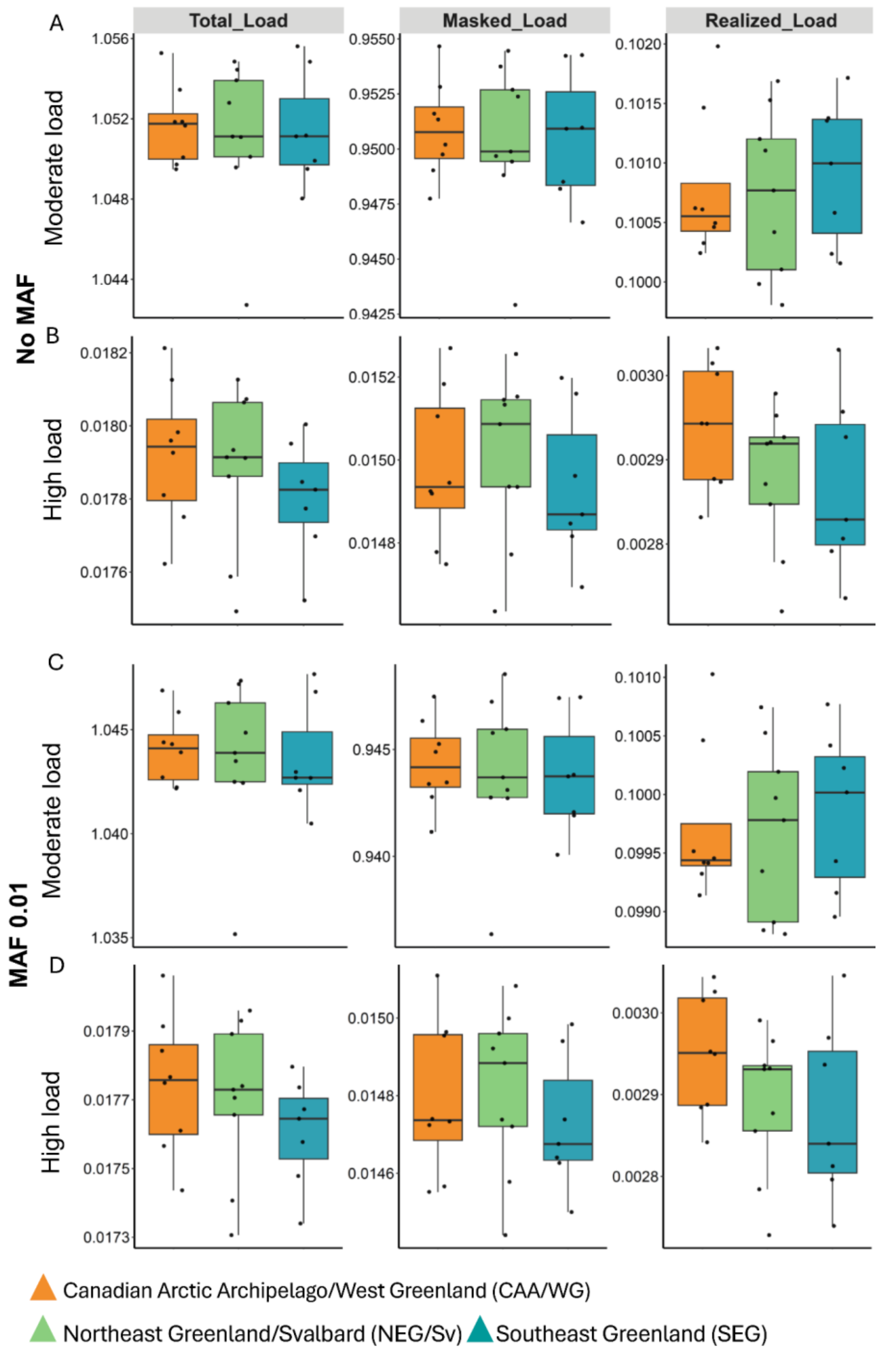


**Supplementary Figure 29. Mutation load statistics.** Genetic load is calculated as the ratio between derived allele counts in moderate (A,C) or high (B,D) impact mutations by the derived allele count in synonymous mutations for each population with no MAF filter (A-B) and a MAF filter of 0.01 (C,D). Total load represents the global count of derived alleles in deleterious positions, masked load the count in heterozygous state and realized load the count in homozygous state.

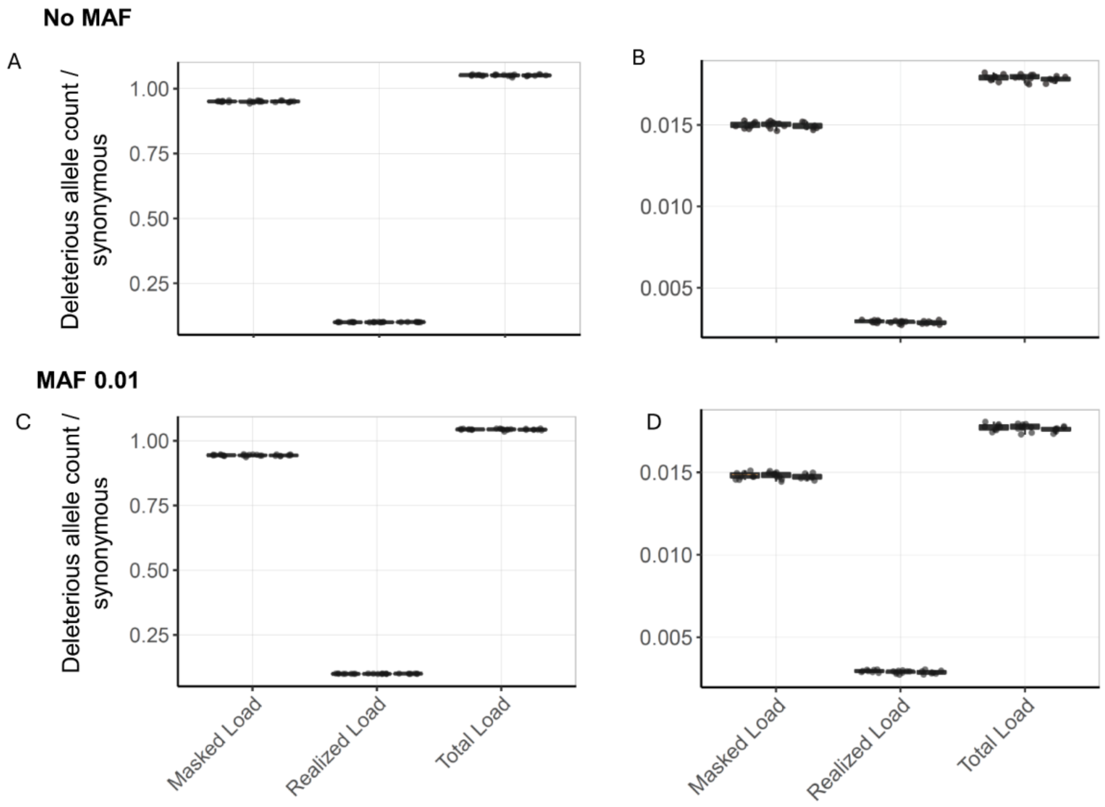


**Supplementary Figure 30. Mutation load statistics summarised by deleterious levels.** Genetic load is calculated as the ratio between derived allele counts in moderate (A,C) and high (B,D) impact mutations by the derived allele count in synonymous mutations for each population with no MAF filter (A,B) and a MAF filter of 0.01 (C,D), in the following order on the figures: Canadian Arctic Archipelago and West Greenland (CAA/WG); Northeast Greenland and Svalbard (NEG/Sv); and Southeast Greenland (SEG). Masked load represents the count in heterozygous state, realized load represents the count in homozygous state, and total load represents the global count of derived alleles in deleterious positions.

### Supplementary Tables

**Supplementary Table S1. Overview of the sample and data.** CGG_ID refer to the Centre for GeoGenetics database sample ID at the University of Copenhagen, Lab_ID to the ID in the laboratory while sample_ID refers to the sample ID in each institution; stock the narwhal stock from which the narwhal is from; locality the location where the sample was collected either from subsistence hunt or biopsy sampling; the sampling date; SRA the Genbank accession number; library built method; PE-SE whether the library built is paired-end or single-end; coverage of the data; platform - data used the sequencing platforms used for the analyses; platform SRA is the sequencing platforms used and for which data is available on SRA.

| CGG_I+1:114D | Lab_ID | sample ID | stock | locality | Date | sex | Biosample AN | library built | PE-SE | coverage | platfom - data used | platfom SRA |
|---|---|---|---|---|---|---|---|---|---|---|---|---|
| CGG_1_017220 | CGG_1_017220 | BI95-328 | Eastern_Baffin_Island | Broughton Island | 30/7/1995 | F | SAMN60394159 | Illumina | SE | 0.23 | Hiseq 2500 | Hiseq 2500 |
| CGG_1_017221 | CGG_1_017221 | BI04-1085 | Eastern_Baffin_Island | Broughton Island | 10/8/2004 | M | SAMN60394160 | Illumina | SE | 0.23 | Hiseq 2500 | Hiseq 2500 |
| CGG_1_017222 | CGG_1_017222 | BI04-1086 | Eastern_Baffin_Island | Broughton Island | 10/7/2004 | M | SAMN60394161 | Illumina | SE | 0.26 | Hiseq 2500 | Hiseq 2500 |
| CGG_1_017225 | CGG_1_017225 | BI08-1118 | Eastern_Baffin_Island | Broughton Island | 2007 | F | SAMN60394162 | Illumina | SE | 0.3 | Hiseq 2500 | Hiseq 2500 |
| CGG_1_017226 | CGG_1_017226 | CR94-203 | Eastern_Baffin_Island | Clyde River | 31/7/1994 | M | SAMN60394163 | Illumina | SE | 0.23 | Hiseq 2500 | Hiseq 2500 |
| CGG_1_017227 | CGG_1_017227 | CR94-206 | Eastern_Baffin_Island | Clyde River | 1/7/1994 | M | SAMN60394164 | Illumina | SE | 0.28 | Hiseq 2500 | Hiseq 2500 |
| CGG_1_017228 | CGG_1_017228 | CR95-361 | Eastern_Baffin_Island | Clyde River | 6/8/1995 | F | SAMN60394165 | Illumina | SE | 0.54 | Hiseq 2500 | Hiseq 2500 |
| CGG_1_017229 | CGG_1_017229 | CR04-1010 | Eastern_Baffin_Island | Clyde River | 2004 | M | SAMN60394166 | Illumina | SE | 0.27 | Hiseq 2500 | Hiseq 2500 |
| CGG_1_017232 | CGG_1_017232 | CR04-1017 | Eastern_Baffin_Island | Clyde River | 2004 | M | SAMN60394167 | Illumina | SE | 0.47 | Hiseq 2500 | Hiseq 2500 |
| CGG_1_017233 | CGG_1_017233 | CR05-1032 | Eastern_Baffin_Island | Clyde River | 2005 | M | SAMN60394168 | Illumina | SE | 0.2 | Hiseq 2500 | Hiseq 2500 |
| CGG_1_017235 | CGG_1_017235 | CB00-7929 | Somerset_Island | Creswell Bay | 24/8/2000 | F | SAMN60394169 | Illumina | SE | 0.2 | Hiseq 2500 | Hiseq 2500 |
| CGG_1_017237 | CGG_1_017237 | CB01-20692 | Somerset_Island | Creswell Bay | 9/8/2001 | M | SAMN60394170 | Illumina | SE | 0.19 | Hiseq 2500 | Hiseq 2500 |
| CGG_1_017238 | CGG_1_017238 | CB01-3960 | Somerset_Island | Creswell Bay | 9/8/2001 | F | SAMN60394171 | Illumina | SE | 0.33 | Hiseq 2500 | Hiseq 2500 |
| CGG_1_017239 | CGG_1_017239 | CB01-7927 | Somerset_Island | Creswell Bay | 7/8/2001 | F | SAMN60394172 | Illumina | SE | 0.22 | Hiseq 2500 | Hiseq 2500 |
| CGG_1_017240 | CGG_1_017240 | CB01-7928 | Somerset_Island | Creswell Bay | 8/8/2001 | M | SAMN60394173 | Illumina | SE | 0.3 | Hiseq 2500 | Hiseq 2500 |
| CGG_1_017244 | CGG_1_017244 | GF99-1017 | Jones_Sound | Grise Fiord | 16/8/1999 | M | SAMN60394174 | Illumina | SE | 0.24 | Hiseq 2500 | Hiseq 2500 |

| | | | | | | | | | | | | |
|---|---|---|---|---|---|---|---|---|---|---|---|---|
| CGG_1_017245 | CGG_1_017245 | GF99-1022 | Jones_Sound | Grise Fiord | 16/8/1999 | M | SAMN60394175 | Illumina | SE | 0.22 | Hiseq 2500 | Hiseq 2500 |
| CGG_1_017246 | CGG_1_017246 | GF03-1095 | Jones_Sound | Grise Fiord | 9/8/2003 | M | SAMN60394176 | Illumina | SE | 0.2 | Hiseq 2500 | Hiseq 2500 |
| CGG_1_017247 | CGG_1_017247 | GF03-1107 | Jones_Sound | Grise Fiord | 10/8/2003 | M | SAMN60394177 | Illumina | SE | 0.26 | Hiseq 2500 | Hiseq 2500 |
| CGG_1_017248 | CGG_1_017248 | GF07-1124 | Jones_Sound | Grise Fiord | 27/7/2007 | M | SAMN60394178 | Illumina | SE | 0.17 | Hiseq 2500 | Hiseq 2500 |
| CGG_1_017249 | CGG_1_017249 | GF07-1129 | Jones_Sound | Grise Fiord | 31/7/2007 | M | SAMN60394179 | Illumina | SE | 0.31 | Hiseq 2500 | Hiseq 2500 |
| CGG_1_017253 | CGG_1_017253 | IG95-462 | Somerset_Island | Igloolik | 4/8/1995 | F | SAMN60394180 | Illumina, BEST | SE, PE | 0.26 | Hiseq 2500, Hiseq Xten | Hiseq 2500, Hiseq Xten |
| CGG_1_017254 | CGG_1_017254 | IG95-465 | Somerset_Island | Igloolik | 4/8/1995 | F | SAMN60394181 | Illumina | SE | 0.23 | Hiseq 2500 | Hiseq 2500 |
| CGG_1_017256 | CGG_1_017256 | IG95-481 | Somerset_Island | Igloolik | 4/8/1995 | F | SAMN60394182 | Illumina | SE | 0.29 | Hiseq 2500 | Hiseq 2500 |
| CGG_1_017257 | CGG_1_017257 | IG06-1222 | Somerset_Island | Igloolik | 2006 | F | SAMN60394183 | Illumina | SE | 0.24 | Hiseq 2500 | Hiseq 2500 |
| CGG_1_017258 | CGG_1_017258 | IG06-1231 | Somerset_Island | Igloolik | 2006 | M | SAMN60394184 | Illumina | SE | 0.23 | Hiseq 2500 | Hiseq 2500 |
| CGG_1_017259 | CGG_1_017259 | IG06-1238 | Somerset_Island | Igloolik | 2006 | M | SAMN60394185 | Illumina | SE | 0.23 | Hiseq 2500 | Hiseq 2500 |
| CGG_1_017260 | CGG_1_017260 | IG06-1242 | Somerset_Island | Igloolik | 2006 | M | SAMN60394186 | Illumina | SE | 0.28 | Hiseq 2500 | Hiseq 2500 |
| CGG_1_017262 | CGG_1_017262 | PG95-548 | Eastern_Baffin_Island | Pangnirtung | 27/7/1996 | M | SAMN60394187 | Illumina | SE | 0.19 | Hiseq 2500 | Hiseq 2500 |
| CGG_1_017263 | CGG_1_017263 | PG93-170 | Eastern_Baffin_Island | Pangnirtung | 27/7/1996 | M | SAMN60394188 | Illumina | SE | 0.44 | Hiseq 2500 | Hiseq 2500 |
| CGG_1_017264 | CGG_1_017264 | PG96-151 | Eastern_Baffin_Island | Pangnirtung | 25/8/1996 | F | SAMN60394189 | Illumina | SE | 0.25 | Hiseq 2500 | Hiseq 2500 |
| CGG_1_017265 | CGG_1_017265 | PG04-1032 | Eastern_Baffin_Island | Pangnirtung | 7/7/2004 | M | SAMN60394190 | Illumina | SE | 0.36 | Hiseq 2500 | Hiseq 2500 |
| CGG_1_017266 | CGG_1_017266 | PG04-1179 | Eastern_Baffin_Island | Pangnirtung | 7/7/2004 | M | SAMN60394191 | Illumina | SE | 0.27 | Hiseq 2500 | Hiseq 2500 |
| CGG_1_017267 | CGG_1_017267 | PG04-1183 | Eastern_Baffin_Island | Pangnirtung | 7/7/2004 | M | SAMN60394192 | Illumina | SE | 0.28 | Hiseq 2500 | Hiseq 2500 |
| CGG_1_017268 | CGG_1_017268 | PI82-19 | Eclipse_Sound | Pond Inlet | 8/8/1982 | F | SAMN60394193 | Illumina | SE | 0.25 | Hiseq 2500 | Hiseq 2500 |
| CGG_1_017272 | CGG_1_017272 | PI10-1323 | Eclipse_Sound | Pond Inlet | 22/8/2010 | F | SAMN60394194 | Illumina | SE | 0.37 | Hiseq 2500 | Hiseq 2500 |
| CGG_1_017273 | CGG_1_017273 | PI11-001 | Eclipse_Sound | Pond Inlet | 18/8/2011 | F | SAMN60394195 | Illumina | SE | 0.25 | Hiseq 2500 | Hiseq 2500 |
| CGG_1_017274 | CGG_1_017274 | PI11-003 | Eclipse_Sound | Pond Inlet | 19/8/2011 | F | SAMN60394196 | Illumina | SE | 0.36 | Hiseq 2500 | Hiseq 2500 |
| CGG_1_017275 | CGG_1_017275 | PI11-006 | Eclipse_Sound | Pond Inlet | 16/8/2011 | F | SAMN60394197 | Illumina | SE | 0.35 | Hiseq 2500 | Hiseq 2500 |
| CGG_1_017276 | CGG_1_017276 | PI12-001 | Eclipse_Sound | Pond Inlet | 13/8/2012 | F | SAMN60394198 | Illumina | SE | 0.51 | Hiseq 2500 | Hiseq 2500 |
| CGG_1_017277 | CGG_1_017277 | PI12-003 | Eclipse_Sound | Pond Inlet | 14/8/2012 | M | SAMN60394199 | Illumina | SE | 0.3 | Hiseq 2500 | Hiseq 2500 |
| CGG_1_017280 | CGG_1_017280 | RB95-146 | Northern_Hudson_Bay | Repulse Bay | 17/8/1995 | F | SAMN60394200 | Illumina | SE | 0.3 | Hiseq 2500 | Hiseq 2500 |
| CGG_1_017282 | CGG_1_017282 | RB00-1092 | Northern_Hudson_Bay | Repulse Bay | 21/7/2000 | M | SAMN60394201 | Illumina | SE | 0.31 | Hiseq 2500 | Hiseq 2500 |
| CGG_1_017283 | CGG_1_017283 | RB00-1094 | Northern_Hudson_Bay | Repulse Bay | 24/7/2000 | M | SAMN60394202 | Illumina | SE | 0.36 | Hiseq 2500 | Hiseq 2500 |
| CGG_1_017286 | CGG_1_017286 | RB06-1231 | Northern_Hudson_Bay | Repulse Bay | 15/7/2006 | F | SAMN60394203 | Illumina | SE | 0.44 | Hiseq 2500 | Hiseq 2500 |
| CGG_1_017287 | CGG_1_017287 | RB06-1251 | Northern_Hudson_Bay | Repulse Bay | 15/7/2006 | M | SAMN60394204 | Illumina | SE | 0.27 | Hiseq 2500 | Hiseq 2500 |
| CGG_1_017288 | CGG_1_017288 | RB07-001 | Northern_Hudson_Bay | Repulse Bay | 2007 | F | SAMN60394205 | Illumina | SE | 0.38 | Hiseq 2500 | Hiseq 2500 |
| CGG_1_017289 | CGG_1_017289 | RB07-002A | Northern_Hudson_Bay | Repulse Bay | 2007 | M | SAMN60394206 | Illumina | SE | 0.41 | Hiseq 2500 | Hiseq 2500 |
| CGG_1_017295 | CGG_1_017295 | RE02-1094 | Somerset_Island | Resolute Bay | 14/8/2002 | M | SAMN60394207 | Illumina | SE | 0.27 | Hiseq 2500 | Hiseq 2500 |
| CGG_1_017296 | CGG_1_017296 | RE02-1106 | Somerset_Island | Resolute Bay | 14/8/2002 | M | SAMN60394208 | Illumina | SE | 0.31 | Hiseq 2500 | Hiseq 2500 |
| CGG_1_017297 | CGG_1_017297 | RE06-1158 | Somerset_Island | Resolute Bay | 1/9/2006 | M | SAMN60394209 | Illumina | SE | 0.29 | Hiseq 2500 | Hiseq 2500 |
| CGG_1_017298 | CGG_1_017298 | RE02-1100 | Somerset_Island | Resolute Bay | 14/8/2002 | M | SAMN60394210 | Illumina | SE | 0.36 | Hiseq 2500 | Hiseq 2500 |
| CGG_1_017302 | CGG_1_017302 | SB96-398 | Somerset_Island | Taloyoak | 20/8/1996 | F | SAMN60394211 | Illumina | SE | 0.32 | Hiseq 2500 | Hiseq 2500 |
| CGG_1_017303 | CGG_1_017303 | SB09-1050 | Somerset_Island | Taloyoak | 29/8/2009 | F | SAMN60394212 | Illumina | SE | 0.35 | Hiseq 2500 | Hiseq 2500 |
| CGG_1_017530 | CGG_1_017530 | 579 | Eclipse_Sound | Tremblay Sound | 19/8/1997 | M | SAMN60394213 | Illumina | SE | 0.31 | Hiseq 2500 | Hiseq 2500 |

| | | | | | | | | | | | | |
|---|---|---|---|---|---|---|---|---|---|---|---|---|
| CGG_1_017532 | CGG_1_017532 | 591 | Eclipse_Sound | Tremblay Sound | 24/8/1997 | M | SAMN60394214 | Illumina | SE | 0.4 | Hiseq 2500 | Hiseq 2500 |
| CGG_1_017534 | CGG_1_017534 | 829 | Eclipse_Sound | Tremblay Sound | 15/8/1999 | F | SAMN60394215 | Illumina | SE | 0.28 | Hiseq 2500 | Hiseq 2500 |
| CGG_1_017536 | CGG_1_017536 | 831 | Eclipse_Sound | Tremblay Sound | 16/8/1999 | F | SAMN60394216 | Illumina | SE | 0.19 | Hiseq 2500 | Hiseq 2500 |
| CGG_1_017537 | CGG_1_017537 | 832 | Eclipse_Sound | Tremblay Sound | 15/8/1999 | F | SAMN60394217 | Illumina | SE | 0.17 | Hiseq 2500 | Hiseq 2500 |
| CGG_1_017542 | CGG_1_017542 | 841 | Melville_Bay | Melville Bugten | 10/9/2006 | M | SAMN60394218 | Illumina | SE | 0.2 | Hiseq 2500 | Hiseq 2500 |
| CGG_1_017543 | CGG_1_017543 | 842 | Melville_Bay | Melville Bugten | 11/9/2006 | F | SAMN60394219 | Illumina | SE | 0.15 | Hiseq 2500 | Hiseq 2500 |
| CGG_1_017544 | CGG_1_017544 | 843 | Melville_Bay | Melville Bugten | 11/9/2006 | M | SAMN60394220 | Illumina | SE | 0.78 | Hiseq 2500 | Hiseq 2500 |
| CGG_1_017545 | CGG_1_017545 | 844 | Melville_Bay | Melville Bugten | 11/9/2006 | F | SAMN60394221 | Illumina | SE | 0.56 | Hiseq 2500 | Hiseq 2500 |
| CGG_1_017546 | CGG_1_017546 | 845 | Melville_Bay | Melville Bugten | 11/9/2006 | F | SAMN60394222 | Illumina | SE | 0.24 | Hiseq 2500 | Hiseq 2500 |
| CGG_1_017547 | CGG_1_017547 | 846 | Melville_Bay | Melville Bugten | 11/9/2006 | F | SAMN60394223 | Illumina | SE | 0.18 | Hiseq 2500 | Hiseq 2500 |
| CGG_1_017548 | CGG_1_017548 | 847 | Melville_Bay | Melville Bugten | 11/9/2006 | M | SAMN60394224 | Illumina | SE | 0.39 | Hiseq 2500 | Hiseq 2500 |
| CGG_1_017551 | CGG_1_017551 | 871 | Melville_Bay | Melville Bugten | 4/9/2007 | F | SAMN60394225 | Illumina | SE | 0.3 | Hiseq 2500 | Hiseq 2500 |
| CGG_1_017566 | CGG_1_017566 | 1680 | Inglefield_Bredning | Qaanaaq | /8/1993 | F | SAMN60394226 | Illumina | SE | 0.27 | Hiseq 2500 | Hiseq 2500 |
| CGG_1_017567 | CGG_1_017567 | 1681 | Inglefield_Bredning | Qaanaaq | /8/1993 | M | SAMN60394227 | Illumina | SE | 0.35 | Hiseq 2500 | Hiseq 2500 |
| CGG_1_017569 | CGG_1_017569 | 1683 | Inglefield_Bredning | Qaanaaq | /8/1993 | F | SAMN60394228 | Illumina | SE | 0.22 | Hiseq 2500 | Hiseq 2500 |
| CGG_1_017571 | CGG_1_017571 | 1685 | Inglefield_Bredning | Qaanaaq | /8/1993 | M | SAMN60394229 | Illumina | SE | 0.22 | Hiseq 2500 | Hiseq 2500 |
| CGG_1_017572 | CGG_1_017572 | 1686 | Inglefield_Bredning | Qaanaaq | /8/1993 | F | SAMN60394230 | Illumina | SE | 0.19 | Hiseq 2500 | Hiseq 2500 |
| CGG_1_017165 | Mm7165 | MM98-1 | Svalbard | svalbard | 11/8/1998 | F | SAMN60394231 | Illumina, BEST | SE, PE | 9.24 | Novaseq | Hiseq 2500, Hiseq Xten, Novaseq |
| CGG_1_017166 | Mm7166 | MM98-2 | Svalbard | svalbard | 14/8/1998 | M | SAMN60394232 | Illumina, BEST | SE, PE | 10.14 | Novaseq | Hiseq 2500, Hiseq Xten, Novaseq |
| CGG_1_017167 | Mm7167 | MM98-3 | Svalbard | svalbard | 14/8/1998 | M | SAMN60394233 | Illumina, BEST | SE, PE | 7.07 | Novaseq | Hiseq 2500, Hiseq Xten, Novaseq |
| CGG_1_017173 | Mm7173 | 829 | Eclipse_Sound | Tremblay Sound | 15/8/1999 | F | SAMN60394234 | BEST | PE | 3.31 | Hiseq Xten | Hiseq Xten |
| CGG_1_017186 | Mm7186 | 964 | Hjornedal | Hjørnedal | /8/2013 | M | SAMN60394235 | BEST | PE | 8.45 | Novaseq | Hiseq Xten, Novaseq |
| CGG_1_017187 | Mm7187 | 965 | Hjornedal | Hjørnedal | /8/2013 | M | SAMN60394236 | Illumina, BEST | SE, PE | 8.92 | Novaseq | Hiseq 2500, Hiseq Xten, Novaseq |
| CGG_1_017188 | Mm7188 | 968 | Hjornedal | Hjørnedal | /8/2013 | M | SAMN60394237 | Illumina, BEST | SE, PE | 7.98 | Novaseq | Hiseq 2500, Hiseq Xten, Novaseq |
| CGG_1_017190 | Mm7190 | 987 | Hjornedal | Hjørnedal | 19/8/2012 | F | SAMN60394238 | BEST | PE | . | Novaseq | Hiseq Xten, Novaseq |
| CGG_1_017191 | Mm7191 | 988 | Hjornedal | Hjørnedal | 19/8/2012 | M | SAMN60394239 | Illumina, BEST | SE, PE | 10.38 | Novaseq | Hiseq 2500, Hiseq Xten, Novaseq |
| CGG_1_017192 | Mm7192 | 989 | Hjornedal | Hjørnedal | 19/8/2012 | M | SAMN60394240 | Illumina, BEST | SE, PE | 9.16 | Novaseq | Hiseq 2500, Hiseq Xten, Novaseq |
| CGG_1_017195 | Mm7195 | 994 | Hjornedal | Hjørnedal | 17/8/2012 | F | SAMN60394241 | Illumina, BEST | PE | 12.55 | Novaseq | Hiseq 2500, Novaseq |
| CGG_1_017198 | Mm7198 | 1000 | Smith_Sound | Iskant-Smith Sound | 14/6/2013 | M | SAMN60394242 | BEST | PE | 2.17 | Hiseq Xten | Hiseq Xten |
| CGG_1_017204 | Mm7204 | 996 | Hjornedal | Hjørnedal | 17/8/2012 | F | SAMN60394243 | BEST | PE | 10.09 | Novaseq | Novaseq |
| CGG_1_017208 | Mm7208 | AB87-105 | Admiralty_Inlet | Arctic Bay | 16/7/1987 | M | SAMN60394244 | BEST | PE | 11.29 | Novaseq | Hiseq Xten, Novaseq |
| CGG_1_017209 | Mm7209 | AB87-108 | Admiralty_Inlet | Arctic Bay | 17/7/1987 | M | SAMN60394245 | BEST | PE | 3.34 | Hiseq Xten | Hiseq Xten |
| CGG_1_017212 | Mm7212 | AB04-007 | Admiralty_Inlet | Arctic Bay | 2004 | M | SAMN60394246 | BEST | PE | 4.42 | Hiseq Xten | Hiseq Xten |
| CGG_1_017213 | Mm7213 | AB09-001 | Admiralty_Inlet | Arctic Bay | 15/8/2009 | F | SAMN60394247 | BEST | PE | 2.78 | Hiseq Xten | Hiseq Xten |
| CGG_1_017214 | Mm7214 | AB09-006 | Admiralty_Inlet | Arctic Bay | 18/8/2009 | F | SAMN60394248 | BEST | PE | 10.09 | Novaseq | Hiseq Xten, Novaseq |
| CGG_1_017217 | Mm7217 | BI95-288 | Eastern_Baffin_Island | Broughton Island | 2/8/1995 | M | SAMN60394249 | BEST | PE | 3.53 | Hiseq Xten | Hiseq Xten |

| | | | | | | | | | | | | |
|---|---|---|---|---|---|---|---|---|---|---|---|---|
| CGG_1_017224 | Mm7224 | BI07-1126 | Eastern_Baffin_Island | Broughton Island | 2007 | M | SAMN60394250 | Illumina, BEST | SE, PE | 2.34 | Hiseq 2500, Hiseq Xten | Hiseq 2500, Hiseq Xten |
| CGG_1_017231 | Mm7231 | CR04-1014 | Eastern_Baffin_Island | Clyde River | 2004 | F | SAMN60394251 | Illumina, BEST | SE, PE | 3.94 | Hiseq 2500, Hiseq Xten | Hiseq 2500, Hiseq Xten |
| CGG_1_017242 | Mm7242 | GF99-1010 | Jones_Sound | Grise Fiord | 16/8/1999 | M | SAMN60394252 | Illumina, BEST | SE, PE | 7.7 | Novaseq | Hiseq 2500, Hiseq Xten, Novaseq |
| CGG_1_017243 | Mm7243 | GF99-1015 | Jones_Sound | Grise Fiord | 16/8/1999 | M | SAMN60394253 | Illumina, BEST | SE, PE | 3.35 | Hiseq 2500, Hiseq Xten | Hiseq 2500, Hiseq Xten |
| CGG_1_017250 | Mm7250 | GF07-1131 | Jones_Sound | Grise Fiord | 28/7/2007 | F | SAMN60394254 | Illumina, BEST | SE, PE | 7.35 | Novaseq | Hiseq 2500, Hiseq Xten, Novaseq |
| CGG_1_017251 | Mm7251 | GF07-1137 | Jones_Sound | Grise Fiord | 27/7/2007 | M | SAMN60394255 | BEST | PE | 8.95 | Novaseq | Hiseq Xten, Novaseq |
| CGG_1_017252 | Mm7252 | IG95-441 | Somerset_Island | Igloolik | 3/8/1995 | F | SAMN60394256 | Illumina, BEST | SE, PE | 2.51 | Hiseq 2500, Hiseq Xten | Hiseq 2500, Hiseq Xten |
| CGG_1_017255 | Mm7255 | IG95-480 | Somerset_Island | Igloolik | 5/8/1995 | F | SAMN60394257 | Illumina, BEST | SE, PE | 2.43 | Hiseq 2500, Hiseq Xten | Hiseq 2500, Hiseq Xten |
| CGG_1_017261 | Mm7261 | PG96-524 | Eastern_Baffin_Island | Pangnirtung | 27/7/1996 | M | SAMN60394258 | Illumina, BEST | SE, PE | 3.94 | Hiseq 2500, Hiseq Xten | Hiseq 2500, Hiseq Xten |
| CGG_1_017281 | Mm7281 | RB99-1026 | Northern_Hudson_Bay | Repulse Bay | 6/7/1999 | M | SAMN60394259 | Illumina, BEST | SE, PE | 3.4 | Hiseq 2500, Hiseq Xten | Hiseq 2500, Hiseq Xten, Novaseq |
| CGG_1_017284 | Mm7284 | RB00-1100 | Northern_Hudson_Bay | Repulse Bay | 7/7/2000 | F | SAMN60394260 | Illumina, BEST | SE, PE | 5.23 | Hiseq 2500, Hiseq Xten | Hiseq 2500, Hiseq Xten, Novaseq |
| CGG_1_017285 | Mm7285 | RB01-1142 | Northern_Hudson_Bay | Repulse Bay | 27/7/2001 | F | SAMN60394261 | Illumina, BEST | SE, PE | 3.29 | Hiseq 2500, Hiseq Xten | Hiseq 2500, Hiseq Xten, Novaseq |
| CGG_1_017304 | Mm7304 | SB09-1053 | Somerset_Island | Taloyoak | 29/8/2009 | F | SAMN60394262 | Illumina, BEST | SE, PE | 2.4 | Hiseq 2500, Hiseq Xten | Hiseq 2500, Hiseq Xten |
| CGG_1_017305 | Mm7305 | SB09-1060 | Somerset_Island | Taloyoak | 29/8/2009 | F | SAMN60394263 | Illumina, BEST | SE, PE | 2.25 | Hiseq 2500, Hiseq Xten | Hiseq 2500, Hiseq Xten |
| CGG_1_017528 | Mm7528 | 363 | Eastern_Baffin_Island | Broughton I. | 1992 | M | SAMN60394264 | BEST | PE | 2.68 | Hiseq Xten | Hiseq Xten |
| CGG_1_017529 | Mm7529 | 574 | Eclipse_Sound | Tremblay Sound | 21/8/1997 | M | SAMN60394265 | Illumina, BEST | SE, PE | 1.96 | Hiseq 2500, Hiseq Xten | Hiseq 2500, Hiseq Xten |
| CGG_1_017531 | Mm7531 | 590 | Eclipse_Sound | Tremblay Sound | 21/8/1997 | M | SAMN60394266 | Illumina, BEST | SE, PE | 3.12 | Hiseq 2500, Hiseq Xten | Hiseq 2500, Hiseq Xten |
| CGG_1_017549 | Mm7549 | 869 | Melville_Bay | Melville Bugten | 26/8/2007 | F | SAMN60394267 | Illumina, BEST | SE, PE | 3.35 | Hiseq 2500, Hiseq Xten | Hiseq 2500, Hiseq Xten |
| CGG_1_017550 | Mm7550 | 870 | Melville_Bay | Melville Bugten | 3/9/2007 | F | SAMN60394268 | Illumina, BEST | SE, PE | 7.53 | Novaseq | Hiseq 2500, Hiseq Xten, Novaseq |
| CGG_1_017565 | Mm7565 | 1679 | Inglefield_Bredning | Qaanaaq | /8/1993 | M | SAMN60394269 | Illumina, BEST | SE, PE | 2.74 | Hiseq 2500, Hiseq Xten | Hiseq 2500, Hiseq Xten |
| CGG_1_017568 | Mm7568 | 1682 | Inglefield_Bredning | Qaanaaq | /8/1993 | M | SAMN60394270 | Illumina, BEST | SE, PE | 2.77 | Hiseq 2500, Hiseq Xten | Hiseq 2500, Hiseq Xten |
| CGG_1_017570 | Mm7570 | 1684 | Inglefield_Bredning | Qaanaaq | /8/1993 | F | SAMN60394271 | Illumina, BEST | SE, PE | 9.03 | Novaseq | Hiseq 2500, Hiseq Xten, Novaseq |
| CGG_1_017573 | Mm7573 | 1687 | Inglefield_Bredning | Qaanaaq | /8/1993 | M | SAMN60394272 | Illumina, BEST | SE, PE | 11.31 | Novaseq | Hiseq 2500, Hiseq Xten, Novaseq |
| CGG_1_017574 | Mm7574 | 1688 | Inglefield_Bredning | Qaanaaq | /8/1993 | F | SAMN60394273 | Illumina, BEST | SE, PE | 2.37 | Hiseq 2500, Hiseq Xten | Hiseq 2500, Hiseq Xten |
| CGG_1_017631 | Mm7631 | MM2012/01 | Svalbard | Kongsfjorden | 2012 | M | SAMN60394274 | BEST | PE | 9.75 | Novaseq | Hiseq Xten, Novaseq |
| CGG_1_024371 | NEG_1 | Mm19-01 | NE_Greeland | 79-24N, 17-21W | 6/9/2019 | M | SAMN60394275 | BEST | PE | 8.07 | Novaseq | Hiseq Xten, Novaseq |
| CGG_1_024372 | NEG_3* | Mm19-03 | NE_Greeland | 79-08,827N, 17-59,927W | 7/9/2019 | M | SAMN60394276 | BEST | PE | 8.59 | Novaseq | Hiseq Xten, Novaseq |

| | | | | | | | | | | | | |
|---|---|---|---|---|---|---|---|---|---|---|---|---|
| CGG_1_024373 | NEG_4 | Mm19-04 | NE_Greeland | 78-42,36N, 16-52,00W | 7/9/2019 | F | SAMN60394277 | BEST | PE | 8.22 | Novaseq | Hiseq Xten, Novaseq |
| CGG_1_024374 | NEG_5** | Mm19-05 | NE_Greeland | 77-29,837N, 17-37,486W | 8/9/2019 | F | SAMN60394278 | BEST | PE | 11.51 | Novaseq | Hiseq Xten, Novaseq |
| CGG_1_024375 | NEG_6 | Mm19-06 | NE_Greeland | 79-11,485N, 5-41.290W | 12/9/2019 | F | SAMN60394279 | BEST | PE | 11.99 | Novaseq | Hiseq Xten, Novaseq |
| NA | NA | PRJNA508363 | | Uummannaq | 1993 | | | Illumina | | 59.9 | Hiseq | Hiseq |
| NA | NA | PRJNA520934 | | Milne Inlet, Baffin island | 8/8/2013 | | | | | 56 | PacBio Sequel CLR; Dovetail Omni-C | |

* NEG3: Novaseq and Hiseq data were merged for PSMC, totalling a coverage of 15.0 x

** NEG5: Novaseq and Hiseq data were merged for PSMC, totalling a coverage of 20.3 x

**Supplementary Table S2. Demographic parameters estimated under the divergence model.** Parameter estimates as obtained in FASTSIMCOAL2 and the moments softwares. Estimates of effective size ($N_e$) are given in the number of haploids. Present and historical migration is given as a rate expressing the probability that any gene moves out from the population j to i at each generation. Time is given in generations. The estimated $log_{10}(L)$ of the model was equal to -11,995,868.492 under FASTSIMCOAL2.

| fastsimcoal | | moments |
|---|---|---|
| **Parameter** | **ML estimate** | **Difusion Approx.** |
| $N_e$ CAA/WG ($NPOP_0$) | 19,436 | 21,691 |
| $N_e$ SEG ($NPOP_1$) | 9,800 | 18,100 |
| $N_e$ NEG/Sv ($NPOP_2$) | 25,123 | 50,777 |
| $N_e$ Ancestral East ($NANC_{21}$) | 3,983 | 3,485 |
| $N_e$ Ancestral (NANC) | 9,425 | 5,253 |
| Split time CAA/WG - East Ancestral ($T_1$) | 10,188 | 2,223 |
| Split time SEG - NEG/Sv ($T_0$) | 1,234 | 562 |
| SEG-CAA/WG ($MIG_{01}$) | 4.98e-04 | 3.52e-07 |
| NEG/Sv - SEG ($MIG_{12}$) | 9.94e-04 | 9.97e-04 |
| CAA/WG - NEG/Sv ($MIG_{20}$) | 4.96e-04 | 1.9e-04 |
| CAA/WG - Ancestral East ($MIGA_{01}$) | 4.98e-04 | 2.53e-03 |

**Supplementary Table S3**. **Demographic parameters and search ranges as used in FASTSIMCOAL2.** Note that certain parameters were constrained by an upper limit preventing them from exceeding this threshold (bounded). In particular, the divergence time between the eastern and northeastern ancestor populations ($T_0$) could not exceed the estimated split time ($T_1$), which was capped at 50,000 generations. All other parameters were permitted to surpass the maximum value defined within the search range. All parameters, except for migration, were sampled from uniform distributions. The migration parameter was sampled from log-uniform distributions. Time is in units of generations and population sizes in the number of haploid individuals. See Supplementary Fig. S20 for a visual representation of the parameters.

| Parameter | Distribution | Minimum | Maximum | Bounded |
|---|---|---|---|---|
| $N_e$ CAA/WG ($NPOP_0$) | Uniform | 1.00E+03 | 4.00E+05 | no |

| | | | | |
|---|---|---|---|---|
| $N_e$ SEG ($NPOP_1$) | Uniform | 1.00E+03 | 4.00E+05 | no |
| $N_e$ NEG/Sv ($NPOP_2$) | Uniform | 1.00E+03 | 4.00E+05 | no |
| $N_e$ Ancestor SEG+North-East ($NANC_{21}$) | Uniform | 1.00E+03 | 4.00E+05 | no |
| $N_e$ Ancestor (NANC) | Uniform | 1.00E+03 | 4.00E+05 | no |
| Split time SEG+NEG/Sv ($T_0$) | Uniform | 100 | T1 | yes |
| Split time Ancestor ($T_1$) | Uniform | 1.00E+04 | 5.00E+04 | yes |
| 2NM | loguniform | 1.00E-04 | 10 | yes |

**Supplementary Table S4. Divergence time estimates** for fastsimcoal2, moments and SMC++ adjusting for the generation time of the narwhal.

| Parameter | fastsimcoal | moments | SMC++ |
|---|---|---|---|
| Split time CAA/WG - East Ancestral ($T_1$) | 305,640 | 66,690 | |
| Split time CAA/WG - NEG/Sv | | | 5,520 |
| Split time CAA/WG - SEG | | | 6,186 |
| Split time SEG - NEG/Sv ($T_0$) | 37,020 | 16,860 | 1,172 |

**Supplementary Table S5. Genes associated with PBS outliers.** List of the 35 genes associated with the population branch statistics (PBS) outliers in each population (99.99th percentile) and putative functions.

| **Population** | **Genes** | **Putative functions** |
|---|---|---|
| SEG | ATAD2B | chromatin binding and lysine-acetylated histone binding |
| SEG | ATP7B | Copper ion transmembrane transporter involved in the export of copper out of the cells |
| SEG | BTBD9 | protein-protein interactions |
| SEG | CCDC172 | |
| SEG | LOC114907696 | |

| | | |
|---|---|---|
| SEG | PNLIPRP3 | enable triglyceride lipase activity, involved in lipid catabolic process, active in extracellular space |
| SEG | SPOPL | enable ubiquitin protein ligase binding activity, involved in negative regulation of protein ubiquitination and proteasome-mediated ubiquitin-dependent protein catabolic process. |
| SEG | SUGCT | succinate-hydroxymethylglutarate CoA-transferase activity |
| CAA/WG | CAMTA1 | encodes a calcium-responsive transcriptional regulator that is highly expressed in the cerebral cortex and cerebellum, associated with human episodic memory that depends on the function of the hippocampus, influence long-term memory formation, and is associated with neuropsychological test performance |
| CAA/WG | ERBB4 | protein homodimerization activity and protein kinase activity, essential for neuronal development, may be important in activity-dependent synaptic plasticity, critical for learning and memory. ErbB4 signaling in dopaminergic axonal projections increases extracellular dopamine levels and regulates spatial/working memory behaviors. Also underlies the emergence of PV interneuron plasticity and social memory |
| CAA/WG | LOC114907278 | |
| CAA/WG | LOC114907296 | |
| CAA/WG | LOC114907300 | |
| CAA/WG | LOC114907338 | |
| CAA/WG | LOC114907518 | |
| CAA/WG | LOC114907519 | |
| CAA/WG | LOC114908175 | |
| CAA/WG | PGBD1 | nucleic acid binding and scavenger receptor activity |
| CAA/WG | SMARCC1 | ATP-dependent chromatin remodeler that regulates gene expression required for neural stem cell proliferation, differentiation, and survival during telencephalon development |
| CAA/WG | TRNAS-GCU_6 - also called SARS1 | encodes the cytosolic seryl-tRNA synthetase, which is responsible for charging tRNA with serin, catalyzes the first step of selenocysteine synthesis |

| | | |
|---|---|---|
| NEG/Sv | ACSL1 | convert free long-chain fatty acids into fatty acyl-CoA esters, plays a key role in both the synthesis of cellular lipids and the degradation of fatty acid |
| NEG/Sv | ALDH7A1 | Multifunctional enzyme mediating important protective effects, play a major role in the detoxification of aldehydes generated by alcohol metabolism and lipid peroxidation, involved in lysine catabolism that is known to occur in the mitochondrial matrix. |
| NEG/Sv | ASB7 | involved in protein degradation by acting as a bridge between substrate proteins and E3 ubiquitin protein ligases, regulates spindle dynamics and genome integrity, cytoskeletal organization during oocyte maturation |
| NEG/Sv | ITGA11 | involved in attaching muscle tissue to the extracellular matrix. |
| NEG/Sv | LOC114892843 | |
| NEG/Sv | LOC114893080 | |
| NEG/Sv | LOC114896581 | |
| NEG/Sv | LOC114897696 | |
| NEG/Sv | LOC114905412 | |
| NEG/Sv | MAP10 | promotes microtubule (MT) stability and participates in cytokinesis |
| NEG/Sv | NTPCR | RNA binding and ribonucleoside triphosphate phosphatase activity. |
| NEG/Sv | PHAX | RNA binding and toxic substance binding |
| NEG/Sv | PRORP | encodes the endonuclease subunit of the mitochondrial ribonuclease (RNase) P complex (EC 3.1.26.5), which functions in tRNA maturation |
| NEG/Sv | RABGAP1L | GTPase activator activity and small GTPase binding, promotes antiviral activity |
| NEG/Sv | TEX43 | act upstream of or within flagellated sperm motility |

### Supplementary Results

#### Population structure

##### 117-sample range-wide dataset

The best estimate for the number of populations (*K*) was determined using evalAdmix (Garcia-Erill and Albrechtsen 2020); *K*=3 was identified as the best fit (Supplementary Fig. S6A). Our findings indicated admixture in the lower coverage CAA/WG individuals, with either or both the SEG and NEG/Sv populations (Supplementary Fig. S5). This may be the result of the low coverage of our data, as we do not observe this admixture pattern in the 24-sample dataset with higher coverage (see below).

##### 24-sample, population-level dataset

For the 24-sample dataset, *K*=3 was also identified as the best fit (Fig. 1E, Supplementary Fig. S6B, S9). One of the 24 narwhals did not have a 100% membership proportion to its population of origin. This individual, CGG_1_017165, was sampled from Svalbard, and showed admixture proportions of 21% to CAA/WG, 52% to NEG/Sv, and 27% to SEG. However, in runs with lower likelihoods, other single individuals within NEG/Sv, showed admixture, at similar levels (Supplementary Fig. S9B), highlighting that the admixture results within NEG/Sv are not robust.

We did not find any substructure within the eight individuals from CAA/WG from the 24-sample dataset, which included two individuals from Admiralty Inlet, three from Jones Sound, one from Melville Bay and two from Inglefield Bredning. However, the sample size may be insufficient for meaningful inferences on population subdivision.

#### Evolutionary relationships

##### Treemix

We inferred the best number of migration events to 1 using an ad hoc analysis of the second order rate of change in the log-likelihood (Evanno) method implemented in the R package OptM using the finless porpoise as the outgroup. 99.99% of the variance was explained at M=1. For the beluga, Treemix could not infer more than 1 migration edge from the data, even when running >1 migration event and therefore the OptM package could not be run. The increase in likelihood between 0 and 1 migration event was similar as when using the finless porpoise as the outgroup. All the internal branch lengths were very short, indicating that each population experienced low levels of population-specific drift (Fig. 2A, Supplementary Fig. S11). We observe a first split between east of Greenland (Northeast Greenland/Svalbard (NEG/Sv) and Southeast Greenland (SEG)) and west of Greenland (Canadian Arctic Archipelago/West Greenland (CAA/WG)), and then a split between the two populations east of Greenland. A migration event is inferred from the CAA/WG lineage to NEG/Sv. When using the finless porpoise the migration edge is inferred from the CAA/WG population while when using the beluga, it is inferred from the ancestral population to the CAA/WG population.

##### *D*-statistics

The topology (SE, NEG; BB, OUT) did not significantly deviate from 0 (Z score of 0.18). The topologies (SE, BB; NEG, OUT) and (NEG, BB; SE, OUT) were significantly negative indicating SE and NEG share an excess of derived alleles.

#### Genetic diversity

##### ROH

We find SEG individuals present the highest mean maximum ROH length and highest mean ROH length, but differences are not statistically significant among the three populations (Kruskal–Wallis $\chi^2 = 4.81$, df = 2, P = 0.09 for the maximum and (n=24, ANOVA, F(2,21)=2.32, P=0.12, and pairwise comparisons are not significant after correcting for multiple comparisons, Supplementary Fig. S28). The proportion of the genome in ROH ($F_{ROH}$) does not differ significantly among the three populations (ANOVA, F(2,21)=3.10, P=0.07), although SEG narwhals have a significantly higher $F_{ROH}$ than the CAA/WG narwhals, after correcting for multiple comparisons (Tukey HSD $P_{CAA/WG-SEG} = 0.05$, Fig. 4B).

We found no differences among populations in total size of small ROH (0.5 to 1.5 Mb, reflecting ~2640 – 870 years ago (Kruskal-Wallis $X^2 = 0.37$, df = 2, p-value = 0.83, none of the pairwise comparisons are significant), indicating past inbreeding due to low long-term effective population size and similar demographic histories in the more distant past (Fig. 4C). The two eastern populations had a larger part of their genome in ROH of intermediate size: 1.5 Mb to 4 Mb, reflecting ~870 – 330 years ago, than CAA/WG (ANOVA F(2,21) = 10.24, P = 0.0008, $P_{CAA/WG-NEG/Sv} = 0.03$ and $P_{CAA/WG-SEG} = 0.0002$ - although the significance was marginal for CAA/WG vs NEG/Sv when correcting for multiple comparisons Tukey HSD $P_{CAA/WG-NEG/Sv} = 0.07$ and Tukey HSD $P_{CAA/WG-SEG} <0.001$). The mean total length of intermediate ROH was also marginally significantly higher in SEG than in NEG/Sv ($P_{NEG/Sv-SEG} = 0.03$, but only 0.06 after correcting for multiple comparisons). Although the mean total length of long ROH (> 4 Mb) is only marginally different among populations (Kruskal-Wallis $X^2 = 5.42$, df = 2, p-value = 0.07), SEG narwhals have the highest mean total length of long ROH (> 4 Mb, reflecting the past ~330 years). This metric is significantly higher in SEG narwhals than in CAA/WG narwhals (Tukey HSD $P_{CAA/WG-SEG} =0.05$), but not different from NEG/Sv narwhals (Tukey HSD $P_{CAA/WG-NEG/Sv} = 0.08$).

Mutation load

In the whole dataset, we find 507 high impact deleterious mutations and 21,856 moderately deleterious mutations. We did not find significant differences among the three populations for high impact mutations in the total genetic load (F = 0.75, DF = 2,21, P = 0.49), masked load (F = 0.27, DF = 2,21, P = 0.76) and realised load (F = 1.50, DF = 2,21, P = 0.24) and for moderate impact mutations in the total genetic load (F = 0.06, DF = 2,21, P = 0.94), masked load (F = 0.05, DF = 2,21, P = 0.95) and realised load (F = 0.17, DF = 2,21, P = 0.84).

### Supplementary Methods

#### Samples and localities

We collected tissue samples from 121 narwhals sampled from across the species range from all 12 narwhal stocks recognized by the North Atlantic Marine Mammal Commission (NAMMCO) (Fig. 1A). A stock refers to a summer aggregation and constitutes a management unit based on distribution, telemetry, ecological and, in some cases, genetic data (Hobbs et al. 2019). Three stocks included several sample localities (Fig. 1A). Samples were collected during subsistence hunts and through biopsy sampling during satellite-tagging studies or, for the samples from Northeast Greenland during targeted biopsy sampling from an helicopter. All samples were collected during summer months between 1982 and 2019 (Supplementary Table S1). Samples were sent to Denmark under CITES exemption number DK003. From those 120 samples, we ran preliminary analyses of population structure and then sequenced further 24 samples from the three genetic clusters we identified: Canadian Arctic Archipelago and West Greenland (CAA/WG); Northeast Greenland and Svalbard (NEG/Sv); and Southeast Greenland (SEG, Fig. 1C) to sufficient coverage for demographic history, inbreeding and selection analyses. We added a 25th sample, not included in the 120 samples, coming from SEG. Note that these sample numbers include related individuals, and therefore the final number of samples included in the analyses differs (117 and 24 sample-dataset respectively). We included publicly available high coverage genome data in the long-term demographic history analysis from previous studies from two individuals collected in the Canadian Arctic Archipelago and West Greenland (coverage of 59.9× and 56.0×, one sampled in Uummannaq/West Greenland in 1993 (Westbury et al. 2019, PRJNA508363) and the other sample in Baffin island in eastern Canada in 2013 (PRJNA520934).

We used the *Monodontidae* sister species beluga, *Delphinapterus leucas*, and a member of the closest family to the *Monodontidae (*McGowen et al. 2020), the finless porpoise, *Neophocaena phocaenoides*, which is part of the *Phocoenidae* to reconstruct the ancestral states of alleles and as outgroups. Hybridization can occur between narwhals and belugas (Skovrind et al. 2019), which is an additional reason why we ran our analyses twice, with the beluga or the finless porpoise as outgroups, and we checked for consistency of the results. The finless porpoise SRR940959 originated from South Korea (Yim et al. 2014) and the beluga SAMN12242088 originated from Bristol Bay in Alaska, where narwhals are not known to currently occur.

#### DNA laboratory work

We extracted DNA from tissue samples using the Qiagen Blood and Tissue Kit following the manufacturer's protocol with minor modifications. The volume of proteinase K was increased to 40 µL and the incubation time was extended to 24 hours. Genomic DNA was diluted to 15 ng/µL. Libraries were prepared and sequenced according to two different protocols.

For 102 samples, the DNA was sheared to an average size of ~350-550 bp using a Bioruptor run with 4 cycles of 25 seconds on, and 90 seconds off. Further cycles were added if the fragments were too long. Libraries were built from the fragmented DNA extracts using Illumina NeoPrep following the NeoPrep Library Prep System Guide applying default settings. PCR amplification, quantification and normalization were all carried out by the NeoPrep Library Prep System. The libraries were screened for size distribution on an Agilent 2100 Bioanalyzer and pooled in equimolar ratios before sequencing on an Illumina HiSeq 2500 with 80bp SE technology.

For 50 samples (including 31 out of the 102 samples and an additional 19 samples), DNA was fragmented using the Covaris M220 Focused-ultrasonicator to create ~350-550 base pair (bp) fragment lengths. Settings included a minimum, set and maximum temperatures of 18, 20 and 22 °C, respectively. We used a treatment of 50 seconds, peak point of 75, duty factor of 10 and 200 cycles/burst. Paired-end (PE) sequencing libraries were built on the sheared DNA extracts using the BEST protocol (i.e. Blunt-End Single-Tube library building for modern and ancient DNA (Carøe et al. 2018)). This method involved three steps: i) end repair with an incubation of 30 min at 20°C followed by 30 minutes at 65°C and cooling at 4°C; ii) ligation with an incubation of 30 min at 20°C followed by 30 minutes at 65°C and cooling at 4°C; iii) fill-in buffer with an incubation of 20 min at 65°C followed by 20 minutes at 80°C and cooling at 4°C. Libraries were cleaned using SPRI bead purification following (Rohland and Reich 2012).

Libraries were subsequently double-index amplified with unique 6 bp sequences for 15 cycles using TaqGold Polymerase (5U/μl), 10X Buffer, 25 mM MgCl2, 25 Mm dNTPs, in 50 μL or 100 μL reactions. The libraries were purified using SPRI beads. The DNA concentration of the libraries was measured using a TapeStation. Libraries were pooled approximately equimolarly and sequenced on an Illumina HiSeq Xten with 150bp PE technology for 48 samples and on a Novaseq 6000 with 150bp PE technology for 28 samples. Of these 26 were sequenced on both platforms, including samples from NEG. To avoid any differences in coverage and issue with differences in sequencing platforms, we did not merge data from Hiseq and Novaseq and only used the Novaseq data for the 25 samples of the population-level dataset but are making all data accessible on SRA (Table S1). For NEG samples, we included both Novaseq and Hiseq for PSMC to increase coverage.

#### Bioinformatics

##### Mapping

Sequencing reads were processed using Paleomix v. 1.2.13.1 (Schubert et al. 2014). The first step involved trimming residual adapter sequence contamination from FASTQ reads as well as low-quality stretches at read ends (i.e. consecutive stretches of N's and of bases with a quality score of ≤3) using AdapterRemoval v. 2.2.2 (Schubert et al. 2016). Sequence reads that were ≤25 bp following trimming were discarded. Read quality was inspected using FastQC. The remaining reads were mapped to the narwhal reference genome (GCF_005190385.1) using bwa v. 0.7.15 with the mem algorithm (Li and Durbin 2009) requiring a mapping quality score of 30 for the narwhal raw sequencing reads, while we use a mapping quality score of 20 for the finless porpoise and beluga raw sequencing reads given we are mapping to another species (the narwhal). Read groups were added using picard-tools v. 2.6.0 (http://broadinstitute.github.io/picard/faq.html). The same program was used to merge the bam files from each individual from different lanes and remove duplicate reads. Indel realignment was performed using GATK v. 3.8.1 (McKenna et al. 2010). We set a filter of 20 for the base score quality as we include Novaseq data. We only considered bases with a base score quality of 20, as our data includes Novaseq sequencing, with the two-color chemistry.

We only kept the 22 largest scaffolds (> 35000 Mb, NW_021703765.1 to NW_021703786.1) and further filtered the data to only keep the autosomes. To identify the scaffolds corresponding to the X and Y chromosomes, we mapped the narwhal reference genome to the X chromosome of the cow (GCA_002263795.2) and the Y chromosome of the human (GCA_000001405.28) using the SeXY pipeline (Cabrera et al. 2022). We removed part of NW_021703768.1, NW_021703774.1 and NW_021703780.1 and all of NW_021703779.1 using bedtools v. 2.25.0 and samtools v. 1.2 (Li et al. 2009; Quinlan and Hall 2010). In addition, we also ran a PCA for each large scaffold to see if we detect any show clustering by sex, as we had issues with individuals clustering by sex when mapping to the beluga reference genome (ASM228892v2_HiC) (Dudchenko et al. 2017, 2018; Jones et al. 2017) in preliminary analyses when the narwhal reference genome used in this study was not yet available. We identified two scaffolds where individuals clustered by sex rather than geographical locations: NW_021703765.1 and NW_021703771.1. These two scaffolds were removed using a “region” filter in analyses run using ANGSD v. 0.939 (Korneliussen et al. 2014), or from the vcf file in analyses based on genotype calling.

Repeat regions were extracted from the GENBANK reference genome website and were removed from the bam files using bedtools and samtools. We also removed regions of excessive coverage, as the high coverage of these regions can potentially be the result of unmasked repeat regions in nuclear mitochondrial DNA (NUMTs), or some other mapping artifact (e.g. paralogous loci). Coverage was estimated for the 120-sample dataset and the 25 sample-dataset using the doDepth function in ANGSD. Regions that were more than twice the mean coverage (>639x for the 120-sample dataset and >481x for the 25-sample dataset) were considered of excessive coverage. These regions were detected using the CALLABLELOCI tool in GATK. Then they were removed from the bam files using bedtools and samtools as above. Mean coverage was again estimated using the doDepth function in ANGSD and plotted using R v. 4.0.5 (R Core Team 2021).

##### Variant calling

We called SNPs taking genotype uncertainty into account by calculating genotype likelihoods in ANGSD for both the 120- and 25-sample dataset, creating a beagle file. We used the following options for the 25 sample dataset: -GL 1 -minMapQ 30 -minQ 23 -doGlf 2 -doMajorMinor 1 -doMaf 2 -SNP_pval 1e-6 -minMaf 0.05 -minInd 18 -skipTriallelic 1 -remove_bads 1 -uniqueOnly 1 -doCounts 1 -setMinDepthInd 3 -rf regions2.txt. The region filter was used to remove the two scaffolds showing clustering by sex in the PCA.

We called genotypes (i.e. generation of a vcf file) for the 25-sample dataset using bcftools v. 1.12 mpileup multiallelic and rare-variant calling, option –m on the filtered bam files (Li et al. 2009; Danecek et al. 2014). Variable sites with a minimum phred score quality of 20, minimum depth of 3, and genotype quality of 20 were retained in vcftools v. 0.1.16 (Danecek et al. 2011). We kept SNPs with a minimum MAF of 0.05 and having genotype data in a minimum of 75% of all the individuals and a minimum of five individuals in each of the three populations. The vcf file was also filtered for monomorphic and non-biallelic sites. Coverage was estimated using vcftools.

#### Population structure analyses

##### Inferring relatedness

We ran NgsRelate (Korneliussen and Moltke 2015; Hanghøj et al. 2019) to identify pairs of related individuals both in the 120-sample and 25-sample datasets using the pairwise relatedness measure (Hedrick and Lacy 2015). Note that to obtain reliable relatedness estimates, NgsRelate should generally be run within populations, as relatedness estimates should be based on population specific allele frequencies. Our main aim here was only to remove strongly related individuals as those were driving our population structure results, and we therefore ran NgsRelate on the full dataset (120-sample dataset) and did not run NgsRelate per identified population. We used the following options: -gl 1 -domajorminor 1 -snp_pval 1e-6 -domaf 2 -minmaf 0.05 -doGlf 3 -minMapQ 30 -minQ 23 -skipTriallelic 1 -remove_bads 1 -uniqueOnly 1 -doCounts 1 -rf regions2.txt (to exclude the two scaffolds showing clustering by sex). We ran NgsRelate with and without a minimum depth filter, and results were similar. We removed three individuals in pairs of related narwhals from the CAA/WG population in the 120-sample dataset, therefore referred to as 117-sample dataset (r values were 0.50, 0.48 and 0.23) and one individual from the SEG population in the 25-sample dataset, therefore referred to as 24-sample dataset (r value was 0.19). We plotted the pairwise relatedness coefficients in heatmaps for the 120- and 117 sample-dataset using R v. 4.0.5 (R Core Team 2021) and packages ggplot2 (Wickham 2016a), dplyr (Wickham et al. 2023), tidyr (Wickham and Girlich 2022) and readr (Wickham et al. 2022).

##### Inferring linkage disequilibrium

We used NgsLD (Fox et al. 2019) to obtain a set of unlinked SNPs both for the 117-sample and 24-sample dataset. We first ran it with a maximum distance of 1,000 kb for SNPs to be considered in linkage disequilibrium (LD) and LD decay was inspected using R. As LD decay was decreasing rapidly, NgsLD was then re-run using a maximum distance of 100 kb. LD decay was inspected and showed LD stabilized after a distance of 20 kb. Then, a set of unlinked sites was produced considering SNPs are in LD until 20 kb, and using a minimum weight of 0.5. Population structure analyses were run on the set of 486,709 unlinked SNPs for the 117-sample dataset and 274,852 SNPs for the 24-sample dataset. Note that population structure results were similar when including (299,268 SNPs) or when excluding the two scaffolds showing clustering by sex (274,852 SNPs). Thus, for PCA and NGSadmix, we only show the results obtained using 274,852 SNPs. All the other analyses (demographic history or selection) were run on the unpruned set of SNPs (excluding the two scaffolds showing clustering by sex) as either the program include the pruning itself (e.g. Treemix) or linkage information is important for the inferences.

#### PCA and admixture analyses

To infer population structure, we used PCAs in PCAngsd (Meisner and Albrechtsen 2018) and admixture analysis in NGSAdmix (Skotte et al. 2013) using the sets of 486,709 for the 117-sample dataset and 274,852 unlinked SNPs for the 24-sample dataset. NGSAdmix was run 100 times for each *K* value between 2 and 5, using a tolerance for convergence of $1e^{-10}$ and a minimum likelihood ratio value of $1e^{-6}$. Consistency between runs was checked, and the runs with the highest likelihood for each *K* value were plotted in R.

We used evalAdmix (Garcia-Erill and Albrechtsen 2020) to estimate the best number of clusters (*K*) using the NGSadmix results from both datasets. EvalAdmix outputs a matrix of pairwise correlation of residuals between individuals. Correlation values close to 0 indicate a good fit of the data to the admixture model. When the admixture model does not fit the data, individuals with similar demographic histories (usually from the same population) will be positively correlated, while individuals with different histories, but modelled as having shared ancestry, will be negatively correlated. We plotted the residual matrix using R v. 4.0.5. In both datasets, we observe that for *K*=2, individuals from SEG and individuals from NEG/Sv show negative correlation, while individuals within each region are positively correlated (Supplementary Fig. S6). This indicates that they are two separate populations. For K=3, we observe a much better fit of the data to the admixture model (Supplementary Fig. S6). We also checked admixture proportions amongst the 100 runs for various R values, and found they were consistent among runs for *K*=3, but not for higher values of *K*.

We run PCAngsd on the 117- and 24-sample datasets, as well as on a dataset of 101 Canadian Arctic Archipelago and West Greenland narwhals (101-sample dataset) to test for fine-scale population structure within this region. We plotted the results from PCAngsd using R.

For the 117- and 101-sample datasets, we also ran the single-read PCA method in ANGSD, which involves random sampling of a single read for each sample at each site. Note that we did not perform LD pruning for this analysis, as it is run directly on the bam files. We used the following options for the 117 samples: -minMapQ 30 -minQ 23 -GL 1 -doMajorMinor 1 -doMaf 2 -SNP_pval 2e-6 -doIBS 1 -doCounts 1 -minInd 58 -doCov 1 -makeMatrix 1 -minFreq 0.05 -setMinDepth 106 -remove_bads 1 -uniqueOnly 1 -rf regions2.txt (to remove the scaffolds were individuals clustered by sex). We set the minimal depth as ⅓ of the mean depth as estimated in ANGSD using the doDepth function. We also ran the analyses with a minFreq (i.e. MAF) of 0.01. For the 101-sample dataset, we used the same options, but changed -minInd to 50 and -setMinDepth to 56. We plotted the results using R.

#### Fixation statistics

For the 117-sample dataset, we estimated $F_{ST}$ using ANGSD. We computed the unfolded site frequency spectrum (SFS) for each population, and the 2D-SFS for each pair of populations in ANGSD and winsfs using a two step procedure (Nielsen et al. 2012) using the finless porpoise as the ancestral state, as well as the folded 2D-SFS. First, We used the dosaf 1 function to calculate the site allele frequency spectrum likelihood (saf) based on individual genotype likelihoods assuming Hardy-Weinberg equilibrium (HWE). We then used winsfs (Rasmussen et al. 2022) to optimize the saf and estimate the SFS and the 2D-SFS for each pair of populations as it better accounts for low-coverage data. We estimated $F_{ST}$

between pairs of populations using the realSFS fst index function. We used the same procedure for the 24-sample dataset, apart from that we used ANGSD to estimate the 2D-SFS.

#### Evolutionary relationships

##### Treemix

We reconstructed the evolutionary relationships of the three inferred narwhal populations as a Maximum Likelihood bifurcating tree using TreeMix v. 1.13 (Pickrell and Pritchard 2012). We used stacks v 2.61 to generate the Treemix input file using the populations function. We ran TreeMix using either one individual beluga or one individual finless porpoise as a root. We first ran TreeMix ten times for each value of M (migration events) ranging from 0 to 4, using blocks of 1,000 SNPs to estimate the covariance matrix (-k 1000), and the global flag (-global). We estimated the optimal number of migration events as 1 using the optM R package (https://cran.r-project.org/web/packages/OptM/index.html). We then ran TreeMix 100 times for 1 migration event and obtained a consensus tree and bootstrap values using the R package BITE (Milanesi et al. 2017). We also estimated the residual covariance matrix for the consensus tree using TreeMix.

##### Admixture graphs - qpBrute and Admixture Bayes

We sought to reconstruct the evolutionary history of the three populations as an admixture graph (Patterson et al. 2012). We used a heuristic search algorithm, qpBrute (https://github.com/ekirving/qpbrute) which explores the space of all possible admixture graphs of a given maximum complexity, under a brute-force approach (Ní Leathlobhair et al. 2018; Liu et al. 2019). We ran the analysis using either the finless porpoise (823,928 SNPs) and the beluga (768,978 SNPs) as outgroups and results were similar.

We ran qpBrute using the following parameters: outpop: NULL, useallsnps: YES, blgsize: 0.05 (5 Mb which is the block size for Jackniffe), forcezmode: YES, lsqmode: YES, diag: .0001, bigiter: 6, hires: YES, lambdascale: 1, inbreed: NO. As we have only one individual to root the graph, we did not attempt to use the allele frequency of the outgroup to normalise the weighting of each SNP in the ingroup and we used the option output: NULL that is SNPs are flat-weighted. In qpBrute, leaf nodes were added to the graph using a stepwise addition order algorithm. At each step, the insertion of a new node was tested at all branches of the graph, apart from the outgroup branch. All possible admixture combinations were tried where a node could not be added without producing *f4*-statistics outliers (i.e. $|Z| \geq 3$). The sub.graph was discarded when a node could not be inserted via these approaches. Where a node was successfully added, the remaining nodes were recursively inserted into the graph. The package tried all possible six starting graph orders, and we found three graphs with no *f4* outliers among a total of 14 unique graphs, two of which were mirror graphs of each other.

We computed the mean log-likelihoods of the three models and their Bayes Factors using the MCMC algorithm implemented in the R package ADMIXTUREGRAPH v. 1.0.289 using the default chain settings implemented in qpBrute (two chains, each with two million iterations, five heated chains, a burn-in of 50%, and no thinning). We assessed the convergence of the chains using the output from the R package CODA v. 0.19-490

(Plummer et al. 2006) also generated in qpBrute. The Bayes factors indicated that the models had similar log-likelihoods and were equally likely (K<0.1). We increased the chains to four million iterations with a burn-in of three millions, but it did not help in discriminating between the three graphs.

We also used Admixture Bayes to estimate high-probability admixture graphs (Nielsen et al. 2023). Admixture Bayes is a Bayesian approach that uses a reversible jump Markov Chain Monte Carlo (MCMC) to sample high-probability admixture graphs. It also does not require any a priori information on the topology of the graph or number of migration events. It has the advantage of searching the entire state space to find the best fitting graph(s) and to provide a level of confidence in the sampled graphs. We ran the analysis twice, using either the finless porpoise or the beluga as the outgroup. We ran three independent MCMC chains with 400,000 iterations and 16 chains to run the Metropolis-Coupled MCMC (MCMCMC) of the "runMCM" step. We used the default values for the "analyzeSample" step discarding the first 50% samples as burn-in and a thinning rate of 10 and the –slower option to compute branch length and admixture proportion estimates. We also assessed convergence by examining the trace plots of the chains and Gelman-Rubin convergence diagnostic of parallel chains using the EstimateConvergence R script (https://github.com/avaughn271/AdmixtureBayes). We plotted the graphs using the makeplot.py script and including the –write_ranking options to get the posterior probabilities of the best graphs.

##### *D*-stats /*f3*-statistics

We used *D*-statistics and *f3*-statistics to further assess the relationships of three narwhal populations. *D*-statistics describe an excess of shared derived alleles between taxa which could be the result of introgression or ancestral population structure. It thus allows to detect departure from the 'tree-ness' of a given topology (Green et al. 2010; Durand et al. 2011; Patterson et al. 2012). We calculated *D*-statistics of the form (H1, H2; H3, OUT): (SEG, CAA/WG; NEG/Sv, OUT), (SEG, NEG/Sv; BB, OUT) and (NEG, BB; SE, OUT) using the qpDstat tool in Admixtools v. 7.0.2 (Patterson et al. 2012) with the f4mode option enabled using either the finless porpoise or the beluga as the outgroup (OUT).

It is important to note that Admixtools calculates the *D*-statistics using the following formula: *D* = (nBABA-nABBA) / (nBABA+nABBA).
nBABA represents the number of sites where only H1 and H3 share a derived allele (BABA sites) and nABBA the number of sites where only H2 and H3 share a derived allele (ABBA sites). Under the null hypothesis assuming that the given topology is correct, we expect equal numbers of BABA and ABBA sites and thus *D* = 0. A statistic differing significantly from 0 indicates either gene flow between one in-group population (H1 or H2) and H3, or that the tree is incorrect. To test for significance, we calculated a Z-score based on blocked jackknife estimates of the standard deviation of the *D*-statistics; block size was 5 Mb which should be higher than the LD in the populations. This Z-score relies on the assumption that the *D*-statistics, under the null hypothesis, follows a normal distribution with mean 0 and a standard deviation equal to a standard deviation estimate computed using the "delete-m jackknife for unequal m" method described in Busing et al 1999 (Busing et al. 1999).

We also estimated *f3*-statistics, which measure allele frequency correlations between populations (Patterson et al. 2012), to test whether a target population (C) is admixed between two source populations (A and B). They are calculated as the product of allele frequency differences between population C to A and B, respectively:
$F3(C;A,B)=\langle(c-a)(c-b)\rangle$

We estimated *f3*-statistics using the threepop function in Treemix. They were estimated in 797 blocks of size 1000, totalling 797,565 SNPs. We estimated the statistics for the data with the individuals correctly assigned to their populations and after randomly reshuffling the individuals among populations to test whether our results differ from random.

#### Demographic reconstruction

##### Population size reconstruction with PSMC, SMC++ and stairwayplot

We used the pairwise sequential Markovian coalescent (PSCM (Li & Durbin, 2011) to infer deep-time changes in effective population size ($N_e$) through time. To infer more recent demographic history we used the methods Sequential Markov Coalescent + plenty of unlabelled samples (SMC++) (Terhorst et al. 2017) and stairwayplot (Liu and Fu 2015, 2020), which are based on population scale data and can include individuals with lower coverage than PSMC. We also used SMC++ to estimate divergence times among the three inferred populations. We scaled the results of all three analyses using a generation time of 30 years (Garde et al. 2015), and the per-generation mutation rate previously estimated for the species: 1.56e-08 (Westbury et al. 2019). We plotted all the results in R with packages ggplot2, scales (Wickham 2016b), RcolorBrewer (Neuwirth 2014) and reshaped2 (Wickham 2007)**.**

##### PSMC

We ran PSMC on single individuals with the highest coverage from each of the three identified populations: two published genomes from Canadian Arctic Archipelago and West Greenland (coverage of 59.9 and 56.0×) (Westbury et al. 2019), three genomes new to this study including two from Northeast Greenland and Svalbard (20.3× and 15.0×), and one genome from Southeast Greenland (12.7×).

We built a consensus sequence for each bam file in fastq format using i) the samtools mpileup command with the -C50 option to reduce the effect of reads with excessive mismatches, ii) bcftools view -c to call variant in a vcf file and iii) the perl utility vcfutils.pl vcf2fq to transform the generated vcf file to fastq format. We removed sites with less than a third or more than twice the mean depth of coverage, as estimated in ANGSD, and with a phred score less than 20. We only used autosomes, and removed the scaffolds where individuals clustered by sex.

We ran the PSMC inference using the parameter values for human autosomes, which are as follows: 25 iterations, maximum TMRCA ($T_{max}$) = 15, number of atomic time intervals (n) = 64 (following the pattern (1*4 + 25*2 + 1*4 + 1*6), and initial theta ratio (r) = 5. We also ran PSMC with 100 rounds of bootstrapping.

We acknowledge that two of our samples have depth of coverage <20× which may lead to false negative detection of heterozygous sites and impact the results (Nadachowska-Brzyska et al. 2016). However, we observed highly similar trajectories across samples until ~40,000 years ago, and PSMC results are not reliable in the past ~20,000 years (Li & Durbin, 2011). We therefore did not apply any correction related to coverage to our analyses. To infer more recent demographic history we used the methods SMC++ and stairwayplot, which are based on population scale data and can include individuals with lower coverage than PSMC.

##### SMC++

SMC++ incorporates both the SFS and LD information in a coalescent Hidden Markov Model (HMM) (Terhorst et al. 2017). SMC++ is an extension of the PSMC method, for a larger number of unphased samples per population. PSMC uses the distribution of heterozygous sites throughout the genome where the heterozygosity information is emitted as binary, while SMC++ emits the allele frequency of an extra n-2 haplotypes. The latter is based on the SFS conditioned on the time to the most recent common ancestor (TMRCA) of a single 'distinguished individual'.

We generated the vcf file as described earlier, but no MAF filter was applied, as the analysis is based on the SFS. The vcf file was converted to SMC++ format using the vcf2smc function for each retained autosomal scaffold. The regions identified as repeats and showing excessive coverage were included in a mask file, so that they were not misidentified as very long runs of homozygosity, which could impact the population trajectories and create false sudden decreases in $N_e$ in the last 10-100 generations.

As mentioned earlier, SMC++ estimates the SFS conditioned on the TMRCA of a single individual, called "distinguished" individual hereafter. We made the distinguished individual vary over three individuals with the highest coverage within each of the three populations, which has the advantage of incorporating genealogical information from additional individuals into the analysis, and may lead to improved estimates.

Population size histories were estimated using the estimate option in SMC++ using the default settings for the estimate function, and the narwhal's generation time and mutation rate (Garde et al. 2015; Westbury et al. 2019). As recommended on the program's github we tried different thinning parameters (1,500 and 2,000) in addition to the default and obtained similar results. To estimate population splits, we first estimated population histories using the estimate option, i.e. the marginal estimates. We computed the joint frequency spectrum for each pair of populations using the vcf2smc function. We used the split function to refine the marginal estimates into an estimate of the joint demography and estimate divergence times between pairs of populations.

##### Stairwayplot

We also used stairwayplot v. 2 (Liu and Fu 2015, 2020) to reconstruct changes in $N_e$ through time. Stairwayplot is based on the SFS and calculates the expected composite likelihood of a given SFS, treating each SNP independently to reduce the computational costs. We ran the analysis using the unfolded SFS, generated using both winSFS and ANGSD, and obtained similar demographic trajectories; the results based on the SFS generated in winsfs are presented here. We ran the methods both including and excluding singletons, and obtained similar trajectories, results including singletons are therefore presented.

##### GONE

We ran GONE (Santiago et al. 2020) using the default parameters and also changed the hc parameters from 0.05 to 0.01 for one of the population (CAA/WG). To format the file for GONE, we sub-sampled the vcf file to only include the individuals from each population and used Plink v. 1.90b6.21 (Purcell et al. 2007) to generate the .map format needed by the program.

##### Demographic Modeling with FASTSIMCOAL2 and moments

To investigate the historical processes that shaped the current population structure and take gene flow into account, we estimated demographic parameters from the site frequency spectrum (SFS) using FASTSIMCOAL2 v. 2.7 (Excoffier et al. 2013, 2021) and moments softwares (Jouganous et al. 2017).

We built the demographic models based on the evolutionary relationship results from Treemix and admixture graphs (Fig. 2A, Supplementary Fig. S11-S14).The joint folded 3D-SFS was estimated from winsfs v. 0.7.0 (Rasmussen et al. 2022) as most samples had depth ~10x and the unfolded SFS showed an excess of derived alleles. We used winsfs that iteratively estimates the SFS in small blocks of data and updates the overall estimate by averaging the block estimates over a genome window and has the advantage of reducing overfitting, especially for multidimensional SFS. We used the following options to generate the saf file for each of the three populations: -doSaf 1 -anc ancestral.fa -GL 1 -minMapQ 30 -minQ 23 -remove_bads 1 -uniqueOnly 1 -minInd 7 -rf regions.txt before generating the 3D-SFS in winsfs.

FASTSIMCOAL2 implements a composite-likelihood method using coalescent simulations to infer divergence times, population tree topologies and historical migration rates from the site frequency spectrum. On the basis of the population structure analysis, we implemented a simple demographic model with three populations, corresponding to CAA/WG; NEG/Sv and SEG. The model assumes that an ancestral population first split into two populations, the CAA/WG and the ancestral eastern populations at time T1 generations ago. Later, the SEG and NEG/Sv populations further diverged at time T0 generations ago. We assumed constant and symmetric migration events among all present and ancestral populations (Supplementary Fig. S20). A summary of the defined parameters and their search ranges are given in Supplementary Table S3**.** Input files including template and parameter estimation files (.tpl and .est, respectively) are available in the github repository (see data availability section).

We considered the minor allele frequency spectrum (folded SFS) due to an observed excess of fixed derived alleles, indicating potential misidentification of the ancestral state. The mutation rate was set to 1.56e-08 (Westbury et al. 2019). We estimated the divergence times (T0 and T1), the effective population sizes of each population (CAA/WG, NEG/Sv and SEG) and the migration rates among each pair of populations. The expected likelihood under the model was obtained using 1,000,000 coalescent simulations and 100 Brent optimization cycles. We ran 100 independent simulations, using different starting values, and selected the parameter estimates of the run attaining the maximum likelihood estimate. To evaluate whether the model could accurately reproduce the observed data, we visually compared the expected and observed marginal one-dimensional SFS, as well as the three two-dimensional SFS pairs.

#### Local adaptation

##### PBS

To test for signatures of positive selection in the three geographically distinct inferred populations, we used population branch statistics (PBS) (Yi et al. 2010) and PCAdapt (Yi et al. 2010; Luu et al. 2017; Privé et al. 2020).

PBS identifies alleles that have undergone significant frequency shifts in a target population, compared to two other populations. The approach identifies highly differentiated genomic regions in each population. We computed population branch statistics and $F_{ST}$ in ANGSD for each population. We used the site allele frequency likelihoods generated for each population, and the 2D-SFS generated as described in the fixation statistics section including both the unfolded 2D-SFS with the finless porpoise as the ancestral state, or the folded 2D-SFS. We computed $F_{ST}$ using the Hudson estimator and PBS using the realSFS function in ANGSD in windows of 50 kb with a 10 kb slide. We adopted an outlier-based approach, and selected the most extreme values of the empirical distribution—specifically >99.95th or >99.99th percentile. For each identified outlier region, we identified overlapping genes and analyzed haplotype patterns of genes of interest across samples.

##### PCAdapt

We also performed a genome scan using the package PCAdapt (Luu et al. 2017; Privé et al. 2020). PCAdapt identified SNPs whose allele frequency differences among populations, characterised by PCA, are unexpectedly large, as outlier candidates for local adaptation. It has the advantage of not requiring grouping of individuals a priori into populations, and is robust when there are admixed individuals (Luu et al. 2017). We first evaluate the best number of principal components (PCs) to consider by examining the percentage of variance explained on a scree plot, and which PCs ascertain putative population structure on the PCA. PCAdapt models the correction between individual allele count and the first $K$ PCs. It evaluates the association strength using the robust Mahalanobis distance, and detects outliers by converting these distances into P-values based on a chi-squared distribution with $K$ degrees of freedom. We ran the analyses with all SNPs and after SNP thinning to account for linkage disequilibrium using a window size of 100 SNPs and an r2 threshold of 0.1.

We identified outlier SNPs (α = 0.1) in each accounting for a false-discovery rate of 10% by calculating q-values with the qvalue function in R package “qvalue” (Storey et al. 2020). We also apply the Bonferroni or Benjamini– Hochberg correction, as a comparison.

#### Diversity statistics

##### Heterozygosity

We estimated heterozygosity based on the 1DSFS generated by ANGSD, as the product of the number of variants in the second column of the est.ml file, which corresponds to the heterozygotes, divided by the total number of sites. We tested whether mean heterozygosity was significantly different among the three populations using a Kruskall-Wallis tests.

##### ROH

Runs of homozygosity (ROH) arise when an offspring receives two copies of the same ancestral haplotype from both parents. ROH can be broken up by recombination in subsequent generations. The ROH length distribution across the genome can inform on demographic history and levels of inbreeding (Ceballos et al. 2018). We used both the window-based approach implemented in Plink v. 1.90b6.21 (Purcell et al. 2007) and the MCMC approach implemented in bcftools v. 1.15 to identify ROH.

We used Plink to transform the vcf file to the Plink ped and map format. We generated a vcf file with the same filtering options as described above with and without a filter of MAF 0.05. We estimated ROH with no MAF filter for all the parameters combination but made a comparison including a filter for MAF 0.05.

In Plink, we set the sliding window size to 300 kb (--homozyg-kb 300) with a minimum of 50 SNPs (--homozyg-snp 50) at a minimum density of 1 SNP per 50 kb (--homozyg-density 50) required to call a ROH. We varied the minimum density to 1 SNP per 40 kb and 1 SNP per 60 kb - results for 40 kb are presented. To take genotyping errors into account, we allowed up to 3 heterozygous sites per 300 kb window (--homozyg-window-het 3) within ROHs (Ceballos et al. 2008). We ran the analysis on the dataset without a MAF filter and with a MAF of 0.05. We also increased this number to 5 heterozygous sites and obtained similar results, in terms of comparable differences when comparing the three inferred narwhal populations, albeit a larger part of the genome was found in ROH. We set the length between two SNPS to be 1000 kb for them to be considered in two different ROH (--homozyg-gap 1000). We also tried varying this length to 500 and 5000 kb. We tried different values (5, 10, 15, 20) for the number of missing calls per window (--homozy-windows-missing) and evaluated when the number of usable windows plateaued, which was 10.

We used bcftools to parse the vcf file per population. We also calculated allele frequencies using bcftools +fill-tags and included it to the vcf file per population and over all individuals. We identified ROH using bcftools with the option G30 specifying that allele frequencies should be estimated from each of the three inferred populations (which included 7 to 9 individuals each), or from the total of 24 individuals when working with the entire vcf file.

For both programs, we calculated mean ROH length and maximum ROH length per individual (unit Mbp or kb). We estimated FROH as the fraction of the autosomal scaffolds within ROH per individual, after removing filtered regions such as repeats (total unmasked and unfiltered genome size) grouped by population. We plotted mean ROH length, maximum ROH length and mean FROH per population in R using the package ggplot2.

The length of the homozygote tracts decline as a function of recombination rate and time, and the parts of the genome in ROH of different lengths can provide information about past changes in effective population size. Therefore ROH – due to an ancestral bottleneck or low effective population size – are expected to be shorter than ROH due to recent inbreeding, as the former would have been broken up by recombination. We can calculate the age of the ROH using the formula from Thompson (2013): g = 100/2*ROHlength*r, where g is the age of the ROH in generations and r the recombination rate in cM. We used the recombination rate of humans of 1.133 cM/Mb (Dumont and Payseur 2008) (Dumont and Payseur 2008) and we are mindful that recombination rate may be different in narwhals. We converted it to years using the generation time of 30 years for narwhals (Garde et al. 2007).

We statistically compared the mean ROH length and maximum ROH length, FROH, and the length of ROH in different size class among the three populations using a linear model (anova) or Kruskall-Wallis test depending on normality and homogeneity of variances.

#### Mutation load

We filtered out genotypes with a depth of coverage <5× as well as heterozygous genotypes, where the minor allele was not supported by at least 2 reads. We annotated the vcf file with snpeff version 5.2c (Cingolani et al., 2012) and the available annotation for the narwhal, which was already in the snpeff database. We identified the ancestral state of each allele by creating a consensus sequence using the beluga and finless porpoise genomic data.

We estimated coverage using the doDepth function in ANGSD and downsampled the finless porpoise genome to 17× so that it matches the coverage of the beluga genome. We merged the two genomes using samtools and inferred the consensus sequence by selecting the most common base using the doFasta 2 option together with doCounts 1 in ANGSD. We used Plink, vcftool, bedtools v. 2.30.0, and R following (Humble et al. 2023) pipeline to incorporate the ancestral state and polarise the alleles in the vcf file. Note that we removed the sites with warnings after running snpeff, and restricted the analysis to the sites where the reference is ancestral and the alternate is derived based on our outgroup.
(roughly 94% of the sites). We also run the analysis without the filter of the heterozygous sites, where the minor allele was not supported by at least 2 reads, and with a MAF of 0.01.

Following Rasmussen et al. (2023) we counted the total number of derived alleles categorised as high impact and moderate impact (total allele count) for each of the 24 individuals, as well as separated by zygosity: in homozygous state (realized allele count) and in heterozygous state (masked allele count). Moderate-impact variants are nondisruptive variants that might change protein effectiveness (i.e. missense variants). High-impact variants are assumed to disrupt the protein, and can cause protein truncation, loss of function (LoF) or initiate nonsense-mediated decay (i.e. stop codons, splice donor variant and splice acceptor and start codon loss).

To take differences in coverage into account, we normalized the derived allele count by the derived allele count per sample in synonymous positions, to estimate the total load, realized load and the masked load. To compare the three inferred populations, we calculated the averages of relative counts of genetic load (for high- and moderate-impact variants) per population. We statistically compared total, masked, and realised load for high and moderate impact mutations among the three populations using a linear model (anova) after checking for normality and homogeneity of variances. Our data satisfied normality and homogeneity of variance.
